# Indazolone-Based Molecular Glue Degraders as a Tunable Platform for Reprogramming Cereblon Substrate Specificity

**DOI:** 10.64898/2026.03.16.712139

**Authors:** Hui-Jun Nie, Jiamin Wang, Hao Xu, Yin-Jue Zhou, Guang-Liang Yin, Xiaomim Xu, Gaoya Xu, Beijing Chen, Xian Li, Xiaobei Hu, Yubo Zhou, Jia Li, Xiao-Hua Chen

## Abstract

Molecular glue degraders (MGDs) represent a transformative modality in drug discovery, with the cereblon (CRBN)–MGD axis offering a strategic gateway to systematically address the historically undruggable proteome. However, the limited chemical diversity of existing CRBN-mediated MGDs has constrained the full exploration of the degradable landscape. Here, we introduce an innovative indazolone-based platform that extends beyond canonical isoindolinone-glutarimide scaffolds, significantly expanding the accessible chemical space for CRBN modulation. Guided by mechanistic insights into CRBN-MGD complex conformational plasticity, we rationally designed indazolone architectures as novel CRBN ligands, developing diverse MGDs with potent ligase binding and exceptional substrate programmability. This platform enables fine-tuned control over neo-substrate recognition, supporting diverse degradation profiles ranging from the broad targeting of critical proteins (IKZF1/3, ZFP91, and LIMD1) to the exquisitely selective degradation of CK1α and IKZF2. Beyond expanding the chemical landscape for intractable targets, our indazolone-based MGDs exhibit favorable pharmacokinetic properties, offering robust promise for therapeutic development. Collectively, the remarkable tunability and exceptional neo-substrate programmability of the indazolone-based platform provide a highly transformative blueprint for next-generation MGD discovery, enabling systematic exploration of the vast CRBN-accessible proteome. Furthermore, as superior CRBN ligands, this platform holds immense potential for PROTAC discovery to systematically address previously intractable targets.

## Introduction

Targeted protein degradation (TPD) has emerged as a revolutionary paradigm in drug discovery, offering a transformative modality to eliminate previously undruggable proteins beyond the reach of conventional inhibition.^1–5^ Unlike occupancy-based inhibitors, TPD harnesses the cell’s endogenous proteostasis machinery to selectively degrade disease-associated proteins, thereby providing innovative therapeutic strategies for challenging targets pivotal in complex diseases.^1^ Within the TPD realm, molecular glue degraders (MGDs) have garnered exceptional interest, primarily due to their distinct pharmacological advantages over heterobifunctional proteolysis-targeting chimeras (PROTACs), including lower molecular weight and superior drug-likeness.^1–4^ Mechanistically, MGDs operate by orchestrating a topographical remodeling of E3 ligase surfaces, inducing neomorphic protein-protein interaction (PPI) interfaces that enable the recruitment of non-native substrates (neo-substrates).^6–10^ Specifically, neo-substrate recognition is exquisitely sensitive to molecular geometry; even subtle structural perturbations to an MGD structure can profoundly reprogram the E3 ligase’ s substrate preference (Figure 1a).^8–10^ Consequently, this unique capacity to redirect E3 specificity holds profound implications for drug discovery, ultimately offering transformative potential for systematically expanding the druggable proteome.^10–17^

**Figure 1.**
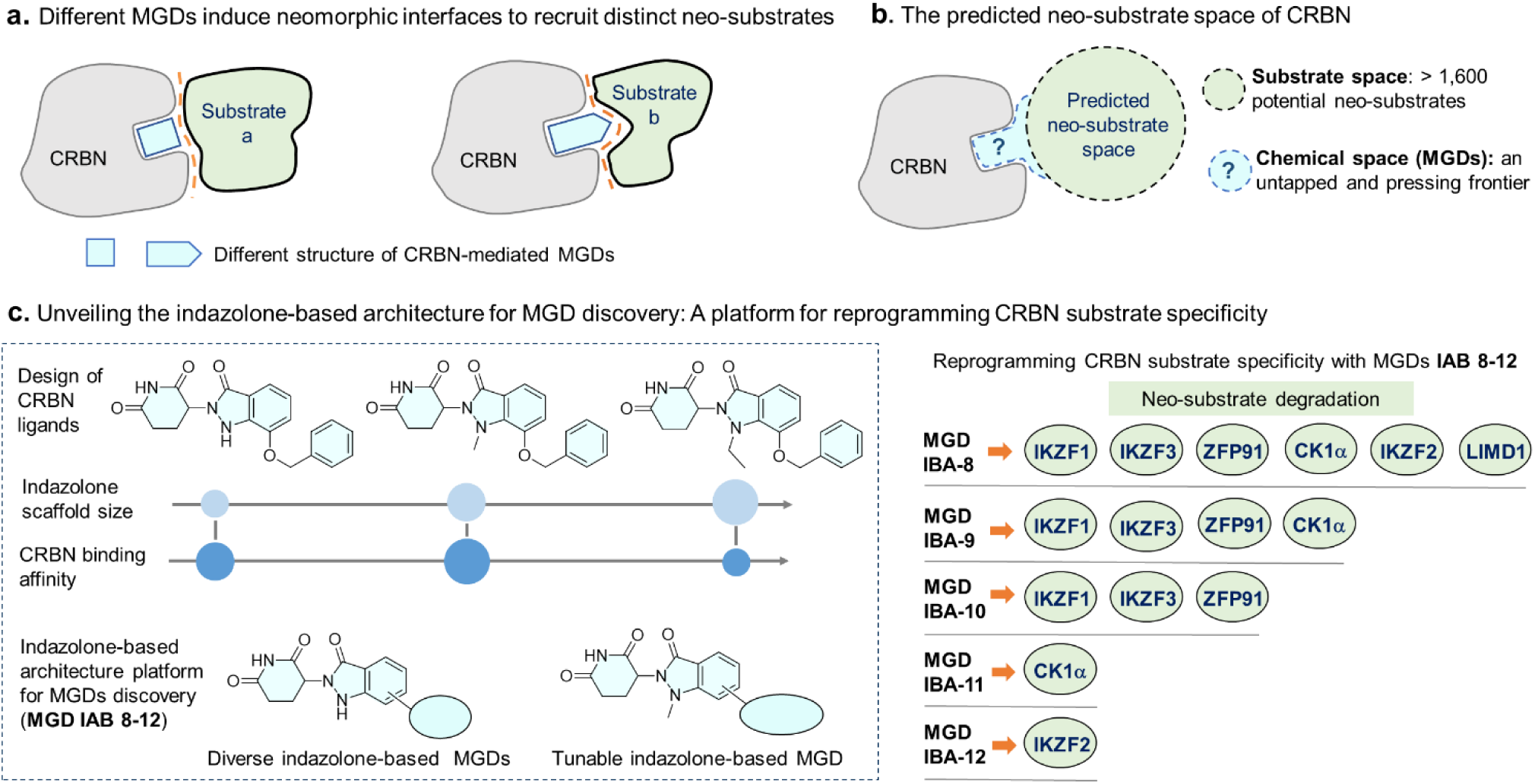
Mechanisms of CRBN neo-substrate recruitment and design of an indazolone-based MGD platform for reprogramming substrate specificity. **a)** MGD-induced neomorphic interfaces enabling selective substrate recruitment, and the predicted CRBN-accessible degradome. **b)** The predicted neo-substrate space of CRBN. **c)** Design of indazolone-based architectures as novel CRBN binders, and the development of structurally diverse MGDs to reprogram CRBN substrate specificity, thereby expanding the MGD-accessible degradome.

The clinical validation of this reprogramming principle is exemplified by the immunomodulatory imide drugs (IMiDs)—including thalidomide, lenalidomide (**LENA**), and pomalidomide.^10, 18–20^ These landmark therapeutics fundamentally demonstrated that small molecules can redirect the E3 ligase cereblon (CRBN) to eliminate critical oncogenic drivers, such as IKZF1/3^21–24^ and CK1α^25–29^ , thereby establishing CRBN as the preeminent hotspot for degrader discovery.^10–12, 20, 30^ More importantly, this clinical success has not only marked a new era for targeting previously intractable proteins but has also catalyzed an intensive pursuit of CRBN next-generation MGDs targeting an expanding array of disease-associated proteins, including GSPT1,^31–34^ IKZF2,^27, 35–38^ WIZ, ^39, 40^ Wee1,^41^ VAV1,^42^ and NEK7.^43,44^ Furthermore, beyond these established successes, recent systematic mining of the CRBN-accessible substrate space has unveiled an extraordinary, yet largely unexplored, proteomic breadth (Figure 1b).^11, 12, 16, 17^ Genomic and structural profiling suggests that the human proteome harbors a vast repertoire of over 1,600 potential neo-substrates—spanning numerous essential transcription factors and kinases—that are theoretically susceptible to CRBN-mediated recruitment.^12^ These targets can be engaged through a spectrum of recognition mechanisms, ranging from the recognition of cryptic structural motifs, such as helical G-loops, to the recruitment of substrates via non-canonical degrons. Therefore, the CRBN-MGD axis represents an extensive resource for therapeutic intervention, serving as a strategic gateway to systematically access the vast, historically undruggable proteome. ^10–12, 16, 17, 20, 45^

However, despite this expansive theoretical degradome, a significant gap persists between the vast landscape of CRBN-accessible targets and the current chemical repertoire.^10–12^ To date, the development of clinical and emerging CRBN-based MGDs has remained predominantly dependent upon the canonical isoindolinone glutarimide scaffold.^10, 46, 47^ Given the stringent structural requirements for neo-substrate recognition and specificity, this over-reliance on isoindolinone glutarimide architecture may inadvertently restrict the degradable landscape to a narrow subset of the proteome,^11, 48^ leaving vast regions of the target space unreachable.^15^ Furthermore, while alternative CRBN-binding ligands, such as phenyl,^49^ benzamide,^50^ and anilino glutarimides,^51^ as well as other aryl glutarimide derivatives,^26, 52, 53^ have been successfully employed in heterobifunctional PROTAC development, their potential to function as monovalent MGD scaffolds for reprogramming substrate specificity remains largely underexplored.^54, 55^ Consequently, the development of novel CRBN-engaging scaffolds beyond the traditional isoindolinone glutarimide paradigm represents a strategic necessity for the discovery of next-generation MGDs.^15, 46, 47, 50, 51^ Such unprecedented architecture-driven exploration is poised to facilitate the discovery of novel MGDs, offering a profound opportunity to systematically expand the boundaries of the druggable proteome.^10, 56^

Herein, we introduce an innovative indazolone-based discovery platform that effectively expands the chemical space for CRBN-mediated MGDs (Figure 1c), successfully transcending the architectural constraints of canonical isoindolinone glutarimide scaffolds. Guided by mechanistic insights into the conformational plasticity of the CRBN-MGD complex, we rationally designed diverse innovative indazolone architectures. This platform demonstrates unprecedented programmability over CRBN substrate specificity. Leveraging the intrinsic tunability of the indazolone architecture, we achieved fine-tuned control over neosubstrate recognition, enabling diverse degradation profiles —ranging from broad-spectrum degradation of critical proteins (IKZF1/3, ZFP91, LIMD1, CK1α, and IKZF2) to exquisitely selective target degradation. Furthermore, our findings validate that this distinct chemotype not only maintains highly effective CRBN binding but also exhibits favorable pharmacokinetic properties, making it highly suitable for preclinical translation. Collectively, by decoupling MGD discovery from traditional structural constraints, this versatile indazolone-based platform—serving as a highly efficient CRBN ligand—provides a blueprint to catalyze next-generation MGD and PROTAC development, offering a strategic gateway to systematically explore the vast CRBN-accessible target space and address intractable human diseases.

## Results and discussion

### Shifted Conformation-Inspired Design of Indazolone-Based Architectures as CRBN Binders

Structural insights from the recently reported CRBN-LENA-CK1α complex reveal a fascinating conformational adaptation within the CRBN E3 ligase (Figure 2a). Upon recruitment of the neo-substrate CK1α, LENA undergoes a significant 2.5 Å positional shift toward Glu377.^57^ This pivotal observation uncovers a crucial structural secret: the IMiD-binding pocket is not a restrictive, rigid cavity, but rather harbors pre-existing permissive space that allows for ligand movement. Such inherent plasticity is essential for accommodating the subtle structural rearrangements required to facilitate optimal protein-protein interactions.

**Figure 2.**
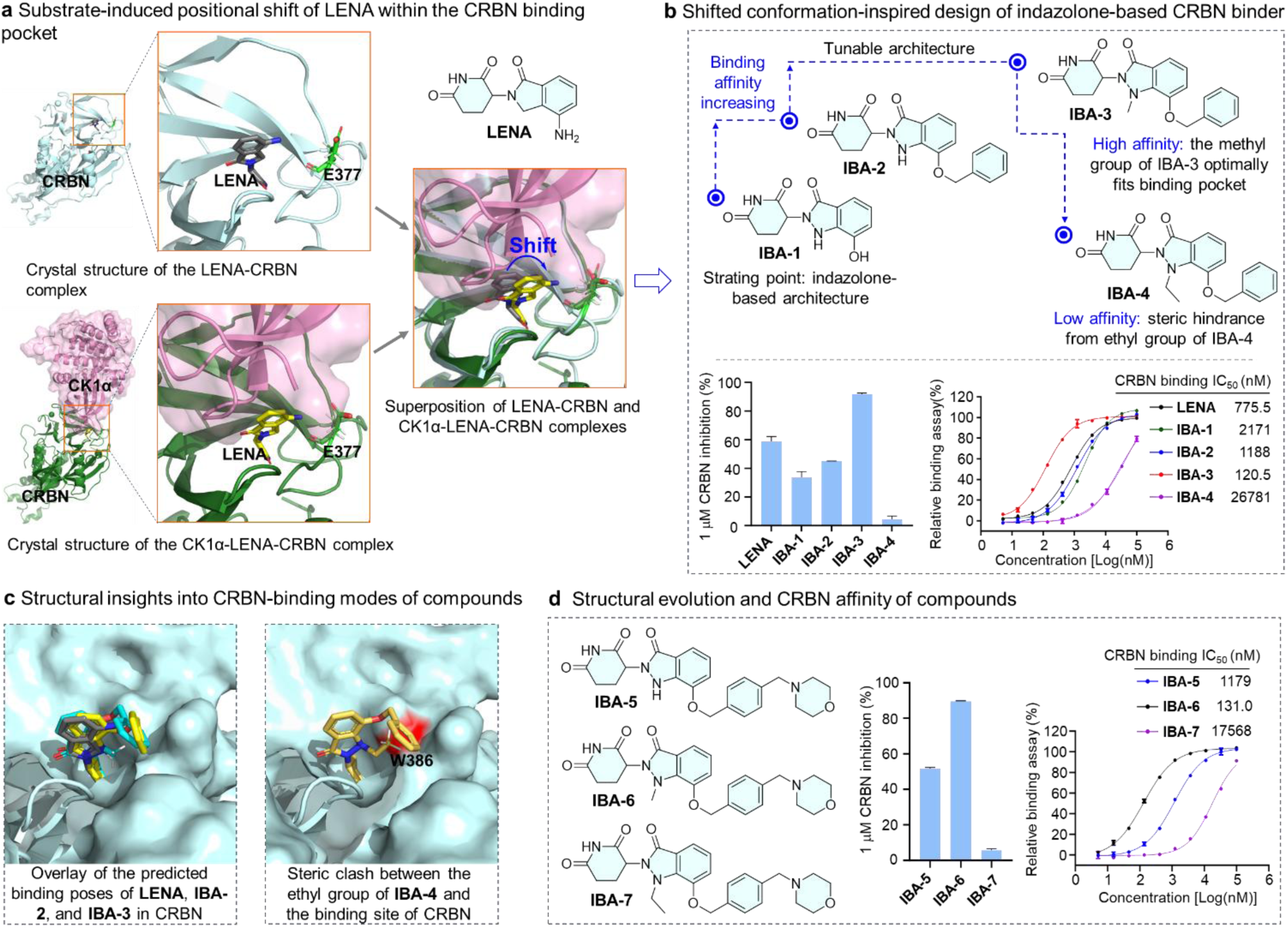
Establishment of a tunable indazolone-based architecture platform for MGDs discovery. **a)** Overlay of the crystal structures of the LENA-CRBN complex (PDB ID: 4TZ4) and the CK1α-LENA-CRBN complex (PDB ID: 5FQD), upon recruitment of the neo-substrate CK1α, LENA undergoes a significant 2.5 Å positional shift toward Glu377. **b)** Chemical structures of designed tunable indazolone derivatives (**IBA-1** to **IBA-4**) with various N-substituents. Evaluation of the relative CRBN binding affinities of lenalidomide and compounds **IBA-1** to **IBA-4** using a TR-FRET assay, displaying the relative inhibition at 1 μM, and IC_50_ values of compounds. Data represent mean ± SD (n = 2 biologically independent samples). **c)** Molecular docking poses of the synthesized compounds. The left panel shows the overlay of **LENA** (gray), **IBA-2** (yellow), and **IBA-3** (cyan) within the CRBN pocket (PDB ID: 4TZ4). The right panel displays the predicted position of the N-ethyl group of **IBA-4** within the CRBN binding site. **d)** Chemical structures of compounds **IBA-5** to **IBA-7**. Assessment of their relative CRBN binding affinities via TR-FRET analysis, detailing the percent inhibition at 1 μM and the IC_50_ values of compounds. Data represent mean ± SD (n = 2 biologically independent samples).

Inspired by the existence of this navigable spatial volume, and leveraging our expertise in light-induced primary amine and o-nitrobenzyl alcohol cyclization (PANAC) chemistry,^58–64^ we envisioned that the indazolone-based architecture—a product of PANAC photoclick chemistry—could serve as a promising structural platform for designing next-generation CRBN binders (Figure 2b). Central to our design is the intrinsic N-substitutability of the indazolone framework, which provides a highly tunable structural template. Unlike the rigid classic isoindolinone scaffold, this optimizable platform allows for the precise installation of functional groups to strategically occupy and exploit the movable space identified by the 2.5 Å shift (Figure 2a). By fine-tuning these N-substituents, we aim to move beyond the traditional isoindolinone paradigm, exploring untapped chemical space to achieve superior geometric complementarity and enhanced neo-substrate recruitment (Figure 1b).

Driven by this hypothesis, we synthesized compounds **IBA-1** and **IBA-2** (Figure 2b). Then, we employed a TR-FRET CRBN binding assay to assess the relative binding affinities. To our delight, when evaluated for CRBN inhibition at a concentration of 1 μM, these indazolone-based architectures exhibited binding affinities comparable to that of **LENA**. This initial result strongly indicated that the indazolone scaffold could serve as a promising CRBN ligand. To further explore this, we synthesized compound **IBA-3** by introducing a methyl substituent at the N-position of the indazolone core. Remarkably, this modification led to a substantial enhancement in binding capacity (91% inhibition for **IBA-3** vs. 59% for **LENA**) (Figure 2b). Subsequent IC50 determinations revealed that **IBA-3** achieved a binding affinity of 120 nM, representing a more than 6-fold improvement over **LENA** (120 nM vs. 776 nM). Furthermore, to probe the spatial limits, we introduced a bulkier ethyl group at the nitrogen atom (**IBA-4**), which resulted in an almost complete loss of CRBN binding affinity (IC50 > 26 μM). To rationalize these divergent structure-activity relationships, we performed molecular docking analyses of these compounds with CRBN (Figure 2c). The predicted binding poses revealed that these indazolone derivatives share a binding mode highly similar to that of **LENA** observed in the crystal structure, successfully overlaying within the pocket. Together, the experimental data and computational predictions suggest that the N-methyl group of **IBA-3** optimally exploits the pre-existing permissive space within the CRBN binding pocket, thereby conferring high binding affinity. Conversely, the unsubstituted indazolone scaffolds (**IBA-1** and **IBA-2**) display comparable binding affinities to LENA due to their similar steric bulk to the classic isoindolinone scaffold. However, the introduction of the ethyl group in **IBA-4** induces severe steric clashes with Trp386 (Figure 2c, 2d), profoundly abrogating its ability to bind CRBN.

To further investigate whether increased structural complexity would compromise the CRBN-binding affinity of these indazolone-based architectures, we synthesized compounds **IBA-5** to **IBA-7** by incorporating a morpholinomethyl group onto the phenyl ring (Figure 2d). As anticipated, **IBA-5** (the unsubstituted derivative) exhibited a binding profile comparable to its corresponding analogue, **IBA-2** (Figure 2b and Figure 2c). Meanwhile, **IBA-6**, which bears an N-methyl substituent on the indazolone core, demonstrated robustly enhanced CRBN-binding activity, analogous to **IBA-3**. Conversely, the introduction of an N-ethyl group in **IBA-7** once again resulted in poor binding affinity, mirroring the behavior of **IBA-4**. Compellingly, these findings validate that our rationally designed indazolone-glutarimide architecture can serve as a robust next-generation CRBN binder. More importantly, the strategic introduction of a methyl group at the nitrogen atom yields a highly tunable architecture, offering a broader repertoire of choices for the design of CRBN-targeting MGDs.

This binding affinity trend was also reflected in the antiproliferative activity. The series showed potent activity in MM.1S cells, where **IBA-2**, **IBA-3**, **IBA-5**, and **IBA-6** inhibited cell proliferation with IC50 values of 351.3, 160.1, 150.8, and 30.4 nM (Table S1), respectively. Notably, **IBA-6** exhibited superior potency compared to the positive control lenalidomide (**LENA**, IC50 = 91.7 nM), whereas the N-ethyl analogues **IBA-4** and **IBA-7** were inactive (Table S1). Consistent with the CRBN-binding data, the N-methyl analogues **IBA-3** and **IBA-6** were more potent than their unsubstituted counterparts. In addition, Rat liver microsomal stability further revealed the favorable metabolic profile of these architectures (Table S2). Moreover, all compounds from **IBA-2** to **IBA-6** exhibited negligible hERG inhibition (IC50 > 40 μM), suggesting a favorable cardiac safety profile. Furthermore, to assess the chiral stability of our architecture, pure enantiomers of **IBA-2** were obtained by chiral separation and examined in phosphate-buffered saline (PBS) (pH 7.4) and cell culture medium. **IBA-2** underwent nearly complete racemization within 4 h in PBS and within 1 h in cell culture medium. This feature of the indazolone glutarimide architecture is highly consistent with that of canonical IMiD scaffolds (e.g., thalidomide and pomalidomide), which are known to undergo rapid racemization at the C3 chiral center of the glutarimide ring in aqueous and biologically relevant media.^23, 65^ Accordingly, all compounds in this study were synthesized and evaluated as racemates.

Taken together, these results demonstrate that increasing the molecular complexity based on either the unsubstituted or N-methylated indazolone architectures does not compromise productive CRBN engagement, while maintaining favorable cellular activity and metabolic stability. This feature is of paramount importance, as it successfully unlocks novel chemical space for the subsequent development of structurally diverse MGDs built upon these two architectures.

### Discovery of Indazolone-Based MGD Targeting Diverse Neo-Substrates

Mechanistically, CRBN-mediated MGDs function by anchoring to the CRBN E3 ligase, thereby forming a neomorphic PPI interface that recruits neo-substrates harboring recognition motifs such as canonical G-loops or G-loop–like features. To guide the design of MGDs, recent structural and mechanistic studies in the field have conceptualized a prevailing model that partitions the MGD into a conserved core and a tunable extended region (Figure 3a)^10^. In this established paradigm, the MGD core must preserve essential interactions within the IMiD-binding pocket while maintaining a compatible geometric shape to accommodate the incoming degrons. Beyond this core, the extended region serves as the primary determinant of the degradation profile. Structural modifications within this extended space—ranging from single-atom substitutions to more complex molecular expansions—synergistically dictate target selectivity and stabilize the neomorphic PPI interface, thereby simultaneously modulating substrate preference and overall binding potency.^22, 23^ Building on this field-wide framework, we recognized that our indazolone-based architectures inherently satisfy the restrictive requirements of the CRBN-binding core, while offering versatile substitution vectors to strategically access and tune this critical extended region. Driven by this rationale, we designed and synthesized a variety of structurally more complex MGDs based on the unsubstituted or N-methylated indazolone architectures.

**Figure 3.**
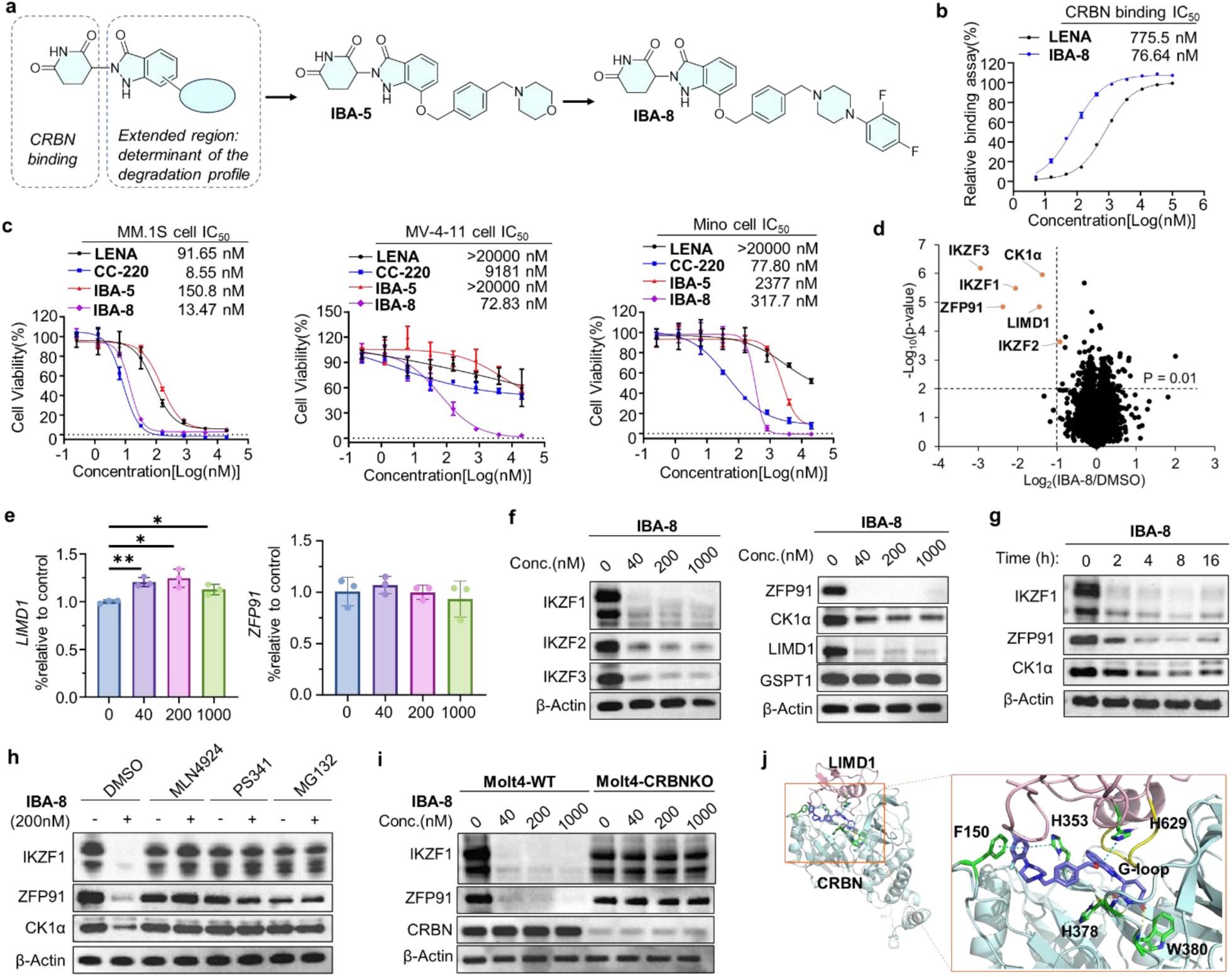
Discovery and degradation landscape of the indazolone-based MGD IBA-8. **a)** Chemical structures of **IBA-8**. **b)** Relative CRBN binding affinities of lenalidomide and **IBA-8** determined by TR-FRET. Data represent mean ± SD (n = 2 independent replicates). **c)** Cell viability assay of **IBA-8** in MM.1S, Mino, and MV-4-11 cells. Data are presented as mean ±SD, (n = 3 independent replicates). **d)** whole-cell proteomic profiling in Mino cells treated with **IBA-8** at 200 nM for 4 h. Significant enrichment: Log_2_ (**IBA-8**/DMSO) < -1 and p-value < 0.01, n = 3. **e)** Quantitative PCR analysis of mRNA expression of *LIMD1* and *ZFP91* in Mino cells treated with DMSO and **IBA-8** (40, 200, 1000 nM) for 8 h. P values were calculated using two-tailed t-test; *p<0.05, **p<0.01. **f)** WB analysis of the concentration-dependent degradation of IKZF1, IKZF2, IKZF3, ZFP91, LIMD1, CK1α, and GSPT1 in Mino cells treated with the various concentrations of **IBA-8** for 8 h. WB, Western Blotting. **g)** WB analysis of the time-dependent degradation of IKZF1, CK1α, and ZFP91 in Mino cells treated with 200 nM **IBA-8** for various durations. **h)** Mino cells were pretreated with DMSO, MLN4924 (0.5 μM), PS341 (1 μM), or MG132 (10 μM) for 2 h, followed by treatment with 200 nM **IBA-8** for 3 h. **i)** WB analysis of IKZF1 and ZFP91 in wild type Molt4 cells and CRBN^⁻/⁻^ Molt4 cells treated with **IBA-8**. **j)** Docking model of the CRBN (cyan): **IBA-8** (light blue): LIMD1 (light pink) ternary complex, **IBA-8** was docked into the CRBN crystal structure (PDB ID: 4TZ4), LIMD1 with β-hairpin G-loop predictions (yellow).

Having established the indazolone–glutarimide architecture as a robust CRBN-binding architecture with tunable substitution vectors, we next sought to determine whether this newly developed platform could be leveraged to induce neo-substrate recruitment and targeted protein degradation. To this end, we designed compound **IBA-8** by extending the **IBA-5** sturcture through replacement of the morpholine moiety with a piperazine linker and incorporation of a substituted aromatic group (Figure 3a). This structural elaboration was intended to probe the accessible interaction space surrounding the CRBN pocket and evaluate whether the indazolone architecture could function as a MGD capable of engaging diverse neo-substrates.

To evaluate its ability to engage CRBN, we measured the binding affinity using a TR-FRET assay. **IBA-8** exhibited potent binding activity with an IC50 of 76.64 nM (Figure 3b), demonstrating an approximately 10-fold enhancement in affinity compared with lenalidomide (IC50 = 775.5 nM). In antiproliferative assays, **IBA-8** showed markedly improved cellular activity over **IBA-5** and lenalidomide, and displayed greater potency than CC-220^22^ in MV-4-11 cells (Figure 3c). **IBA-8** inhibited MM.1S, MV-4-11, and Mino cells with IC50 values of 13.47, 72.83, and 317.7 nM, respectively (Figure 3c). These data indicate that structural elaboration of the **IBA-5** architecture to **IBA-8** substantially enhances cellular potency and broadens its antiproliferative profile. To further evaluate its metabolic stability, the microsomal stability of **IBA-8** was assessed in rat liver microsomes (Table S2). **IBA-8** exhibited a half-life (t1/2) of 51.1 min and a low intrinsic clearance (Clint = 0.0678 mL/min/mg), indicating favorable metabolic stability.

Subsequently, to map the degradation landscape and identify specific targets, we performed whole-cell proteomic profiling in Mino cells treated with **IBA-8**. This proteome-wide analysis revealed significant downregulation of several CRBN neosubstrates (Figure3d), including IKZF1/3, IKZF2, CK1α, ZFP91, and LIMD1. qPCR analysis after 8 h of treatment showed that ZFP91 mRNA remained largely unchanged, whereas LIMD1 transcript levels modestly increased at 40 and 200 nM. These results confirm that the reduced protein levels were driven by compound-induced degradation rather than transcriptional suppression (Figure 3e). Based on these findings, the identified substrates were selected for further orthogonal validation. Western blot analysis demonstrated that IBA-8 induced dose-dependent degradation of IKZF1, IKZF3, ZFP91, IKZF2, and LIMD1 after 8 h of treatment in Mino cells, with substantial degradation evident at 40 nM, whereas CK1α exhibited a more modest degradation profile (Figure 3f). By contrast, no degradation of GSPT1 was detected. Time-course studies further revealed rapid and near-complete degradation of IKZF1 and ZFP91 within 2-4 h at 200 nM; consistent with the dose-response data, CK1α degradation proceeded with slower kinetics and reduced efficiency (Figure 3g).

To investigate the mechanism of action, we examined whether **IBA-8**-induced degradation depends on the ubiquitin–proteasome system (UPS). Degradation of IKZF1, ZFP91, CK1α, IKZF2 and LIMD1 was fully rescued by cotreatment with MLN4924 or the proteasome inhibitors PS341 and MG132 (Figure 3h and Figure S1a), indicating a CRL-dependent proteasomal mechanism. CRBN dependence was further confirmed in wild-type Molt4 cells and CRBN-knockout Molt4 cells, where **IBA-8**-induced IKZF1, ZFP91 and LIMD1 degradation was abolished in the absence of CRBN (Figure 3i and Figure S1b). Similarly, CK1α degradation was detected in wild-type but not CRBN-knockout HEK293T cells (Figure S1c), further supporting the CRBN-dependent activity of **IBA-8**.

To gain further insight into the structural basis of **IBA-8**-mediated LIMD1 degradation, we performed molecular docking studies to investigate the putative binding conformations of **IBA-8** within a CRBN–LIMD1 ternary complex model. As shown in Figure 3j, consistent with the canonical binding mode of previously reported CRBN-recruiting molecular glues, the glutarimide ring of **IBA-8** occupies the tritryptophan pocket of CRBN. Notably, the 2, 4-difluorophenyl moiety of I**BA-8** is predicted to form π-π interactions with Phe150 and His353 of CRBN. In addition, the indazolone ring likely engages in a π-π interaction with the side chain of His629 from LIMD1, providing critical interfacial stabilization. The AlphaFold-predicted structure of LIMD1 contains a canonical β-hairpin G-loop degron that exhibits favorable surface complementarity and direct protein-protein contacts at the CRBN–**IBA-8** interface, thereby enabling MGD-dependent recruitment.

Collectively, these results demonstrate that **IBA-8** functions as a potent CRBN-dependent MGD capable of degrading multiple neo-substrates, including IKZF1/3, ZFP91, LIMD1, IKZF2, and CK1α. This multi-target degradation profile highlights the capacity of the indazolone architecture to generate diverse degradation outcomes, providing the first evidence that an indazolone-based MGD can effectively expand the druggable CRBN neo-degradome.

### Structural Refinement of Indazolone-Based MGDs to Reprogram CRBN Substrate Selectivity

Although **IBA-8** demonstrated potent degradation activity toward multiple CRBN neo-substrates, its broad degradation profile also included LIMD1, a well-recognized tumor suppressor protein.^66, 67^ The degradation of such proteins may introduce potential safety concerns, thereby underscoring the vital need to reprogram substrate selectivity within the indazolone architecture. Driven by this rationale, structure-guided optimization of **IBA-8** led to the design of **IBA-9**, in which subtle modifications to the indazolone architecture and side-chain substitution pattern were introduced to remodel the CRBN–substrate interaction interface (Figure 4a). In a TR-FRET assay, **IBA-9** retained strong CRBN engagement with an IC50 of 115.3 nM, representing a ∼7-fold improvement over lenalidomide (IC50 = 775.5 nM) (Figure 4b). Consistent with this robust binding profile, **IBA-9** exhibited potent antiproliferative activity across diverse hematological malignancy cell models, including MM.1S (IC50 = 28.7 nM), Mino (IC50 = 91.2 nM), and MV-4-11 cells (IC50 = 72.5 nM) (Figure 4c).

**Figure 4.**
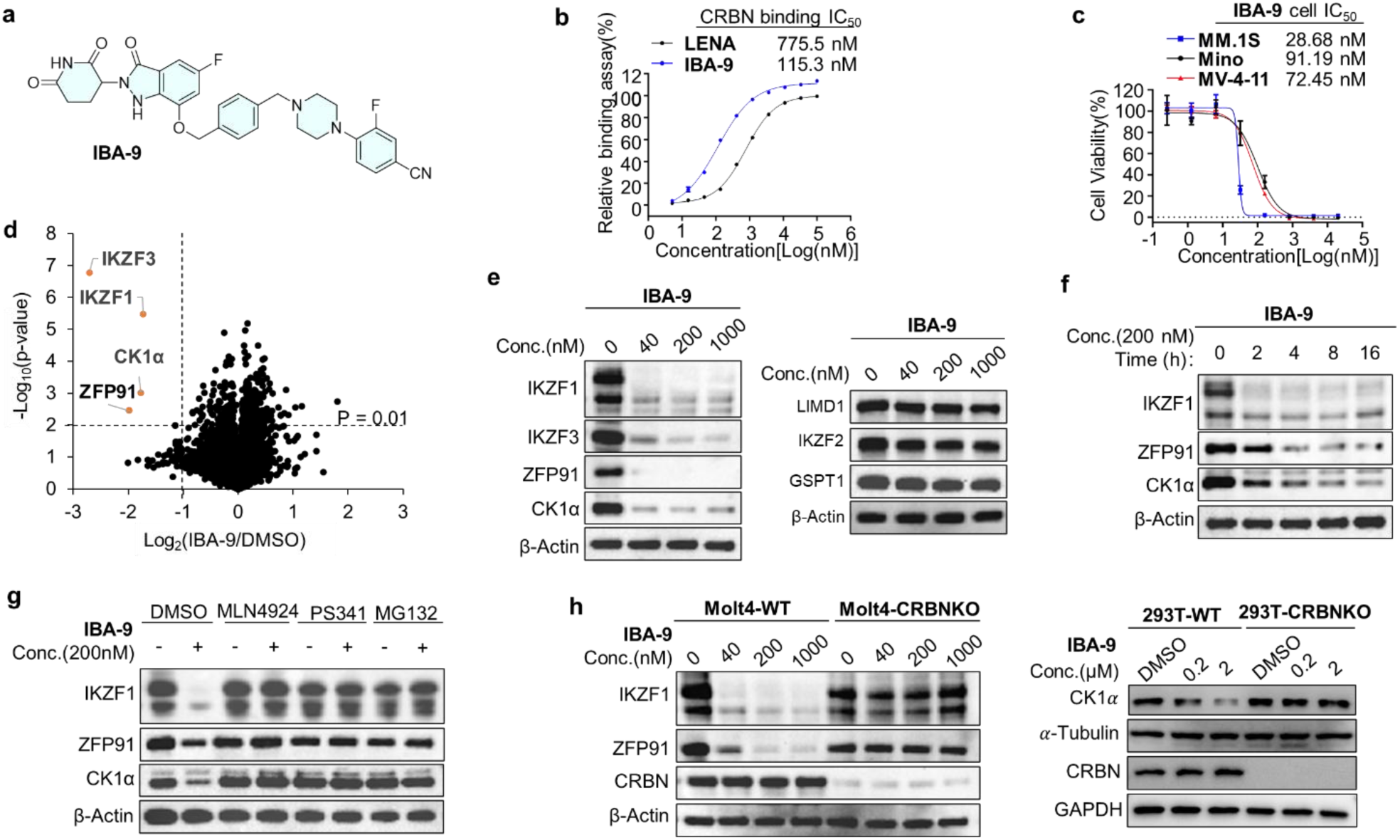
Identification of MGD IBA-9 and validation of its programmable neo-substrate degradation. **a)** Chemical structure of **IBA-9**. **b)** TR-FRET-based evaluation of relative CRBN binding affinities for LENA and **IBA-9**. Date are shown as mean ± SD (n=2 independent replicates). **c)** Anti-proliferative activity of **IBA-9** against MM.1S, Mino, and MV-4-11 cell lines. Date are shown as mean ± SD (n = 3 independent replicates),. **d)** Whole-cell proteomic analysis of Mino cells following treatment with 200 nM **IBA-9** for 4 h. Thresholds for significant protein alterations were set at Log_2_ fold change (**IBA-9**/DMSO) < -1 and p-value < 0.01 (n = 3). **e)** WB analysis of concentration-dependent degradation (IKZF1, IKZF3, ZFP91, CK1α, GSPT1, IKZF2, and LIMD1) in Mino cells exposed to various doses of **IBA-9** for 8 h. **f)** Time-dependent degradation profiles of IKZF1, CK1α, and ZFP91 in Mino cells incubated with 200 nM **IBA-9**. **g)** Rescue of **IBA-9**-induced CK1α degradation by pretreatment with MLN4924 (0.5 μM), PS341 (1 μM), or MG132 (10 μM) for 2 h, followed by treatment with 200 nM **IBA-9** for 3 h in Mino cells. **h)** WB analysis of IKZF1 and ZFP91 in wild type Molt4 cells and CRBN^-/-^ Molt4 cells following treatment with **IBA-9** (left); WB analysis of CK1α in wild-type (WT) and CRBN^-/-^ HEK293T cells following treatment with **IBA-9** (right).

To identify substrates targeted by **IBA-9**, we conducted whole-cell proteomic profiling in Mino cells. Proteome-wide analysis revealed an exquisitely selective degradation signature (Figure 4d), characterized by the significant downregulation of multiple therapeutic neo-substrates, most prominently IKZF1, IKZF3, CK1α, and ZFP91. Crucially, unlike the broad degradation profile of IBA-8, IBA-9 effectively spared the tumor suppressor LIMD1. This striking difference indicates that the introduced structural modifications successfully reshaped substrate selectivity and reprogrammed CRBN-mediated recognition, achieving the desired separation of efficacy from potential liabilities. These targeted proteins were therefore prioritized for subsequent validation and mechanistic studies.

Western blot analysis confirmed that **IBA-9** induced robust, dose-dependent degradation of IKZF1, IKZF3, ZFP91, and CK1α in Mino cells, with clear effects observed at concentrations as low as 40 nM (Figure 4e). In striking contrast, GSPT1, IKZF2, and LIMD1 showed minimal or no degradation even at concentrations up to 1 μM, providing definitive evidence for the improved substrate selectivity of **IBA-9** relative to **IBA-8**. Time-course experiments further revealed rapid degradation kinetics; treatment with 200 nM **IBA-9** led to near-complete depletion of IKZF1 within 2 h (Figure 4f), while CK1α and ZFP91 were substantially degraded by 4 h and nearly completely depleted by 16 h. To clarify the underlying mechanism, we investigated whether this degradation was dependent on the UPS. Co-treatment with MLN4924 or the proteasome inhibitors PS341 and MG-132 fully blocked the target degradation (Figure 4g), confirming a CRL-mediated proteasomal pathway. Consistently, the degradation of IKZF1 and ZFP91 was observed in wild-type Molt4 cells but completely abolished in CRBN-knockout Molt4 cells. Likewise, CK1α degradation was detected in wild-type HEK293T cells but not in CRBN-knockout cells (Figure 4h), reaffirming the stringent CRBN dependence of **IBA-9**.

In addition, **IBA-9** displayed excellent metabolic stability in rat liver microsomes, with an apparent half-life exceeding the assay time window and negligible intrinsic clearance (Table S2). Taken together, these results demonstrate that subtle structural refinement of the indazolone-based MGD can effectively reprogram the CRBN degradome. Compared with the broad substrate spectrum of **IBA-8**, **IBA-9** exhibits a finely tuned degradation profile that eliminates LIMD1 and IKZF2 degradation while maintaining potent activity toward neo-substrates such as IKZF1/3, ZFP91, and CK1a. These findings highlight the programmable nature of the indazolone platform and establish a robust foundation for further tailoring substrate selectivity.

### Rational Tuning of Indazolone-Based MGDs for High-Affinity CRBN Engagement and Refined Selectivity

Encouraged by the ability of **IBA-9** to reshape thedegradation profile relative to **IBA-8**, we next explored whether the indazolone architecture could be further tuned to achieve more precise control over substrate selectivity. Guided by the binding affinity differences between **IBA-2** and **IBA-3**, as well as **IBA-5** and **IBA-6** (Figure 2), we introduced an N-methyl substitution on the indazolone core to enhance CRBN engagement, resulting in the design of **IBA-10** (Figure 5a). This modification was anticipated to further refine the CRBN–substrate interface and modulate the degradation preference. In a TR-FRET binding assay, **IBA-10** exhibited potent CRBN binding (IC50 = 16.6 nM), representing a 47-fold improvement over lenalidomide (IC50 = 775.5 nM) (Figure 5b). We next assessed the antiproliferative activity of **IBA-10** across hematological malignancy cell lines. **IBA-10** demonstrated profound growth inhibition, with IC50 values of 0.86 nM in MM.1S cells and 8.23 nM in Mino cells (Figure 5c), while remaining completely inactive in MV-4-11 cells (> 20 μM). This striking differential sensitivity across cell lines strongly suggests that **IBA-10** may induce a highly cell-context-dependent neo-substrate degradation profile.

**Figure 5.**
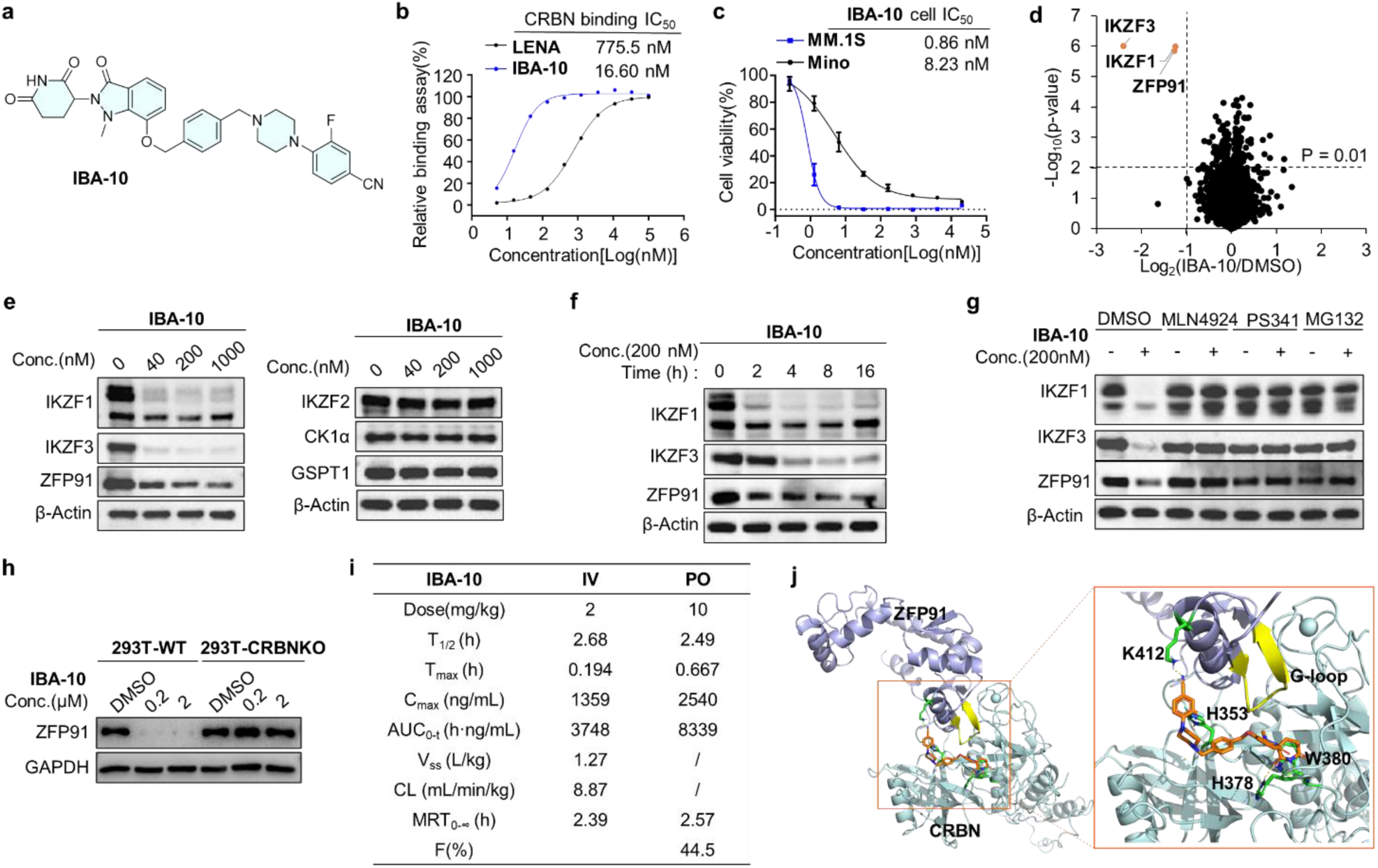
Identification of the indazolone-based MGD IBA-10 and evaluation of its substrate selectivity. **a)** Chemical structure of **IBA-10**. **b)** TR-FRET-based evaluation of relative CRBN binding affinities for LENA and **IBA-10**. Date are shown as mean ± SD (n = 3 independent replicates). **c)** Cell growth inhibition profiles of **IBA-10** in MM.1S and Mino cells. Data represent the mean ± SD (n = 3 independent replicates). **d)** Whole-cell proteomic analysis of Mino cells following treatment with 200 nM **IBA-10** for 4 h. Thresholds for significant protein alterations were set at Log_2_ fold change (**IBA-10**/DMSO) < -1 and p-value < 0.01 (n = 3). **e)** WB analysis illustrating the dose-dependent degradation of multiple CRBN substrates in Mino cells treated with the indicated concentrations of **IBA-10** for 8 h. **f)** Time-dependent degradation profiles of IKZF1, CK1α, and ZFP91 in Mino cells incubated with 200 nM **IBA-10** for various time intervals. **g)** Rescue of **IBA-10**-induced IKZF1/3 and ZFP91 degradation by pretreatment with MLN4924 (0.5 μM), PS341 (1 μM), or MG132 (10 μM) for 2 h, followed by treatment with 200 nM **IBA-10** for 3 h in Mino cells. **h)** WB analysis of CK1α in wild-type and CRBN^-/-^ HEK293T cells treated with **IBA-10**. **i)** Docking binding model of CRBN (cyan): **IBA-10** (yellow): ZFP91 (light blue) complex. **IBA-10** was docked into the CRBN crystal structure (PDB ID: 4TZ4), ZFP91 with β-hairpin G-loop predictions (yellow). **J)** PK Parameters of Compound **IBA-10** in Rats.

To characterize the degradome induced by **IBA-10**, whole-cell proteomic profiling was performed in Mino cells. Proteome-wide analysis revealed a highly selective downregulation of CRBN neo-substrates, primarily targeting IKZF1, IKZF3, and ZFP91 (Figure 5d). Western blot analysis confirmed the robust, dose-dependent degradation of IKZF1 and IKZF3, with substantial depletion observed at 40 nM, whereas the degradation of ZFP91 proceeded with reduced efficiency (Figure 5e). Strikingly, other targets including CK1α, GSPT1, and IKZF2 remained largely unaffected at concentrations up to 1 μM. Time-course experiments further demonstrated rapid degradation kinetics for IKZF1 and IKZF3 within 2–4 h, reaching near-complete depletion by 8 h, while ZFP91 degraded with slower kinetics (Figure 5f).

Mechanistically, pharmacological inhibition with MLN4924, PS341, or MG132 completely abolished target degradation (Figure 5g). Furthermore, genetic deletion of CRBN in HEK293T cells effectively rescued ZFP91 from degradation, confirming that **IBA-10** operates via a CRBN-dependent UPS mechanism (Figure 5h).

To further evaluate the translational potential of **IBA-10**, we investigated its in vitro metabolic stability and in vivo pharmacokinetic (PK) properties. **IBA-10** displayed a favorable metabolic profile in rat liver microsomes, with low intrinsic clearance (Table S2). Subsequent *in vivo* PK profiling in rats (Figure 5i) revealed low systemic clearance (CL = 8.87 mL/min/kg) and a moderate steady-state volume of distribution (Vss = 1.27 L/kg) following intravenous administration. Remarkably, following oral administration at 10 mg/kg, IBA-10 was rapidly absorbed (Tmax = 0.667 h) and achieved a peak plasma concentration (Cmax) of 2540 ng/mL, with a half-life of 2.49 h, translating to an outstanding oral bioavailability (F) of 44.5%. Altogether, these data demonstrate that **IBA-10** possesses a highly favorable metabolic profile and superior drug-likeness, making it an excellent candidate for further preclinical evaluation.

To gain structural insight into ZFP91 recruitment, molecular docking was performed to model the putative CRBN–I**BA-10**–ZFP91 ternary complex. As shown in Figure 5j, the glutarimide moiety of **IBA-10** occupies the canonical tri-tryptophan pocket of CRBN. The 3-fluorobenzonitrile group is predicted to form a π–π interaction with His353 of CRBN, while the cyano group putatively establishes a hydrogen-bond interaction with Lys412 of ZFP91. ZFP91 is predicted to engage the CRBN–**IBA-10** interface through a canonical β-hairpin G-loop. Overall, these results demonstrate that structural optimization of the indazolone architecture can progressively refine the degradation landscape. Compared with the broader activity observed for **IBA-8** and **IBA-9**, **IBA-10** exhibits a more focused degradation profile primarily targeting IKZF1/3 and ZFP91, elegantly illustrating that indazolone-based MGDs enable the stepwise tuning of CRBN substrate selectivity.

### Discovery of an Indazolone-Based Molecular Glue for Highly Selective CK1α Degradation

Having demonstrated that the indazolone architecture allows the progressive modulation of substrate preference, we next sought to determine whether this platform could be further engineered to enable the selective degradation of alternative CRBN neo-substrates. To test this possibility, we designed **IBA-11** by introducing a rigid aryl-substituted N-methylimidazole moiety at the C3 position of the indazolone core, with the aim of reshaping the solvent-exposed region and promoting the recruitment of distinct neo-substrates (Figure 6a). In a TR-FRET assay, **IBA-11** exhibited modest CRBN binding affinity (IC50 = 2.28 μM) (Figure 6b). We next assessed its antiproliferative activity across hematologic malignancy cell lines. Strikingly, in sharp contrast to **IBA-10**, **IBA-11** displayed potent cytotoxicity in MV-4-11 cells (IC50 = 82.11 nM) (Figure 6c), while exhibiting negligible growth inhibition in MM.1S and Mino cells. This inverted phenotypic profile strongly suggested a distinct, target-driven neo-substrate degradation landscape.

**Figure 6.**
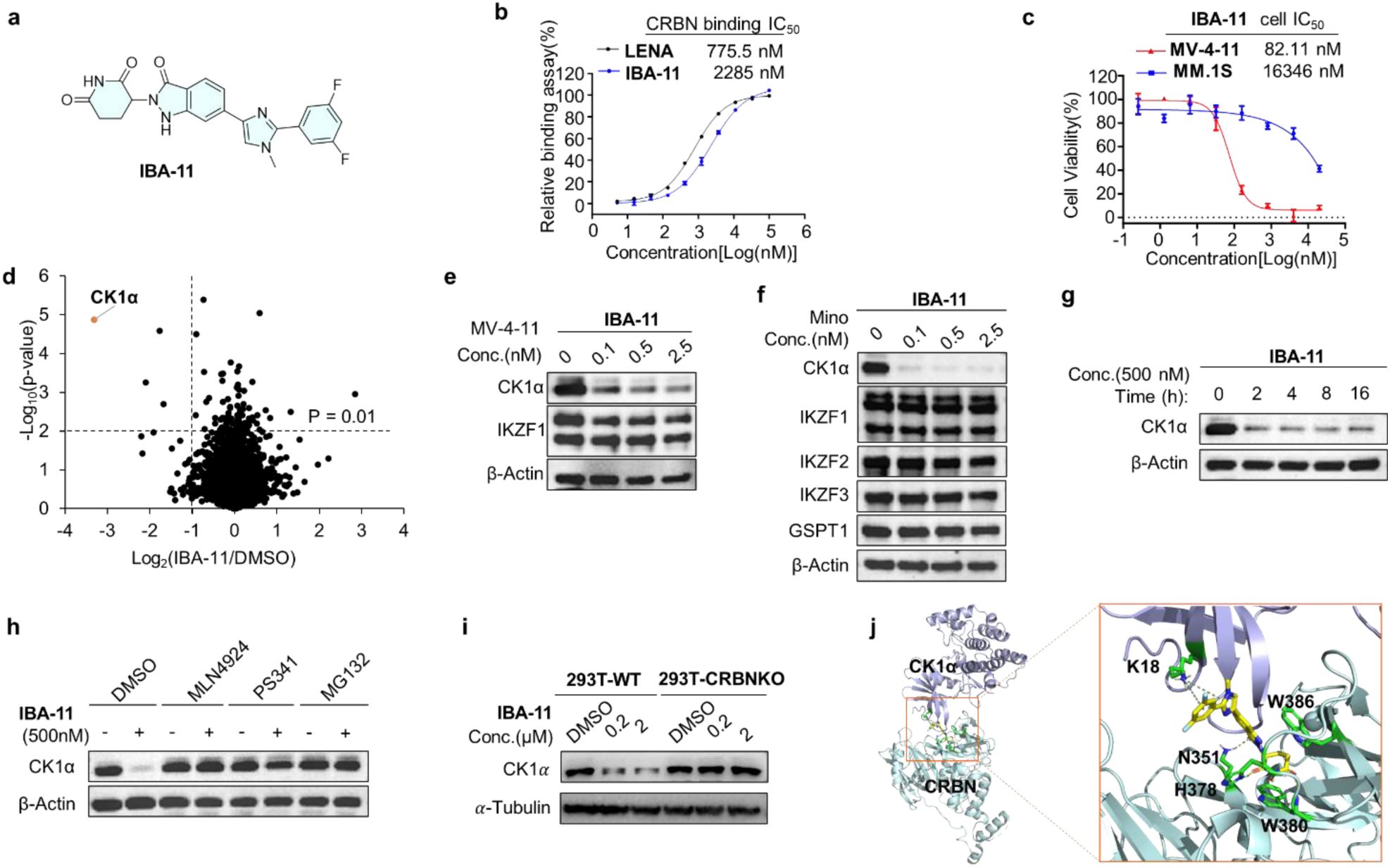
Discovery of the MGD IBA-11 and validation of selective CK1α degradation. **a)** Chemical structure of **IBA-11**. **b)** Assessment of the relative CRBN binding affinities of LENA and **IBA-11** via TR-FRET assay. Data are expressed as mean ± SD (n = 2 independent replicates). **c)** Cell growth inhibition profiles of **IBA-11** in MV-4-11. Data represent the mean ± SD (n = 3 independent replicates). **d)** Whole-cell proteomic analysis of Mino cells following treatment with 200 nM **IBA-11** for 4 h. Thresholds for significant protein alterations were set at Log_2_ fold change (**IBA-11**/DMSO) < -1 and p-value < 0.01 (n = 3). **e)** Western blot analysis of CK1α and IKZF1 in MV-4-11 cells treated with the indicated concentrations of **IBA-11** for 8 h. **f)** The dose-dependent degradation of multiple CRBN substrates in Mino cells treated with the indicated concentrations of **IBA-11** for 8 h. **g)** Time-dependent degradation profiles of CK1α in Mino cells incubated with 500 nM **IBA-11**. **h)** Rescue of **IBA-11**-induced CK1α degradation by pretreatment with MLN4924 (0.5 μM), PS-341 (1 μM), or MG-132 (10 μM) for 2 h, prior to an additional 3h exposure to **IBA-11** (500nM), and subsequently analyzed by WB. **h)** WB analysis of CK1α in wild-type (WT) and CRBN^-/-^ HEK293T cells treated with **IBA-11**. **j)** Docking of **IBA-11** with CRBN and CK1α. Predicted binding mode and interactions of CRBN (cyan): **IBA-11** (yellow): CK1α (light blue) complex (PDB:5FQD).

To identify the underlying targets, whole-cell proteomic profiling was performed in Mino cells to ensure a direct comparison with the degradomes of **IBA-8**, **IBA-9**, and **IBA-10**. The proteome-wide analysis revealed CK1α as the most significantly downregulated protein (Figure 6d). Western blot analysis further confirmed that **IBA-11** induced dose-dependent degradation of CK1α in both MV-4-11 and Mino cells following an 8 h treatment, while IKZF1/2/3 and GSPT1 remained largely unaffected, indicating exceptional orthogonal selectivity (Figure 6e, f). Time-course experiments revealed rapid degradation kinetics, with 500 nM **IBA-11** inducing near-complete CK1α depletion within 2 h (Figure 6g). Mechanistically, this targeted degradation was completely abolished by pretreatment with MLN4924, PS341, or MG132 (Figure 6h), and was notably absent in CRBN-knockout HEK293T cells (Figure 6i), confirming its dependence on the CRBN-mediated UPS. Furthermore, **IBA-11** displayed excellent *in vitro* metabolic stability in rat liver microsomes, featuring a prolonged half-life (t1/2) of 184 min and a correspondingly low intrinsic clearance (Table S2). Crucially, the compound exhibited minimal hERG inhibition (IC50 > 40 μM), indicating a highly favorable cardiac safety profile.

To gain deeper structural insight into the recruitment of CK1α, we performed molecular docking using the established CRBN–CK1α crystal structure (PDB ID: 5FQD) (Figure 6j). Our model suggests that while the glutarimide moiety of IBA-11 anchors into the canonical tri-tryptophan pocket of CRBN, the indazolone core establishes putative hydrogen-bond interactions with N351. Furthermore, Lys18 of CK1α is predicted to form key cation-π interactions with both the fluorophenyl ring and the imidazole moiety of **IBA-11**, cooperatively stabilizing the ternary complex. Collectively, these findings establish **IBA-11** as a highly orthogonal CK1α MGD. More importantly, they elegantly demonstrate that the indazolone architecture can be engineered to recruit discrete CRBN neo- substrates, underscoring its vast potential as a versatile, programmable platform for targeted protein degradation.

### Discovery of Potent Indazolone-Based MGD for Selective IKZF2 Degradation

Encouraged by the programmable substrate selectivity of the indazolone architecture, we sought to determine whether this platform could be broadly deployed to mine the vast repertoire of over 1,600 predicted CRBN neo-substrates. Exploiting the structural plasticity of the solvent-exposed region, we introduced a bulky 1-(3-chlorobenzyl)piperidin-4-yl moiety to generate **IBA-12**, aiming to further reprogram CRBN substrate specificity (Figure 7a).

**Figure 7.**
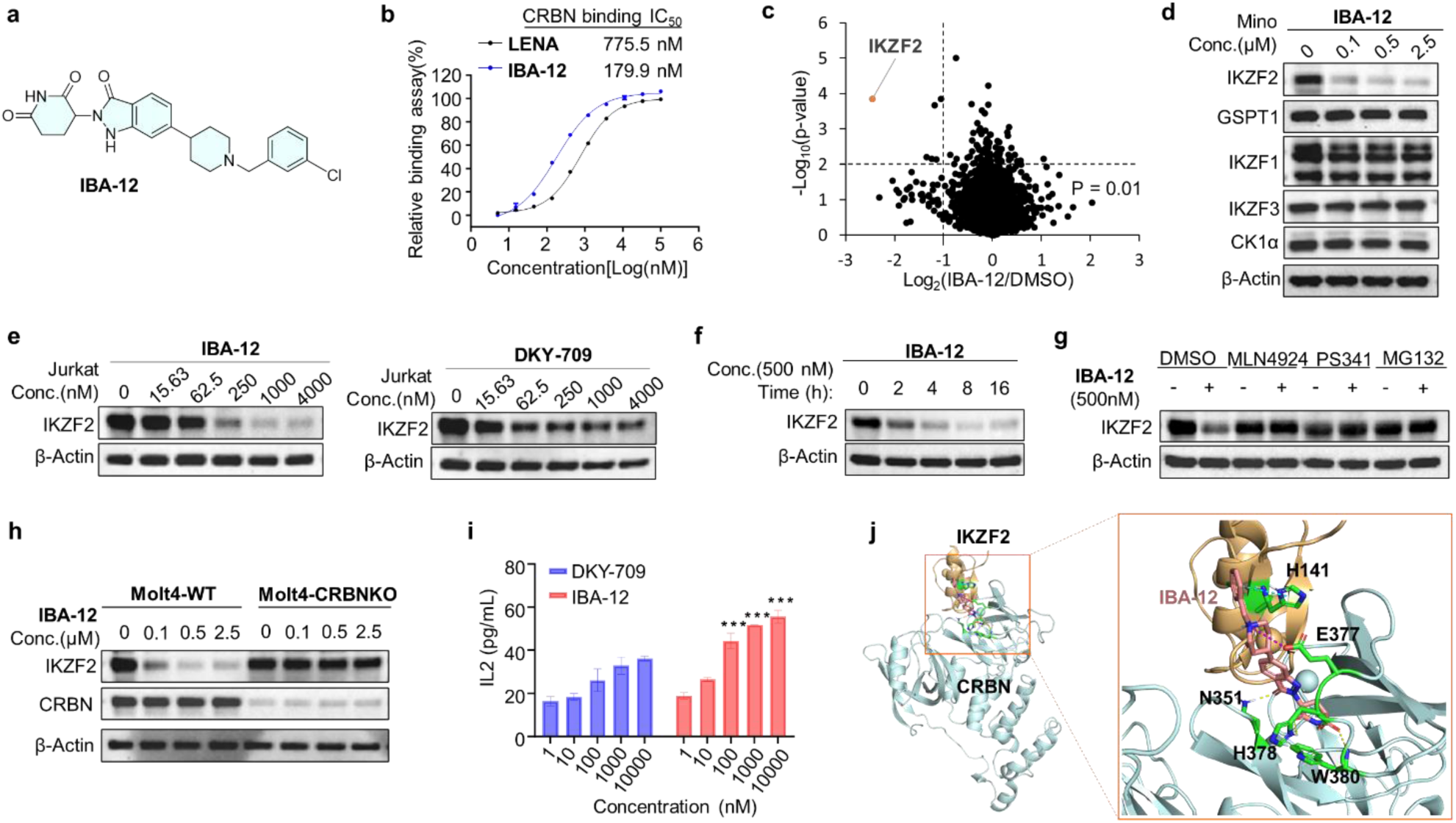
Biological characterization of the IKZF2-selective molecular glue IBA-12. **a)** Chemical structure of **IBA-12**. **b)** CRBN binding affinities of LENA and **IBA-12** determined by TR-FRET assay. Data are expressed as mean ± SD (n = 2 independent replicates). **c)** Proteomic analysis of Mino cells following treatment with 1 μM **IBA-12** for 8 h. Thresholds for significant protein alterations were set at Log_2_ fold change (**IBA-12**/DMSO) < -1 and p-value < 0.01 (n = 3). **d)** Western blot analysis of multiple CRBN substrates in Mino cells treated with the indicated concentrations of **IBA-12** for 8 h. **e)** Western blot analysis of IKZF2 levels in Jurkat cells treated with the indicated concentrations of **IBA-12** for 24 h. **f)** Time-dependent degradation profiles of IKZF2 in Mino cells treated with 500 nM IBA-12. **g)** Mino cells were pretreated with MLN4924 (0.5 μM), PS341 (1 μM), or MG132 (10 μM) for 2 h prior to treatment with 0.5 μM **IBA-12** for 4 h. **h)** Western blot analysis of IKZF1 and ZFP91 in wild-type Molt4 cells and CRBN^⁻/⁻^ Molt4 cells treated with **IBA-12**. **i)** Jurkat cells were stimulated with SEE-pulsed Raji cells in the presence or absence of DKY-709 or **IBA-12**. Culture supernatants were collected after 24 h and IL-2 production was quantified by ELISA. Data are presented as mean ± SEM (n = 3 biological replicates per concentration). “***” P < 0.001. **j)** Predicted binding mode of **IBA-12** (red) in complex with CRBN (cyan) and IKZF2 (wheat) based on molecular docking (PDB ID: 8DEY). **k)** Pharmacokinetic parameters of **IBA-12** in rats.

Evaluation via a TR-FRET assay revealed that **IBA-12** possesses potent CRBN binding affinity (IC50 = 179.9 nM), demonstrating a substantial enhancement over the parent ligand lenalidomide (IC50 = 775.5 nM) (Figure 7b). Motivated by this robust affinity for CRBN and the proven capacity of our platform to yield structurally diverse MGDs, we bypassed conventional phenotypic screening and directly profiled the cellular neo-substrate landscape via whole-cell proteomics. The proteome-wide analysis uniquely revealed IKZF2 as the most significantly and selectively downregulated protein in Mino cells (Figure 7c).

Western blot analysis confirmed the dose-dependent degradation of IKZF2 in Mino cells following a 24 h treatment with **IBA-12**, achieving substantial depletion at just 0.1 μM (Figure 7d). In striking contrast, other CRBN neo-substrates—including GSPT1, IKZF1, IKZF3, and CK1α—remained largely unperturbed even at concentrations up to 2.5 μM, highlighting the exquisite substrate orthogonality of **IBA-12**. In Mino cells, IBA-12 achieved a DC50 value of 461.8 nM, indicating potent degradation activity. Consistent results were observed in Jurkat cells, where IBA-12 induced dose-dependent IKZF2 degradation (DC50 = 276.2 nM, Dmax = 85.37%), demonstrating improved degradation efficiency compared with DKY-709 (Figure 7e). In addition, time-course experiments revealed progressive IKZF2 depletion, with 500 nM **IBA-12** inducing marked protein degradation within a rapid 4- to 8-hour window (Figure 7f).

Mechanistic investigations revealed that IKZF2 degradation was completely blocked by MLN4924 or the proteasome inhibitors PS341 and MG132 (Figure 7g), demonstrating a strict dependence on CRL activity and the UPS. Consistently, this degradation was observed in wild-type Molt4 cells but was completely abolished in CRBN-knockout cells, confirming CRBN dependency (Figure 7h). Given that IKZF2 is known to repress the transcription of interleukin-2 (IL-2) in regulatory T cells, we evaluated the functional consequences of its degradation.^35–38^ Accordingly, treatment of Jurkat cells stimulated by Raji cells with **IBA-12** or DKY-709^36^ resulted in dose-dependent increases in IL-2production (Figure 7i), indicating the successful functional modulation of an IKZF2-regulated transcriptional pathway. **IBA-12** also displayed favorable metabolic stability in rat liver microsomes, with no detectable metabolic turnover (Table S2). We next evaluated its pharmacokinetic properties in rats (Table S3). Following intravenous administration, **IBA-12** exhibited moderate clearance, a large volume of distribution. Oral administration led to rapid absorption, high systemic exposure, with a half-life of 2.54 h and excellent oral bioavailability (Table S3). Taken together, the **IBA-12** represents a highly promising candidate for further preclinical development.

To gain structural insight into **IBA-12**-mediated IKZF2 recruitment, molecular docking studies were performed using the reported IKZF2-CRBN crystal structure (PDB ID: 8DEY). As shown in Figure 7j, the glutarimide moiety of **IBA-12** occupies the canonical tri-tryptophan pocket of CRBN. The protonated nitrogen of the piperidine ring forms a salt bridge with Glu377 of CRBN, a feature also observed in the IKZF2 degrader DKY-709.^36^ At the CRBN–IKZF2 interface, **IBA-12** further establishes π–π stacking interactions with H141, stabilizing the IKZF2–**IBA-12**–CRBN ternary complex. Collectively, these results establish **IBA-12** as a potent and selective IKZF2 molecular glue degrader acting via a CRBN-dependent UPS mechanism pathway.

### Broader Implications and Future Potential of the Indazolone-Based MGD Discovery Platform

MGD has rapidly transformed drug discovery, yet a significant challenge persists: the predominant reliance on a limited number of chemical scaffolds, notably the canonical isoindolinone glutarimide, for recruiting the E3 ligase CRBN. This over-reliance inadvertently constrains the exploration of the vast degradable proteome, limiting both chemical diversity and the scope of therapeutically relevant targets accessible to MGDs. In this context, the indazolone-based MGD discovery platform presented in this work represents a crucial advancement, offering an innovative paradigm that directly addresses these inherent limitations and paves the way for a next generation of MGDs with enhanced substrate programmability and therapeutic potential.

Our research first highlights a profound implication by successfully expanding the chemical space that has long challenged CRBN-MGD discovery. Our efforts demonstrate that the indazolone architecture, a structurally distinct chemotype, serves as a highly effective CRBN binder. Through rigorous evaluation of compounds **IBA-1** to **IBA-7**, we have established this novel structural foundation for MGDs, dramatically expanding the chemical diversity available for modulating CRBN function. By moving beyond traditional scaffolds, our platform effectively broadens previously unexplored chemical space, which is critical for overcoming intrinsic architectural limitations and enabling access to a wider array of neo-substrates, thereby achieving superior pharmacological profiles.

The success of our platform is deeply rooted in design principles guided by mechanistic insights. We leveraged existing structural insights—the observed positional shift in the CRBN-**LENA**-CK1α complex—to inspire the rational design of the indazolone architecture. The intrinsic N-substitutability of the indazolone framework allows us to precisely occupy and strategically exploit this permissive space within the CRBN binding pocket. This strategy led to optimized binding affinities and, critically, differential neo-substrate recruitment. This structure-informed design approach represents a significant step forward, deepening our understanding of the structural principles governing MGD-E3-substrate ternary complex formation and providing a robust blueprint for future MGD architecture engineering.

Crucially, this indazolone platform offers exceptional programmability of CRBN substrate specificity, addressing a central challenge in MGD development: the ability to precisely dictate which neo-substrate is recruited for degradation. Our results provide compelling evidence that subtle structural modifications within the indazolone architecture leading to dramatic shifts in degradation profiles. We have systematically demonstrated how compounds such as **IBA-8**, **IBA-9**, **IBA-10**, **IBA-11**, and **IBA-12**, despite their structural kinship, elicit distinct and often highly selective degradation patterns. For instance, **IBA-8** functions as a potent multi-target degrader of IKZF1/3, CK1α, IKZF2, ZFP91, and LIMD1. Remarkably, modest structural refinements in **IBA-9** reprogram this profile, leading to a more selective degradation of IKZF1/3, CK1α, and ZFP91. Most strikingly, **IBA-12** achieved exquisite selectivity for IKZF2, without affecting other canonical CRBN substrates such as IKZF1/3 or GSPT1. This remarkable ability to fine-tune neo-substrate recognition through structural diversification establishes our indazolone platform as a powerful tool for developing MGDs to target new proteins of interest relevant to specific therapeutic needs.

The broad biomedical application potential of this platform is immense. By generating highly potent and selective degraders for therapeutically relevant targets such as IKZF1/3, ZFP91, CK1α, and IKZF2, our work directly addresses unmet medical needs in various diseases. For instance, the selective degradation of CK1α by **IBA-11** opens promising new avenues for therapy for acute myeloid leukemia (AML), while the highly specific IKZF2 degrader **IBA-12** holds significant promise for immunomodulation and for treating cancers where precise targeting is critical. The ability to generate such diverse pharmacological profiles from a single, tunable scaffold family suggests that the indazolone platform can effectively expand the druggable proteome by converting previously undruggable proteins into tractable therapeutic targets, thereby offering novel therapeutic strategies for challenging diseases.

Looking forward, the human proteome is estimated to harbor a vast and expanding repertoire of over 1,600 computationally predicted CRBN neo-substrates,^12,16,17^ underscoring the immense, yet largely untapped, landscape for MGD discovery. Within this context, our indazolone-based platform provides a compelling and scalable framework for future CRBN-mediated MGD development. The pronounced tunability and validated degradation efficacy of the indazolone architecture highlight its strong potential for systematic expansion, including strategic substituent diversification, stereochemical modulation, and hybrid scaffold design to achieve increasingly precise control over substrate specificity and degradation efficiency. Future studies will focus on elucidating the structural and mechanistic basis underlying indazolone-driven substrate selectivity, particularly through advanced structural biology approaches such as co-crystal or cryo-EM analyses of ternary CRBN–MGD–substrate complexes to resolve interaction networks at atomic resolution. In parallel, the encouraging drug-like and pharmacokinetic properties observed for representative compounds (e.g., **IBA-10** and **IBA-12**) support continued PK/PD optimization toward preclinical development. Notably, CRBN has emerged as one of the most extensively exploited E3 ligases in PROTAC technology, enabling the degradation of a broad spectrum of disease-relevant targets across oncology, immunology, neurodegeneration, and virology.^1–4^ In this regard, the indazolone scaffold represents a new class of CRBN-binding chemotypes, offering versatile entry points for the rational design of next-generation PROTACs. Collectively, this platform not only expands the chemical and functional landscape of CRBN-based degraders but also establishes a generalizable strategy for systematically addressing previously intractable therapeutic targets.

## Conclusion

In summary, our work introduces an innovative indazolone-based platform that critically advances MGD discovery by extending beyond canonical isoindolinone-glutarimide scaffolds and significantly expanding the accessible chemical space. By leveraging mechanistic insights into CRBN-MGD complex conformational plasticity, we rationally designed indazolone architectures that establish a robust foundation for next-generation MGDs. Crucially, this platform offers highly tunable programmability of CRBN substrate specificity, enabling fine-tuned control of neo-substrate recognition. This versatility is demonstrated through diverse degradation profiles, ranging from the broad targeting of critical proteins—such as IKZF1/3, ZFP91, and LIMD1—to the exquisitely selective degradation of CK1α and IKZF2. Beyond expanding the chemical landscape for targeting previously undruggable proteins, this innovation suggests transformative therapeutic strategies for various cancers. Collectively, the indazolone architecture’s remarkable tunability, potent degradation efficacy, and favorable pharmacokinetic properties provide a powerful blueprint for future CRBN-based discovery. Furthermore, these architectures could offer novel CRBN ligands for the rational design of diverse PROTACs, broadening the scope of TPD and paving the way for systematically addressing intractable diseases.

## Supporting Information

Supporting Figures, ^1^H and ^13^C NMR spectra for all compounds, additional experimental details, materials and methods, and tables.

## Notes

The authors declare no competing financial interest.

## Acknowledgements

This study was supported by the National Science Foundation of China (No. 22377136 to X.-H. C); the Strategic Priority Research Program of the Chinese Academy of Science (XDB1260202 to X.-H. C.). the Science and Technology Commission of Shanghai Municipality, China (No.25ZR1402553 to H.-J. N).

