## Supplementary Information for "Indazolone-Based Molecular Glue Degraders as a Tunable Platform for Reprogramming Cereblon Substrate Specificity"

---

[a] Dr. H.-J. Nie, H. Xu, Y.-J. Zhou, G.-L. Yin, Prof. X.-H. Chen
State Key Laboratory of Drug Research
Shanghai Institute of Materia Medica, Chinese Academy of Sciences
Shanghai, 201203, China


[b] J. Wang, G. Xu, H. Hu, Prof. J. Li
State Key Laboratory of Chemical Biology
Shanghai Institute of Materia Medica, Chinese Academy of Sciences
Shanghai, 201203, China


[c] X. Xu, B. Chen, X. Li, X. Hu, Y. Zhou, J. Li
Zhongshan Institute for Drug Discovery
Shanghai Institute of Materia Medica, Chinese Academy of Sciences
Zhongshan, 528400, China

[d] Prof. X.-H. Chen
School of Pharmaceutical Science and Technology
Hangzhou Institute for Advanced Study, University of Chinese Academy of Sciences
Hangzhou, 310024, China

[e] H. Xu, Y.-J. Zhou
University of Chinese Academy of Sciences
Beijing, 100049, China

‡ These authors contributed equally to this work.

### Table of Contnets

### Supplementary Figures

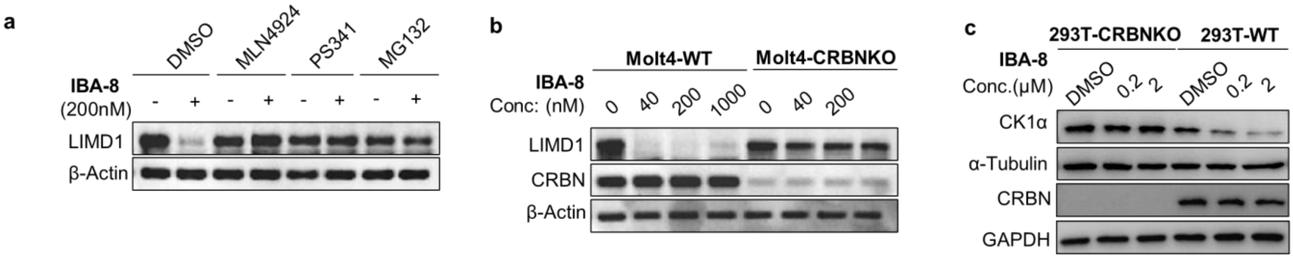

**Figure S1. a)** Mechanistic investigation of target degradation. Mino cells were pretreated with DMSO, MLN4924 (0.5 μM), PS341 (1 μM), or MG132 (10 μM) for 2 h, followed by treatment with 200 nM **IBA-8** for 3 h. Protein levels of LIMD1 was subsequently evaluated by immunoblotting. **b)** Western blot analysis of LIMD1 in wild-type Molt4 cells and CRBN$^{-/-}$ Molt4 cells following treatment with **IBA-8** for 4h. **c)** Western blot analysis of CK1α in wild-type (WT) and CRBN$^{-/-}$ HEK293T cells following treatment with **IBA-8** for 4h.

#### Supplementary Tables

##### Table S1. Cell viability assay of IBA-2 to IBA-7 in Mino, MM.1S and MV-4-11 cells.

| Compound | $IC_{50}$(nM) | | |
|---|---|---|---|
| | Mino | MM.1S | MV-4-11 |
| **IBA-2** | 9044 | 351.3 | >20000 |
| **IBA-3** | >20000 | 160.1 | >20000 |
| **IBA-4** | >20000 | >20000 | >20000 |
| **IBA-5** | 2377 | 150.8 | >20000 |
| **IBA-6** | >20000 | 30.36 | >20000 |
| **IBA-7** | >20000 | >20000 | >20000 |
| **LENA** | >20000 | 91.65 | >20000 |
| **CC-220** | 77.8 | 8.55 | 9181 |

##### Table S2. Metabolic Stability of Compounds IBA-2 to IBA-12 in Liver Microsomes

| Compound | species | $t_{1/2}$(min) | $Cl_{int}$(mL/min/mg) |
|---|---|---|---|
| **IBA-2** | rat | 25.5 | 0.1361 |
| **IBA-3** | rat | 1.61 | 2.1460 |
| **IBA-5** | rat | 1095 | 0.0032 |
| **IBA-6** | rat | 138 | 0.0251 |
| **IBA-8** | rat | 51.1 | 0.0678 |
| **IBA-9** | rat | ∞ | 0 |
| **IBA-10** | rat | 58 | 0.0599 |
| **IBA-11** | rat | 184 | 0.0189 |
| **IBA-12** | rat | ∞ | 0 |
| Testosterone | rat | <5 | / |

##### Table S3. Pharmacokinetic parameters of IBA-12 in rats.

| **IBA-12** | Dose (mg/kg) | $T_{1/2}$ (h) | $T_{max}$ (h) | $C_{max}$ (ng/mL) | $AUC_{0-t}$ (h·ng/mL) | $V_{ss}$ (L/kg) | CL (mL/min/kg) | $MRT_{0-\infty}$ (h) | F(%) |
|---|---|---|---|---|---|---|---|---|---|
| **IV** | 2 | 3.4 | 0.083 | 273 | 651 | 9.42 | 50.8 | 3.10 | |
| **PO** | 10 | 2.54 | 1.33 | 1067 | 3789 | / | / | 2.62 | 116 |

### Supplementary Methods

#### 1.General Information

Unless otherwise noted, all chemical reagents and solvents were obtained from commercial suppliers and used as received without further purification. RPMI 1640 medium (Gibco, Cat. No. 31800-89), IMDM medium (Gibco, Cat. No. 12200-069), Fetal Bovine Serum (Gibco, Cat. No. A5256701) were purchased from Thermo Scientific. CCK-8 (Cat. No. MA0218-3) was purchased from MeilunBio. 4×Laemmli sample buffer (Cat. No. 161-0747) was purchased from Bio-Rad. GSPT1 antibody (Cat. No. 14980), IKZF1 antibody (Cat. No. 14859), IKZF2 antibody (Cat. No. 42427), IKZF3 antibody (Cat. No. 15103) was purchased from Cell Signaling Technology. CK1α antibody (Cat. No. 65200) was purchased from Genuinbiotech. ZFP91 antibody (Bethyl, Cat. No. A303-245A) was purchased from Univ-bio. LIMD1 (Cat. No. 28106-1-AP) was purchased from Proteintech. β-Actin antibody (Cat. No. AM1021B) was purchased from Abcepta. ECL kit (Cat. No.KF8003) was purchased from Affinity Biosciences. CD4 monoclonal antibody (OKT4), APC (Cat. No. 17-0048-42), CD4 monoclonal antibody (RM4-5), Super Bright 600 (Cat. No. 63-0042-82), CD3e monoclonal antibody (145-2C11), PE-Cyanine7 (Cat. No. 25-0031-82), FOXP3 monoclonal antibody (FJK-16s), PE (Cat. No. 12-5773-82), and the Foxp3/Transcription Factor Staining Buffer Set (Cat. No. 00-5523-00) were purchased from eBioscience™. FITC anti-mouse/human Bcl6 antibody (clone 22F6) (Cat. No. 137214) was obtained from BioLegend. PE-Cy7™ mouse anti-human CD3 (Cat. No. 563423) and Fixable Viability Stain 780 (Cat. No. 565388) were purchased from BD Pharmingen™. Human IL-2 recombinant protein (Cat. No. 200-02-10UG) was obtained from PeproTech. LIVE/DEAD Fixable Viability Dye (Cat. No. L34966) was purchased from Life Technologies. RNAiso Plus (TaKaRa, Cat. No. 9108), Evo M-MLV RT Mix Kit (Accurate Biology, Cat. No. AG1178), SYBR Green Premix Pro Taq HS qPCR kit (Accurate Biology, Cat. No. AG11718). The Human IL-2 Uncoated ELISA Kit (Cat. No. 88-7025-88, RRID: AB_2574954) was purchased from Invitrogen. $^{1}$H NMR spectra and $^{13}$C NMR spectra were obtained on a Bruker AVANCE III 400 (400 MHz), BRUKER AVANCE NEO 500, Bruker Avance NEO 500 (Cryo) Spectrometer. Chemical shifts were reported in parts per million (ppm) on the δ scale from an internal standard (NMR descriptions: s, singlet; d, doublet; t, triplet; q, quartet; m, multiplet; br, broad). Coupling constants, *J*, are reported in Hertz. For HR-MS analysis of small molecules, samples were analyzed by flow-injection analysis into an Agilent 1290-6545 UPLC-QTOF. LC-MS analysis of small molecules was performed on Waters UPLC-MS (UPLC: Waters HPLC H-CLASS, MS: Waters SQ Detector 2) with ACQUITY UPLC BEH C18 column (1.7 μm, Waters).

#### 2. Cell lines and cell culture

MM.1S (1640+10%FBS), Mino (1640+15%FBS) and MV-4-11 (IMDM+10%FBS) was purchased from the American Type Culture Collection. Jurkat cells (RRID: CVCL_0065, Cat. No. CL-0129) and raji cells (RRID: CVCL_0511, Cat. No. TCHU44) were cultured in complete RPMI 1640 with 10% FBS. MC38 cells（RRID:CVCL_B288, Cat. No.SNL-505）were cultured in DMEM with 10% FBS and 1%NEAA. PBMC (LDEBIO, Cat. No. 1501) were culture in complete RPMI 1640 with 10% FBS, 10ng/ml IL2 and 50 μM 2-Mercaptoethanol. All cell lines were cultured at 37 °C in an atmosphere of 5% $CO_2$.

#### 3. Cellular Viability Assay

Cells in the logarithmic growth phase were seeded in 96-well culture plates, and the appropriate number of cells was seeded according to the growth rate of the cells. Each well contained 200 μL of culture medium with the corresponding concentration of compounds. After 7 days of culture, 10 μL of CCK-8 was added to each well. After incubation for 2 h, the values were read using a SpectraMax 190 (Molecular Device) microplate reader. The difference in absorbance values between 450 nm and 650 nm was used as the final data. Finally, the $IC_{50}$ value of the compound was calculated by nonlinear regression method using GraphPad 10.1.2 software. Data results are from three independent replicate experiments.

#### 4. Western blot assay

Cells in the logarithmic growth phase were seeded in 12-well culture plates, 5 x $10^5$ cells per well, and cultured with 1 mL culture medium containing the corresponding concentrations of the compounds. After processing, the cells were collected by centrifugation. Cells were washed once with precooled PBS followed by centrifugation to remove PBS, 200 μL loading buffer was added, cells were resuspended, and samples were boiled and denatured at 100 °C for 25 min. Appropriate amounts of protein samples were subjected to Tricine-SDS-PAGE for electrophoresis. After electrophoresis, the proteins were transferred to nitrocellulose membranes, followed by blocking, overnight incubation of the primary antibody at 4 °C, incubation of the secondary antibody in the next day, and finally exposure. Image Lab 6.0.1 was used to perform grayscale analysis.

#### 5. Molecular Docking

##### 5.1 Molecular Modeling

All molecular modeling and docking studies were performed using the Schrödinger software suite (Release 2025-2). The crystal structures of CRBN (PDB ID: 4TZ4), CK1α–CRBN (PDB ID: 5FQD), and IKZF2–CRBN (PDB ID: 8DEY) were obtained from the Protein Data Bank (PDB). All protein complexes were prepared and optimized using the Protein Preparation with default parameters (pH 7.4 ± 2.0). The ligands (IBA series) were prepared using LigPrep to generate low-energy conformations and correct ionization states.

The receptor grid boxes for all systems were centered on the original co-crystallized ligands. Grid dimensions were automatically defined to accommodate ligands of similar size to the workspace ligand. Docking calculations were conducted using Glide SP (Standard Precision) mode with enhanced sampling enabled. Post-docking minimization was performed on the top 10 ranked poses for each system.

Binary Docking: For the CRBN–ligand systems, the ligands **IBA-2**, **IBA-3**, and **IBA-4** were docked into the CRBN template (PDB ID: 4TZ4).

Ternary Docking: To model the ternary arrangements involving specific neo-substrates, the crystal structures of the CK1α–CRBN complex (PDB ID: 5FQD) and the IKZF2–CRBN complex (PDB ID: 8DEY) were utilized as templates for docking **IBA-11** and **IBA-12**, respectively.

The final docking results and molecular interactions were visualized and analyzed using the PyMOL Molecular Graphics System.

##### 5.2 Ternary Complex Modeling via Protein–Protein Docking

To investigate the ternary binding modes, compounds IBA-8 and IBA-10 were first docked into the crystal structure of CRBN (PDB ID: 4TZ4) using the Glide SP (Standard Precision) module within the Schrödinger software suite (Release 2025-2) with default parameters. The top-ranked poses were selected to construct the ligand-bound CRBN receptor models for subsequent protein–protein docking.

The three-dimensional structures of the target proteins, ZFP91 (UniProt ID: Q96JP5) and LIMD1/Leiomodin-1 (UniProt ID: Q9UGP4), were retrieved from the AlphaFold 3 (AF3) protein structure database.

The recruitment of neo-substrates was modeled using the Protein–Protein Docking module in Schrödinger. The pre-assembled CRBN–IBA-8 and CRBN–IBA-10 complexes were defined as the receptor units, while LIMD1 and ZFP91 were utilized as the ligand units, respectively. For each system, the representative binding mode was identified by selecting the conformation from the largest cluster size, ensuring the statistical significance of the predicted pose.

The resulting ternary complex structures were further optimized using the Interface Refinement tool to enhance the local complementarity and interaction energies at the binding interface. structural analysis and visualization of the ternary arrangements and molecular interfaces were performed using the PyMOL Molecular Graphics System.

#### 6. Proteomic Sample Preparation and DIA-MS Analysis

Take out the samples in the frozen state and put it on ice. The samples were lysed in buffer containing 8 M urea and 1% SDS supplemented with protease inhibitors, homogenized by tissue grinding (3 × 180 s), and further disrupted by non-contact cryogenic sonication for 30 min. After centrifugation (16,000 × *g*, 8 °C, 30 min), the supernatants were collected, and protein concentrations were determined using a BCA assay kit (Thermo Scientific). Protein quality was assessed by SDS-PAGE.

For protein digestion, 100 μg of each sample was resuspended in 100 mM triethylammonium bicarbonate (TEAB), reduced with 10 mM tris(2-carboxyethyl)phosphine at 37 °C for 1 h, and alkylated with 40 mM iodoacetamide at room temperature for 40 min in the dark. Proteins were then digested with trypsin (enzyme/protein = 1:50, *w/w*) at 37 °C overnight. The resulting peptides were dried under vacuum, reconstituted in 0.1% trifluoroacetic acid, desalted using HLB cartridges, and quantified by UV absorbance on a NanoDrop One instrument (Thermo Scientific).

Peptides were analyzed on a Vanquish Neo system coupled to an Orbitrap Astral mass spectrometer (Thermo Fisher Scientific) at Majorbio Bio-Pharm Technology Co., Ltd. (Shanghai, China). Separation was performed on a uPAC high-throughput column (75 μm × 5.5 cm) using solvent A ($H_2O$ with 2% MeCN and 0.1% formic acid) and solvent B ($H_2O$ with 80% MeCN and 0.1% formic acid) with a total run time of 8 min. DIA data were acquired over an *m/z* range of 100–1700.

Raw data were processed with Spectronaut 19 using trypsin/P specificity, up to two missed cleavages, carbamidomethylation of cysteine as a fixed modification, and methionine oxidation and protein N-terminal acetylation as variable modifications. Protein and peptide FDRs were both controlled at 1%. Protein quantification was performed using MaxLFQ. P-values and Fold change (FC) for the proteins between the two groups were calculated using R package “t-test”.Statistical analysis was carried out on the Majorbio Cloud platform. Differentially expressed proteins were defined using thresholds of fold change < 0.5 and *P* < 0.01.

#### 7.RT-qPCR

Total RNA was isolated using RNAiso Plus and reverse transcription was performed using the Evo M-MLV RT Mix Kit according to manufacturer’s instructions. RT-qPCR was performed using SYBR Green Premix Pro Taq HS qPCR kit in Stratagene™ Mx3005P system (Agilent Technologies). Data were analyzed using the $2^{-\Delta\Delta CT}$ method. Target mRNA was normalized to the corresponding *GAPDH* mRNA. Primers used in the reaction are listed in **Table S3**.

**Table S3. Primer sequences used for RT-qPCR**

| Primer | Sequence(5’-3’) |
|---|---|
| ZFP91 | Forward: TCCTTGCCCATCCTCGCTATT |
| | Reverse: TGTTTGGCATGTCGCAGAAGT |
| LIMD1 | Forward: TGGGGAACCTCTACCATGAC |
| | Reverse: CACAAAACACTTTGCCGTTG |
| GAPDH | Forward: GTCTCCTCTGACTTCAACAGCG |
| | Reverse: ACCACCCTGTTGCTGTAGCCAA |

#### 8. Liver Microsomal Stability.

##### 8.1 Preparation of Solutions

Stock solution (10 mM) of test compounds was prepared in DMSO. The stock solution for test compound was then diluted into 200 µM with acetonitrile.

##### 8.2 Microsome Incubations

Incubation mixtures were prepared in a total volume of 200 μL with final component concentrations as follows: 0.1 M PBS (pH 7.4), 3 mM $MgCl_2$, NADPH (2 mM), liver microsomes (0.2 mg/mL) and test compound (1 μM) or positive control (1 μM). NADPH or buffer (negative control) was added after a 5-min preincubation of all other components at 37 °C. Pipette-mixed to achieve a homogenous suspension and immediately transferred 20 μL incubate as a 0 min sample to wells in a "Quenching" plate followed by pipette-mixing. At 5, 15, 30, and 60 min, pipette-mixed the incubate and serially transfer samples of 20 μL incubate per time point to wells in a separate "Quenching" plate followed by pipette-mixing. In 'Quenching' plates added 200 μL of acetonitrile with IS.

##### 8.3 Sample Analysis

The 96-well plate was centrifuged at 4000 rpm, 4 °C for 10 min. 30 μL of supernatant was mixed with 120 μL of $ddH_2O$ and then injected onto the LC-MS/MS system for analysis.

#### 9. Pharmacokinetic Study *In Vivo*.

A pharmacokinetic study was conducted on male SD rats. Test compounds were dissolved in 90% saline containing 5% DMSO and 5% Solutol HS-15 for IV dose administration and 90% distilled water containing 5% DMSO and 5% Solutol HS-15 for PO dose administration. For each compound, the rats were given by intravenous administration or oral gavage at a dose of 2 and 10 mg/kg, respectively. Blood samples were harvested through an internal jugular vein into EDTA-$K_2$ tubes at 0.0833, 0.25, 0.5, 1, 2, 4, 7, and 24 h for intravenous administration and oral gavage. Samples were centrifuged at 5000 rpm for 10 min to separate the plasma. The plasma samples were stored at -70 °C until liquid chromatography with tandem mass spectrometry (LC-MS/MS) analysis. The pharmacokinetic parameters were performed with Phoenix WinNonlin (version 8.1, Pharsight; Princeton, NJ). The half-life ($t_{1/2}$), area under the curve (AUC), clearance (CL), mean retention time (MRT), and bioavailability (F) were calculated as shown in the equation.

$$F\ (\%) = (AUC_{0\text{–}t},PO \times DoseIV) / (AUC_{0\text{–}t},IV \times DosePO) \times 100$$

#### 10. IL2 production of Jurkat cell:

Jurkat cells were stimulated with SEE-pulsed Raji cells at a ratio of 1 to 1, with or without DKY-709/IBA-12, supernatant of culture medium was collected after 24 hours, and its IL2 production were measured using commercial kits (Invitrogen, Cat# 88-7025-88).

#### Synthesis methods

##### Synthesis of IBA-2

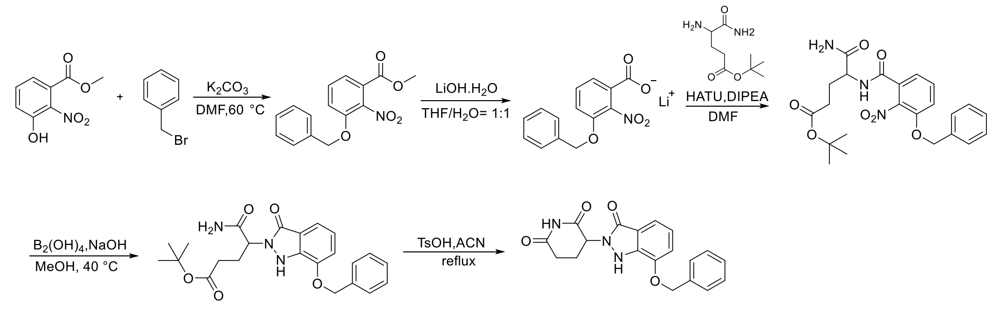

IBA-2

Methyl 3-hydroxy-2-nitrobenzoate (500 mg, 2.54 mmol), benzyl bromide (603 μL, 5.07 mmol), and $K_2CO_3$ (1.05 g, 7.61 mmol) were stirred in N,N-dimethylformamide (DMF, 5 mL) at 60 °C for 3 h. The mixture was cooled to room temperature, diluted with $H_2O$, and extracted with ethyl acetate. The combined organic layers were washed with brine, dried over $Na_2SO_4$, filtered, and concentrated under reduced pressure. Purification by silica gel column chromatography afforded the product as a white solid (727 mg, 99%).$^1$H NMR (500 MHz, DMSO) δ 7.73 (dd, J = 8.4, 1.2 Hz, 1H), 7.68 (t, J = 8.1 Hz, 1H), 7.58 (dd, J = 7.7, 1.2 Hz, 1H), 7.43 – 7.30 (m, 5H), 5.33 (s, 2H), 3.84 (s, 3H).

Methyl 3-(benzyloxy)-2-nitrobenzoate (727 mg, 2.53 mmol) was dissolved in tetrahydrofuran (THF, 4 mL), and an aqueous solution of LiOH·$H_2O$ (319 mg, 7.59 mmol, in 4 mL $H_2O$) was added. The reaction was stirred at room temperature overnight. The mixture was concentrated under reduced pressure, and the crude material was used directly in the next step without purification.

Lithium 3-(benzyloxy)-2-nitrobenzoate, tert-butyl 4,5-diamino-5-oxopentanoate (604 mg, 2.53 mmol), and HATU (962 mg, 2.53 mmol) were dissolved in DMF (10 mL). DIPEA (1.4 mL, 7.59 mmol) was added dropwise, and the mixture was stirred at room temperature for 2 h. The reaction mixture was diluted with ethyl acetate, washed with $H_2O$ and brine, dried over $Na_2SO_4$, filtered, and concentrated. Purification by silica gel column chromatography afforded the product as a white solid (1.15 g, 99%). $^1$H NMR (500 MHz, DMSO) δ 8.75 (d, J = 8.1 Hz, 1H), 7.61 (dd, J = 8.5, 7.6 Hz, 1H), 7.55 (dd, J = 8.5, 1.2 Hz, 1H), 7.43 – 7.37 (m, 5H), 7.37 – 7.32 (m, 2H), 7.15 – 7.08 (m, 1H), 5.31 (d, J = 1.1 Hz, 2H), 4.27 (ddd, J = 9.4, 8.0, 4.9 Hz, 1H), 2.30 – 2.22 (m, 2H), 2.02 – 1.93 (m, 1H), 1.87 – 1.75 (m, 1H), 1.39 (s, 9H).

Tert-Butyl 5-amino-4-(3-(benzyloxy)-2-nitrobenzamido)-5-oxopentanoate (1.15 g, 2.51 mmol) was dissolved in methanol (31 mL) and cooled to 0 °C. Following a reported procedure,[1] tetrahydroxydiboron (1.01 g, 11.26 mmol) was added, and the mixture was stirred for 10 min at 0 °C. A solution of NaOH (1.01 g, 25.14 mmol) in MeOH (31 mL) was then added, and the reaction was stirred for 10 min at 0 °C and subsequently at 40 °C for 1 h. After LC-MS indicated completion, the mixture was cooled to room temperature, quenched with saturated $NH_4Cl$ solution, and extracted with ethyl acetate. The organic layer was washed with $H_2O$ and brine, dried over $Na_2SO_4$, filtered, and concentrated under reduced pressure. Purification by silica gel column chromatography afforded the product as a white solid (824 mg, 77%). $^1$H NMR (500 MHz, DMSO) δ 9.80 (s, 1H), 7.58 – 7.51 (m, 2H), 7.46 (s, 1H), 7.41 (dd, J = 8.2, 6.6 Hz, 2H), 7.38 – 7.33 (m, 1H), 7.31 (s, 1H), 7.24 (d, J = 7.7 Hz, 1H), 7.18 (d, J = 7.7 Hz, 1H), 7.05 (t, J = 7.8 Hz, 1H), 5.25 (s, 2H), 4.88 (dd, J = 9.7, 5.9 Hz, 1H), 2.28 – 2.16 (m, 2H), 2.12 – 1.98 (m, 2H), 1.35 (s, 9H).

Tert-Butyl 5-amino-4-(7-(benzyloxy)-3-oxo-1,3-dihydro-2H-indazol-2-yl)-5-oxopentanoate (400 mg, 0.94 mmol) was dissolved in THF and cooled to 0 °C. Potassium tert-butoxide (1.0 M in THF, 1.9 mL) was added dropwise, and the reaction mixture was stirred at room temperature for 1 h. The mixture was neutralized to pH 7 with 1 M HCl, diluted with $H_2O$, and extracted with ethyl acetate. The organic layer was washed with brine, dried over $Na_2SO_4$, filtered, and concentrated under reduced pressure. Purification by silica gel column chromatography afforded **IBA-2** as a white solid (246 mg, 74%). $^1$H NMR (500 MHz, DMSO) δ 10.97 (s, 1H), 10.11 (s, 1H), 7.54 (d, J = 7.1 Hz, 2H), 7.44 – 7.38 (m, 2H), 7.36 (d, J = 7.3 Hz, 1H), 7.25 (d, J = 7.5 Hz, 1H), 7.20 (d, J = 7.6 Hz, 1H), 7.06 (t, J = 7.8 Hz, 1H), 5.30 (dd, J = 13.1, 5.2 Hz, 1H), 5.25 (s, 2H), 2.92 (ddd, J = 17.2, 13.7, 5.4 Hz, 1H), 2.64 – 2.58 (m, 1H), 2.57 – 2.52 (m, 1H), 2.11 – 1.98 (m, 1H). $^{13}$C NMR (126 MHz, DMSO) δ 172.77, 169.62, 162.76, 145.15, 137.77, 136.54, 128.45, 128.02, 127.87, 121.74, 118.55, 114.83, 112.97, 69.58, 53.87, 30.93, 22.36. HRMS (ESI-Q-TOF): calcd. for $C_{19}H_{18}N_3O_4$ $[M + H]^+$ 352.1292, found 352.1293.

**Synthesis of IBA-1**

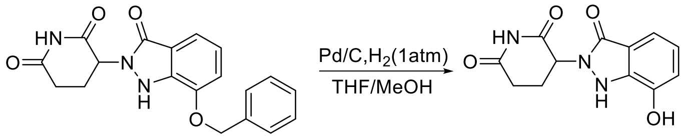

IBA-2 IBA-1

**IBA-2** (105 mg, 0.30 mmol) was dissolved in THF/MeOH (1:1, 8 mL). Pd/C was added, and the mixture was degassed with nitrogen three times and then with hydrogen three times. The reaction mixture was stirred under a hydrogen atmosphere (1atm) at room temperature overnight. The reaction mixture was filtered through Celite, concentrated under reduced pressure. Purification by silica gel column chromatography afforded **IBA-1** as a white solid (60 mg, 76%). $^{1}$H NMR (500 MHz, DMSO) δ 10.97 (s, 1H), 10.09 (s, 1H), 9.72 (s, 1H), 7.11 (dd, J = 7.3, 1.4 Hz, 1H), 6.98 – 6.91 (m, 2H), 5.28 (dd, J = 13.1, 5.2 Hz, 1H), 2.92 (ddd, J = 17.1, 13.7, 5.4 Hz, 1H), 2.62 (ddd, J = 17.4, 4.7, 2.7 Hz, 1H), 2.46 (dd, J = 13.3, 4.5 Hz, 1H), 2.07 (ddd, J = 11.8, 5.9, 3.4 Hz, 1H). $^{13}$C NMR (126 MHz, DMSO) δ 172.81, 169.66, 163.43, 144.09, 137.71, 122.02, 119.09, 115.92, 113.17, 53.92, 30.93, 22.37. HRMS (ESI-Q-TOF): calcd. for $C_{12}H_{12}N_3O_4$ [M + H] $^{+}$ 262.0822, found 262.0821.

**Synthesis of IBA-3**

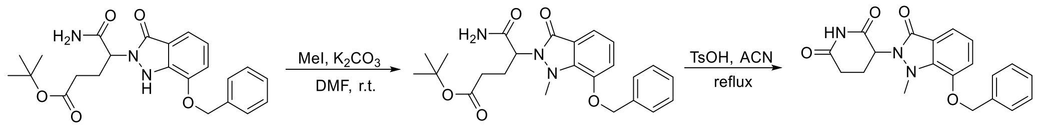

Tert-Butyl 5-amino-4-(7-(benzyloxy)-3-oxo-1,3-dihydro-2H-indazol-2-yl)-5-oxopentanoate (638 mg, 1.50 mmol) was dissolved in DMF (5 mL). $K_2CO_3$ (311 mg, 2.25 mmol) and Iodomethane (112 µL, 1.80 mmol) were added, and the reaction mixture was stirred at room temperature overnight. After LC–MS indicated completion, the mixture was filtered, and the filtrate was diluted with ethyl acetate. The solution was washed with $H_2O$ and brine, and the organic layer was dried over $Na_2SO_4$, filtered, and concentrated under reduced pressure. The crude product was used directly in the next step without further purification.

The crude product was dissolved in acetonitrile (ACN, 5 mL). A solution of *p*-Toluenesulfonic acid monohydrate (2.0 g, 12 mmol) in ACN (10 mL) was added, and the reaction mixture was stirred at 80 °C for 6 h. The solvent was removed under reduced pressure, and the residue was purified by preparative HPLC to afford **IBA-3** as a white solid (215 mg, 39% yield over two steps). $^{1}$H NMR (400 MHz, DMSO) δ 11.03 (s, 1H), 7.54 – 7.48 (m, 2H), 7.47 – 7.40 (m, 2H), 7.40 – 7.36 (m, 1H), 7.36 – 7.31 (m, 1H), 7.30 – 7.26 (m, 1H), 7.21 (t, *J* = 7.7 Hz, 1H), 5.28 (d, *J* = 2.0 Hz, 2H), 5.14 (dd, *J* = 12.5, 5.3 Hz, 1H), 3.24 (s, 3H), 2.89 – 2.76 (m, 1H), 2.73 – 2.56 (m, 2H), 2.10 – 2.00 (m, 1H). $^{13}$C NMR (101 MHz, DMSO) δ 172.67, 169.85, 163.27, 146.09, 141.75, 136.34, 128.60, 128.15, 127.84, 124.15, 120.99,

114.97, 114.87, 70.11, 54.07, 31.13, 22.67. HRMS (ESI-Q-TOF): calcd. for $C_{20}H_{20}N_3O_4$ [M + H] $^+$ 366.1448, found 366.1450.

**Synthesis of IBA-4**

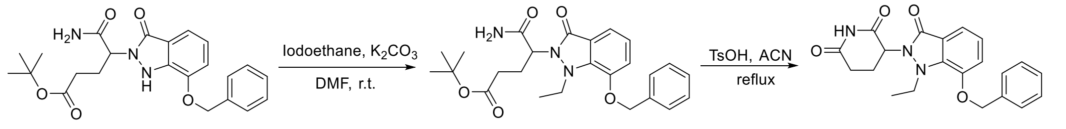

**IBA-4** was synthesized following the same procedure as described for **IBA-3**, using tert-Butyl 5-amino-4-(7-(benzyloxy)-3-oxo-1,3-dihydro-2H-indazol-2-yl)-5-oxopentanoate (638 mg, 1.5 mmol) and Iodoethane (144 μL, 1.80 mmol) as the starting materials. **IBA-4** was obtained as a white solid (236 mg, 41% yield over two steps). $^1$H NMR (400 MHz, DMSO) δ 11.02 (s, 1H), 7.50 (d, J = 7.1 Hz, 2H), 7.43 (t, J = 7.3 Hz, 2H), 7.38 (d, J = 7.2 Hz, 1H), 7.32 (d, J = 7.7 Hz, 1H), 7.27 (d, J = 7.2 Hz, 1H), 7.18 (t, J = 7.8 Hz, 1H), 5.28 (s, 2H), 5.07 (dd, J = 12.3, 5.3 Hz, 1H), 4.09 (dq, J = 13.7, 6.7 Hz, 1H), 3.84 – 3.66 (m, 1H), 2.86-2.74(m, 1H), 2.71-2.55 (m, 2H), 2.09 – 1.98 (m, 1H), 0.70 (t, J = 6.8 Hz, 3H). $^{13}$C NMR (101 MHz, DMSO) δ 172.69, 169.72, 164.09, 145.86, 139.44, 136.42, 128.62, 128.18, 127.81, 123.80, 121.63, 115.00, 114.64, 70.12, 54.20, 43.59, 31.07, 22.42, 8.57. HRMS (ESI-Q-TOF): calcd. for $C_{21}H_{22}N_3O_4$ [M + H] $^+$ 380.1605, found 380.1604.

**Synthesis of IBA-5**

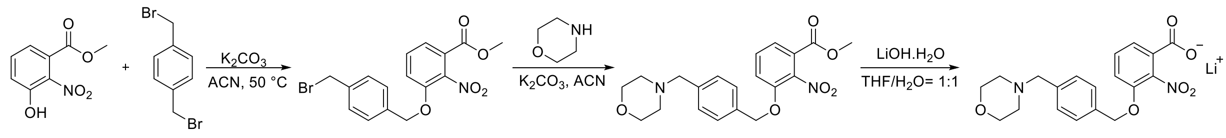

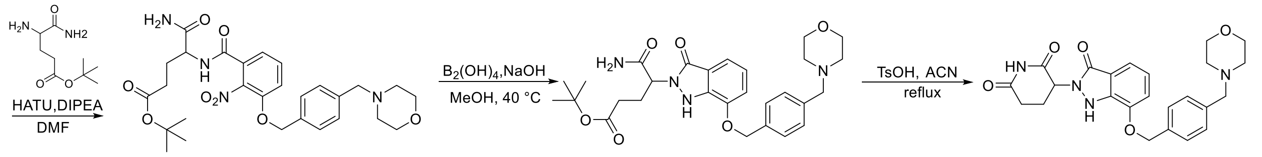

Methyl 3-hydroxy-2-nitrobenzoate (500 mg, 2.54 mmol), 1,4-bis(bromomethyl)benzene (2.0 g, 7.60 mmol), and $K_2CO_3$ (351 mg, 2.54 mmol) were suspended in acetonitrile and heated at 50 °C overnight. After TLC indicated completion, the mixture was cooled to room temperature, filtered, and concentrated under reduced pressure. Purification by column chromatography on silica gel afforded the product as a white solid (910 mg, 94%). $^1$H NMR (500 MHz, $CDCl_3$) δ 7.61 (dd, J = 7.9, 1.1 Hz, 1H), 7.47 – 7.39 (m, 3H), 7.34 (d, J = 8.2 Hz, 2H), 7.24 (dd, J = 8.4, 1.0 Hz, 1H), 5.20 (s, 2H), 4.48 (s, 2H),

3.90 (s, 3H).

Methyl 3-((4-(bromomethyl)benzyl)oxy)-2-nitrobenzoate (350 mg, 0.90 mmol), morpholine (86 μL, 1.01 mmol), and $K_2CO_3$ (191 mg, 1.38 mmol) were suspended in acetonitrile and stirred at room temperature overnight. After TLC indicated completion, the mixture was filtered and concentrated under reduced pressure. Purification by column chromatography afforded the product as a white solid (351 mg, 99%). UPLC–MS (ESI) calcd for $C_{20}H_{23}N_2O_6$ $[M + H]^+$: 387.16; found: 387.18.

Methyl 3-((4-(morpholinomethyl)benzyl)oxy)-2-nitrobenzoate (393 mg, 1.00 mmol) was dissolved in THF (10 mL), and an aqueous solution of $LiOH \cdot H_2O$ (63 mg, 1.5 mmol in 5 mL $H_2O$) was added. The mixture was stirred at room temperature for 2 h. After TLC indicated completion, the reaction mixture was concentrated under reduced pressure, dried under vacuum, and used directly in the next step without purification.

Lithium 3-((4-(morpholinomethyl)benzyl)oxy)-2-nitrobenzoate (201 mg, 0.52 mmol), tert-butyl 4,5-diamino-5-oxopentanoate hydrochloride (124 mg, 0.52 mmol), and HATU (198 mg, 0.52 mmol) were dissolved in anhydrous DMF (10 mL). DIPEA (258 μL, 1.56 mmol) was added, and the reaction mixture was stirred at room temperature overnight. After TLC indicated completion, the reaction was quenched with $H_2O$ and extracted with ethyl acetate. The organic layer was washed with brine, dried over $Na_2SO_4$, filtered, and concentrated under reduced pressure. Purification by column chromatography afforded a gray solid (210 mg, 73%). UPLC–MS (ESI) calcd for $C_{28}H_{37}N_4O_8$ $[M + H]^+$: 557.26; found: 557.62.

Tert-butyl 5-amino-4-(3-((4-(morpholinomethyl)benzyl)oxy)-2-nitrobenzamido)-5-oxopentanoate (202 mg, 0.36 mmol) was dissolved in MeOH (5 mL) and cooled to 0 °C. Tetrahydroxydiboron (161 mg, 1.8 mmol) was added, followed by a solution of NaOH (144 mg, 3.6 mmol) in MeOH (5 mL). The mixture was stirred at 0 °C for 10 min and then at 40 °C overnight. After TLC indicated completion, the reaction was quenched with saturated $NH_4Cl$ solution and extracted with ethyl acetate. The organic layer was washed with $H_2O$ and brine, dried over $Na_2SO_4$, filtered, and concentrated under reduced pressure. Purification by column chromatography afforded the product (160 mg, 85%). $^1H$ NMR (500 MHz, DMSO) δ 9.75 (s, 1H), 7.48 (t, $J$ = 9.4 Hz, 3H), 7.34 (d, $J$ = 7.9 Hz, 2H), 7.30 (s, 1H), 7.23 (d, $J$ = 7.8 Hz, 1H), 7.18 (d, $J$ = 7.9 Hz, 1H), 7.05 (t, $J$ = 7.8 Hz, 1H), 5.22 (s, 2H), 4.87 (dd, $J$ = 9.7, 6.0 Hz, 1H), 3.63 – 3.51 (m, 4H), 3.47 (s, 2H), 2.34 (s, 4H), 2.20 (d, $J$ = 14.5 Hz, 2H), 2.12 – 1.97 (m, 2H), 1.35 (s,

9H). UPLC–MS (ESI) calcd for $C_{28}H_{37}N_4O_6$ $[M + H]^+$: 525.62; found: 525.53.

Tert-butyl 5-amino-4-(7-((4-(morpholinomethyl)benzyl)oxy)-3-oxo-1,3-dihydro-2H-indazol-2-yl)-5-oxopentanoate (75 mg, 0.14 mmol) and *p*-Toluenesulfonic acid monohydrate (160 mg, 0.14 mmol) were dissolved in acetonitrile (6 mL) and stirred at 80 °C overnight. The solvent was removed under reduced pressure, and the residue was purified by preparative HPLC to afford **IBA-5** as a white solid (38 mg, 60%). $^1H$ NMR (500 MHz, DMSO) δ 10.98 (s, 1H), 10.09 (s, 1H), 7.49 (d, J = 8.0 Hz, 2H), 7.34 (d, J = 8.0 Hz, 2H), 7.25 (d, J = 7.7 Hz, 1H), 7.20 (d, J = 7.8 Hz, 1H), 7.05 (t, J = 7.8 Hz, 1H), 5.30 (dd, J = 13.1, 5.2 Hz, 1H), 5.22 (s, 2H), 3.60 – 3.53 (m, 4H), 3.47 (s, 2H), 2.92 (ddd, J = 17.2, 13.8, 5.4 Hz, 1H), 2.65 – 2.58 (m, 1H), 2.57 – 2.51 (m, 1H), 2.34 (s, 4H), 2.10 – 2.03 (m, 1H). $^{13}C$ NMR (126 MHz, DMSO) δ 172.78, 169.63, 165.05, 145.19, 137.75, 135.21, 132.16, 129.00, 127.91, 121.77, 118.55, 116.66, 114.81, 112.95, 69.48, 66.20, 62.14, 53.87, 53.16, 48.61, 30.94, 22.36. HRMS (ESI-Q-TOF): calcd. for $C_{24}H_{27}N_4O_5$ $[M + H]^+$ 451.1976, found 451.1979.

**Synthesis of IBA-6**

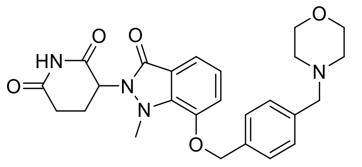

IBA-6

**IBA-6** was prepared following the same procedure as **IBA-3** using tert-butyl 5-amino-4-(7-((4-(morpholinomethyl)benzyl)oxy)-3-oxo-1,3-dihydro-2H-indazol-2-yl)-5-oxopentanoate (300 mg，0.57 mmol) and Iodomethane (43 µL，0.69 mmol) as the starting materials, giving **IBA-6** as a white solid (105 mg, 40% yield over two steps). $^1H$ NMR (500 MHz, DMSO) δ 11.03 (s, 1H), 7.46 (d, J = 8.0 Hz, 2H), 7.38 – 7.31 (m, 3H), 7.27 (d, J = 7.6 Hz, 1H), 7.21 (t, J = 7.8 Hz, 1H), 5.29 – 5.21 (m, 2H), 5.14 (dd, J = 12.4, 5.3 Hz, 1H), 3.57 (t, J = 4.7 Hz, 4H), 3.47 (s, 2H), 3.23 (s, 3H), 2.87 – 2.78 (m, 1H), 2.71 – 2.56 (m, 2H), 2.35 (t, J = 4.4 Hz, 4H), 2.09 – 2.02 (m, 1H). $^{13}C$ NMR (126 MHz, DMSO) δ 172.69, 169.87, 163.27, 146.12, 141.76, 137.83, 135.03, 129.13, 127.78, 124.17, 120.99, 114.97, 114.86, 69.97, 66.17, 62.10, 54.07, 53.16, 31.14, 22.68. HRMS (ESI-Q-TOF): calcd. for $C_{25}H_{29}N_4O_5$ $[M + H]^+$ 465.2132, found 465.2129.

**Synthesis of IBA-7**

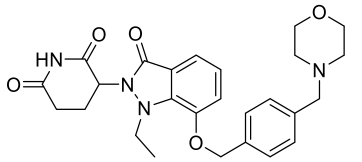

IBA-7

**IBA-7** was prepared following the same procedure as **IBA-4**, using tert-butyl 5-amino-4-(7-((4-(morpholinomethyl)benzyl)oxy)-3-oxo-1,3-dihydro-2H-indazol-2-yl)-5-oxopentanoate (300 mg， 0.57 mmol), Iodoethane (55 µL， 0.69 mmol) as the starting materials, giving **IBA-7** as a white solid (65 mg, 24% yield over two steps). $^1$H NMR (500 MHz, DMSO) δ 11.02 (s, 1H), 7.45 (d, J = 7.8 Hz, 2H), 7.36 (d, J = 7.7 Hz, 2H), 7.32 (d, J = 7.9 Hz, 1H), 7.26 (d, J = 7.6 Hz, 1H), 7.18 (t, J = 7.8 Hz, 1H), 5.26 (s, 2H), 5.07 (dd, J = 12.3, 5.3 Hz, 1H), 4.09 (dq, J = 14.0, 6.8 Hz, 1H), 3.76 (dq, J = 14.0, 6.7 Hz, 1H), 3.57 (t, J = 4.6 Hz, 4H), 3.48 (s, 2H), 2.85 – 2.75 (m, 1H), 2.71 – 2.56 (m, 2H), 2.35 (d, J = 4.9 Hz, 4H), 2.02 (ddd, J = 10.5, 5.5, 3.0 Hz, 1H), 0.69 (t, J = 6.9 Hz, 3H). $^{13}$C NMR (126 MHz, DMSO) δ 172.71, 169.73, 164.09, 145.88, 139.45, 135.12, 129.17, 127.74, 123.81, 121.64, 114.99, 114.65, 69.95, 66.16, 62.09, 54.18, 53.15, 43.59, 31.07, 22.41, 8.55. HRMS (ESI-Q-TOF): calcd. for $C_{26}H_{31}N_4O_5$ [M + H]$^+$ 479.2289, found 479.2289.

**Synthesis of IBA-8**

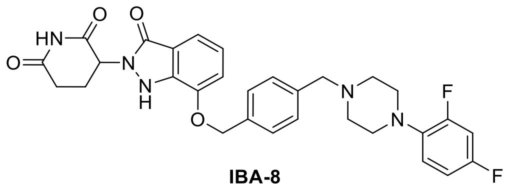

IBA-8

**IBA-8** was prepared following the same procedure as **IBA-5**, using methyl 3-hydroxy-2-nitrobenzoate and 1,4-bis(bromomethyl)benzene as the starting materials, giving **IBA-8** as a white solid. $^1$H NMR (500 MHz, DMSO) δ 10.98 (s, 1H), 10.11 (s, 1H), 7.51 (d, J = 7.6 Hz, 2H), 7.37 (d, J = 7.6 Hz, 2H), 7.25 (d, J = 7.8 Hz, 1H), 7.21 (d, J = 7.9 Hz, 1H), 7.18 – 7.14 (m, 1H), 7.06 (td, J = 9.7, 8.7, 5.4 Hz, 2H), 6.98 (td, J = 8.6, 2.9 Hz, 1H), 5.31 (dd, J = 13.1, 5.2 Hz, 1H), 5.23 (s, 2H), 3.54 (s, 2H), 3.03 – 2.87 (m, 5H), 2.65 – 2.50 (m, 6H), 2.07 (ddd, J = 9.9, 5.6, 2.7 Hz, 1H). $^{13}$C NMR (126 MHz, DMSO) δ 172.78, 169.63, 162.79, 145.20, 137.77, 129.03, 127.94, 121.80, 120.02, 118.58, 114.82, 112.96, 111.07, 110.89, 104.59, 69.49, 61.68, 53.86, 52.55, 50.46, 30.93, 22.35. HRMS (ESI-Q-TOF): calcd. for $C_{30}H_{30}F_2N_5O_4$ [M + H]$^+$ 562.2260, found 562.2260.

**Synthesis of IBA-9**

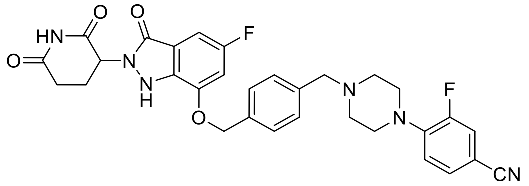

**IBA-9**

**IBA-9** was prepared following the same procedure as **IBA-5**, using methyl 5-fluoro-3-hydroxy-2-nitrobenzoate and 1,4-bis(bromomethyl)benzene as the starting materials, giving **IBA-9** as a white solid. $^1$H NMR (400 MHz, DMSO) δ 11.00 (s, 1H), 10.15 (s, 1H), 7.78 (dd, *J* = 13.1, 1.9 Hz, 1H), 7.69 – 7.61 (m, 3H), 7.58 (d, *J* = 7.9 Hz, 2H), 7.26 – 7.18 (m, 2H), 7.04 (dd, *J* = 7.7, 2.2 Hz, 1H), 5.32 (d, *J* = 6.5 Hz, 3H), 4.43 (s, 2H), 3.72 (s, 2H), 3.51 – 3.07 (m, 6H), 2.93 (ddd, *J* = 17.1, 13.7, 5.4 Hz, 1H), 2.66-2.58 (m, 1H), 2.56-2.51 (m, 1H), 2.16-2.02 (m, 1H). $^{13}$C NMR (101 MHz, DMSO) δ 172.73, 169.49, 166.12, 156.57, 154.45, 152.00, 145.94, 142.31, 142.23, 137.65, 134.75, 131.54, 129.98, 128.30, 120.11, 120.01, 119.86, 118.13, 103.74, 103.64, 69.65, 58.50, 54.08, 50.37, 46.21, 30.89, 22.30. HRMS (ESI-Q-TOF): calcd. for $C_{31}H_{29}F_2N_6O_4$ [M + H] $^+$ 587.2213, found 587.2210.

**Synthesis of IBA-10**

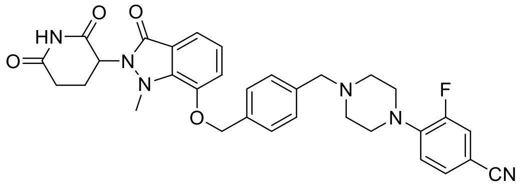

**IBA-10**

**IBA-10** was prepared following the same procedure as **IBA-6**, using tert-butyl 5-amino-4-(7-((4-((4-(4-cyano-2-fluorophenyl)piperazin-1-yl)methyl)benzyl)oxy)-3-oxo-1,3-dihydro-2H-indazol-2-yl)-5-oxopentanoate (331 mg，0.51 mmol) and Iodomethane (39 μL，0.62 mmol) as the starting materials, giving **IBA-10** as a white solid (77 mg, 38% yield over two steps). $^1$H NMR (400 MHz, DMSO) δ 11.04 (s, 1H), 7.79 (dd, *J* = 13.1, 1.9 Hz, 1H), 7.64 (dd, *J* = 8.5, 2.2 Hz, 3H), 7.58 (d, *J* = 8.0 Hz, 2H), 7.34 (d, *J* = 7.8 Hz, 1H), 7.29 (d, *J* = 7.6 Hz, 1H), 7.22 (dt, *J* = 8.7, 4.8 Hz, 2H), 5.35 (s, 2H), 5.13 (dd, *J* = 12.5, 5.3 Hz, 1H), 4.43 (s, 2H), 3.72 (d, *J* = 12.1 Hz, 2H), 3.49 – 3.39 (m, 2H), 3.25 (s, 5H), 3.17 (d, *J* = 12.5 Hz, 2H), 2.87 – 2.76 (m, 1H), 2.70 – 2.58 (m, 2H), 2.11 – 2.02 (m, 1H). $^{13}$C NMR (126 MHz, DMSO) δ 172.76, 169.94, 163.32, 154.26, 152.29, 146.02, 142.35, 142.28, 141.80, 138.10, 131.74, 130.04, 129.27, 128.23, 124.25, 121.14, 120.05, 118.19, 116.94, 115.10, 114.62, 103.78, 69.61, 58.58, 54.16, 50.44, 46.28, 31.19, 22.75. HRMS (ESI-Q-TOF): calcd. for $C_{32}H_{32}FN_6O_4$ [M + H] $^+$ 583.2464, found 583.2462.

#### Synthesis of IBA-11

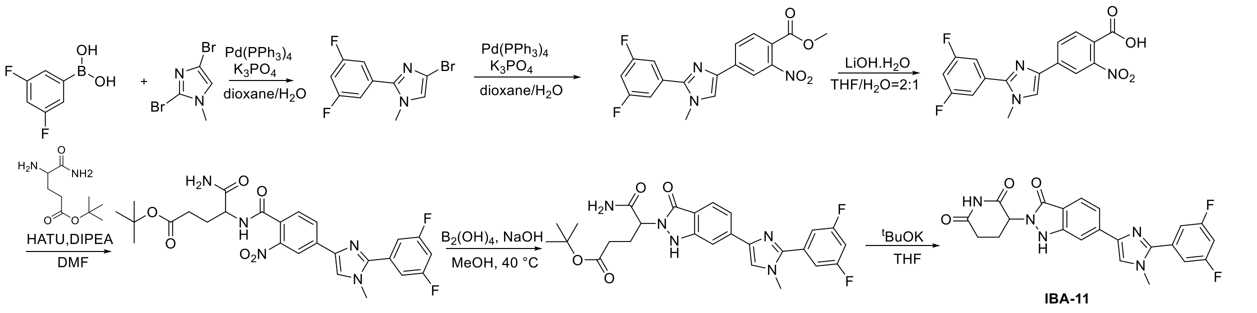

A mixture of compound (3,5-difluorophenyl)boronic acid (186 mg, 1.18 mmol), 2,4-dibromo-1-methyl-1H-imidazole (280 mg, 1.18 mmol), and $Pd(PPh_3)_4$ (136 mg, 0.118 mmol) was dissolved in 1,4-dioxane (12 mL). An aqueous solution of $K_3PO_4$ (751 mg, 3.54 mmol in 4 mL $H_2O$) was added, and the reaction mixture was degassed with nitrogen three times and stirred at 80 °C under nitrogen for 8 h. After LC-MS indicated completion, the mixture was diluted with ethyl acetate, washed with $H_2O$ and brine, dried over $Na_2SO_4$, filtered, and concentrated under reduced pressure. Purification by column chromatography on silica gel afforded the product as a colorless oil (220 mg, 68%). $^1H$ NMR (500 MHz, $CDCl_3$) δ 7.19 (d, J = 6.7 Hz, 2H), 6.97 (s, 1H), 6.88 (s, 1H), 3.77 (s, 3H).

A mixture of 4-bromo-2-(3,5-difluorophenyl)-1-methyl-1H-imidazole (220 mg, 0.81 mmol), (4-(methoxycarbonyl)-3-nitrophenyl)boronic acid (249 mg, 0.81 mmol), and $Pd(PPh_3)_4$ (94 mg, 0.08 mmol) in 1,4-dioxane (12 mL) was treated with an aqueous solution of $K_3PO_4$ (516 mg, 2.43 mmol in 4 mL $H_2O$). The reaction mixture was degassed with nitrogen three times and stirred at 80 °C under nitrogen overnight. After LC-MS indicated completion, the mixture was diluted with ethyl acetate, washed with $H_2O$ and brine, dried over $Na_2SO_4$, filtered, and concentrated under reduced pressure. Purification by column chromatography afforded the product as a yellow solid (218 mg, 72%). UPLC–MS (ESI) calcd for $C_{18}H_{14}F_2N_3O_4$ $[M + H]^+$: 374.10; found: 374.28.

Methyl 4-(2-(3,5-difluorophenyl)-1-methyl-1H-imidazol-4-yl)-2-nitrobenzoate (218 mg, 0.58 mmol) was dissolved in THF (6 mL), and an aqueous solution of $LiOH·H_2O$ (49 mg, 1.17 mmol in 3 mL $H_2O$) was added. The mixture was stirred at room temperature for 2 h. After TLC indicated completion, the reaction mixture was concentrated under reduced pressure, dried under vacuum, and used directly in the next step without purification.

Lithium 4-(2-(3,5-difluorophenyl)-1-methyl-1H-imidazol-4-yl)-2-nitrobenzoate (0.58 mmol), tert-butyl 4,5-diamino-5-oxopentanoate (138 mg, 0.58 mmol), and HATU (221 mg, 0.58 mmol) were dissolved in anhydrous DMF (5 mL). DIPEA (303 μL, 1.74 mmol) was added, and the reaction mixture was stirred at room temperature overnight. After TLC indicated completion, the reaction was quenched with $H_2O$ and extracted with ethyl acetate. The organic layer was washed with brine, dried over $Na_2SO_4$, filtered, and concentrated under reduced pressure. Purification by column chromatography afforded the product as a gray solid (80 mg, 26%). UPLC–MS (ESI) calcd for $C_{26}H_{28}F_2N_5O_6$ $[M + H]^+$: 544.20; found: 544.07.

Tert-butyl 5-amino-4-(4-(2-(3,5-difluorophenyl)-1-methyl-1H-imidazol-4-yl)-2-nitrobenzamido)-5-oxopentanoate (80 mg, 0.15 mmol) was dissolved in MeOH (2 mL) and cooled to 0 °C. Tetrahydroxydiboron (67 mg, 0.75 mmol) was added, followed by a solution of NaOH (60 mg, 1.5 mmol) in MeOH (2 mL). The mixture was stirred at 0 °C for 10 min and then at 40 °C overnight. After TLC indicated completion, the reaction was quenched with saturated $NH_4Cl$ solution and extracted with ethyl acetate. The organic layer was washed with $H_2O$ and brine, dried over $Na_2SO_4$, filtered, and concentrated under reduced pressure. Purification by column chromatography afforded the product as a white solid (51 mg, 66%). UPLC–MS (ESI) calcd for $[M + H]^+$: 512.21; found:512.23.

Tert-butyl 5-amino-4-(6-(2-(3,5-difluorophenyl)-1-methyl-1H-imidazol-4-yl)-3-oxo-1,3-dihydro-2H-indazol-2-yl)-5-oxopentanoate (51 mg, 0.12 mmol) was dissolved in anhydrous THF (2 mL) and cooled to 0 °C. A solution of potassium tert-butoxide (1 M in THF, 240 μL, 0.24 mmol) was added dropwise, and the reaction mixture was stirred at 0 °C for 1 h. After LC–MS indicated completion, the reaction was quenched with 1M HCl (240 μL), diluted with ethyl acetate, and washed with $H_2O$ and brine. The organic layer was dried over $Na_2SO_4$, filtered, and concentrated. The residue was purified by preparative HPLC to afford **IBA-11** as a white solid (38 mg, 60%). $^1H$ NMR (400 MHz, DMSO) δ 11.03 (s, 1H), 9.92 (s, 1H), 7.98 (s, 1H), 7.68 (d, J = 8.2 Hz, 1H), 7.63 (s, 1H), 7.60 – 7.56 (m, 1H), 7.51 (d, J = 6.3 Hz, 2H), 7.42 – 7.32 (m, 1H), 5.33 (dd, J = 13.1, 5.2 Hz, 1H), 3.86 (s, 3H), 2.94 (ddd, J = 18.1, 13.6, 5.5 Hz, 1H), 2.71 – 2.58 (m, 1H), 2.39 (td, J = 13.3, 4.5 Hz, 1H), 2.15 – 2.05 (m, 1H). $^{13}C$ NMR (101 MHz, DMSO) δ 172.75, 169.84, 163.25, 161.20, 161.07, 148.43, 139.07, 137.79, 123.45, 122.30, 118.31, 115.81, 111.49, 111.23, 107.17, 53.98, 34.84, 30.91, 22.47. HRMS (ESI-Q-TOF): calcd. for $C_{22}H_{18}F_2N_5O_3$ $[M + H]^+$ 438.1372, found 438.1373.

**Synthesis of IBA-12**

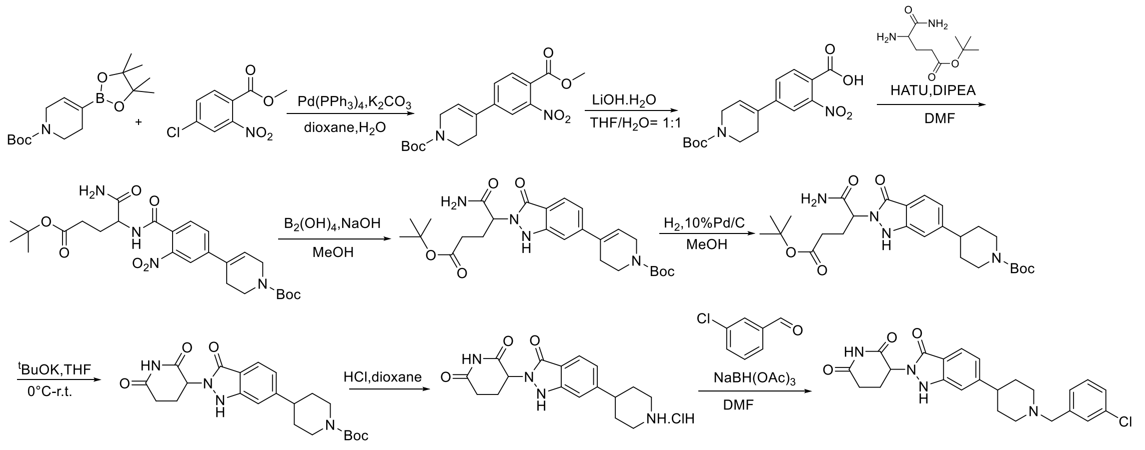

**IBA-12**

Methyl 4-chloro-2-nitrobenzoate (1.0 g, 4.63 mmol), tert-butyl 4-(4,4,5,5-tetramethyl-1,3,2-dioxaborolan-2-yl)-3,6-dihydropyridine-1(2H)-carboxylate (1.43 g, 4.63 mmol), $K_2CO_3$ (1.28 g, 9.26 mmol), and $Pd(PPh_3)_4$ (268 mg, 0.232 mmol) were added to a mixture of 1,4-dioxane (20 mL) and water (5 mL). The reaction mixture was degassed with nitrogen three times and stirred at 90 °C overnight under a nitrogen atmosphere. After completion of the reaction, the solvent was removed under reduced pressure to remove 1,4-dioxane. The residue was extracted with ethyl acetate, and the combined organic layers were washed with brine, dried over $Na_2SO_4$, filtered, and concentrated under reduced pressure. Purification by silica gel column chromatography afforded the product as a yellow oil (1.5 g, 89%).$^1$H NMR (500 MHz, DMSO) δ 8.03 (t, J = 1.1 Hz, 1H), 7.85 (d, J = 1.2 Hz, 2H), 6.48 (s, 1H), 4.08 – 3.99 (m, 2H), 3.84 (s, 3H), 3.54 (t, J = 5.7 Hz, 2H), 2.52 – 2.48(m, 2H), 1.42 (s, 9H).

Tert-Butyl 4-(4-(methoxycarbonyl)-3-nitrophenyl)-3,6-dihydropyridine-1(2H)-carboxylate (1.0 g, 2.76 mmol) was dissolved in THF (10 mL). An aqueous solution of $LiOH·H_2O$ (174 mg, 4.14 mmol) in water (10 mL) was added, and the reaction mixture was stirred at room temperature overnight. The reaction mixture was concentrated under reduced pressure to give the crude product (977 mg), which was used in the next step without further purification.

Lithium 4-(1-(tert-butoxycarbonyl)-1,2,3,6-tetrahydropyridin-4-yl)-2-nitrobenzoate (977 mg, 2.76 mmol), tert-butyl 4,5-diamino-5-oxopentanoate hydrochloride (659 mg, 2.76 mmol), and HATU (1.05 g, 2.76 mmol) were dissolved in DMF (15 mL). DIPEA (1.4 mL, 8.28 mmol) was added, and the reaction mixture was stirred at room temperature for 12 h. After TLC indicated completion, the reaction mixture was diluted with ethyl acetate and washed with $H_2O$ and saturated NaCl solution. The organic layer was dried over $Na_2SO_4$, filtered, and concentrated under reduced pressure. Purification by silica gel column chromatography afforded the product as a yellow oil (1.28 g, 87%). UPLC-MS (ESI): calcd for $C_{26}H_{37}N_4O_8$ $[M + H]^+$ 533.59, found 533.26.

Tert-Butyl 4-(4-((1-amino-5-(tert-butoxy)-1,5-dioxopentan-2-yl)carbamoyl)-3-nitrophenyl)-3,6-dihydropyridine-1(2H)-carboxylate (1.78 g, 3.35 mmol) was dissolved in methanol (20 mL). Tetrahydroxydiboron (1.5 g, 16.76 mmol) was added at 0 °C, followed by a solution of NaOH (1.34 g, 33.5 mmol) in methanol (20 mL). The reaction mixture was stirred at 0 °C for 10 min and then heated to 40 °C and stirred overnight. After TLC indicated completion, the reaction was quenched with saturated $NH_4Cl$ solution and extracted with ethyl acetate. The organic phase was washed with water and brine, dried over $Na_2SO_4$, filtered, and concentrated under reduced pressure. Purification by silica gel column chromatography afforded the product as a yellow solid (1.1 g, 66%). UPLC-MS (ESI): calcd for $C_{26}H_{37}N_4O_6$ $[M + H]^+$ 501.27, found 501.35.

A mixture of tert-butyl 4-(2-(1-amino-5-(tert-butoxy)-1,5-dioxopentan-2-yl)-3-oxo-2,3-dihydro-1H-indazol-6-yl)-3,6-dihydropyridine-1(2H)-carboxylate (100 mg, 0.2 mmol) and 10% Pd/C (10 mg) in methanol (10 mL) was degassed was degassed with nitrogen three times and then with hydrogen three times. The reaction mixture stirred under a hydrogen atmosphere (1atm) at room temperature overnight. After completion, the reaction mixture was filtered through Celite, and the filtrate was concentrated under reduced pressure. The residue was purified by silica gel column chromatography to afford the product as a yellow solid (73 mg, 73%). UPLC-MS (ESI): calcd for $C_{26}H_{39}N_4O_6$ $[M + H]^+$ 503.29, found 503.47.

Tert-Butyl 4-(2-(1-amino-5-(tert-butoxy)-1,5-dioxopentan-2-yl)-3-oxo-2,3-dihydro-1H-indazol-6-yl)piperidine-1-carboxylate (400 mg, 0.80 mmol) was dissolved in THF. A 1 M solution of potassium tert-butoxide in THF (1.6 mL) was added at 0 °C. The reaction mixture was allowed to warm to room temperature and stirred for 1 h. The mixture was neutralized to pH 7 with 1 M HCl, diluted with $H_2O$, and extracted with ethyl acetate. The organic layer was washed with brine, dried over $Na_2SO_4$, filtered, and concentrated under reduced pressure. Purification by silica gel column chromatography afforded a yellow solid (216 mg, 63%). $^1$H NMR (400 MHz, DMSO) δ 10.98 (s, 1H), 9.36 (s, 1H), 7.59 (d, J = 8.1 Hz, 1H), 7.13 – 6.98 (m, 2H), 5.31 (dd, J = 13.1, 5.2 Hz, 1H), 4.08 (d, J = 12.8 Hz, 2H), 3.00 – 2.70 (m, 4H), 2.62 (ddd, J = 17.4, 4.4, 2.4 Hz, 1H), 2.40 (qd, J = 13.1, 4.5 Hz, 1H), 2.06 (tt, J = 7.8, 5.1 Hz, 1H), 1.81 – 1.73 (m, 2H), 1.51 (qd, J = 12.8, 4.6 Hz, 2H), 1.42 (s, 9H).

Tert-Butyl 4-(2-(1-amino-5-(tert-butoxy)-1,5-dioxopentan-2-yl)-3-oxo-2,3-dihydro-1H-indazol-6-

yl)piperidine-1-carboxylate (216 mg, 0.50 mmol) was dissolved in dichloromethane (4 mL). A 4 M solution of HCl in 1,4-dioxane (4 mL) was added, and the mixture was stirred at room temperature overnight. The reaction mixture was concentrated under reduced pressure, and the crude product was used directly in the next step without purification. UPLC-MS (ESI): calcd for $C_{17}H_{21}N_4O_3$ $[M + H]^+$ 329.16, found 329.35.

3-(3-oxo-6-(piperidin-4-yl)-1,3-dihydro-2H-indazol-2-yl)piperidine-2,6-dione hydrochloride (73 mg, 0.2 mmol) was dissolved in a mixture of DMF (1 mL) and dichloromethane (1 mL). m-Chlorobenzaldehyde (0.025 mL, 0.25 mmol) and Sodium triacetoxyborohydride (127 mg, 0.6 mmol) were added. The reaction mixture was stirred at room temperature overnight. After completion, the reaction mixture was diluted with ethyl acetate and washed with saturated NaCl solution. The organic layer was dried over $Na_2SO_4$, filtered, and concentrated under reduced pressure. Purification by silica gel column chromatography afforded **IBA-12** (30 mg, 35%). $^1$H NMR (500 MHz, DMSO) δ 11.02 (s, 1H), 9.96 (s, 1H), 7.68 (s, 1H), 7.64 (d, J = 8.1 Hz, 1H), 7.58 – 7.50 (m, 3H), 7.05 – 6.99 (m, 2H), 5.32 (dd, J = 13.1, 5.2 Hz, 1H), 4.38 (d, J = 3.7 Hz, 2H), 3.49 (d, J = 11.9 Hz, 2H), 3.09 (q, J = 11.3 Hz, 2H), 2.99-2.87 (m, 2H), 2.69 – 2.59 (m, 1H), 2.38 (qd, J = 13.2, 4.5 Hz, 1H), 2.11 – 2.00 (m, 3H), 1.92 (qd, J = 13.0, 3.8 Hz, 2H). $^{13}$C NMR (126 MHz, DMSO) δ 172.77, 169.81, 163.06, 148.71, 148.04, 133.43, 132.04, 131.18, 130.75, 130.17, 129.64, 123.48, 120.49, 116.08, 110.00, 58.40, 53.93, 51.82, 30.91, 29.75, 29.70, 22.46. HRMS (ESI-Q-TOF): calcd. for $C_{24}H_{26}ClN_4O_3$ $[M + H]^+$ 453.1688, found 453.1688.

#### The $^{1}$H NMR and $^{13}$C NMR spectra

$^{1}$H NMR Spectrum of Compound **IBA-1**

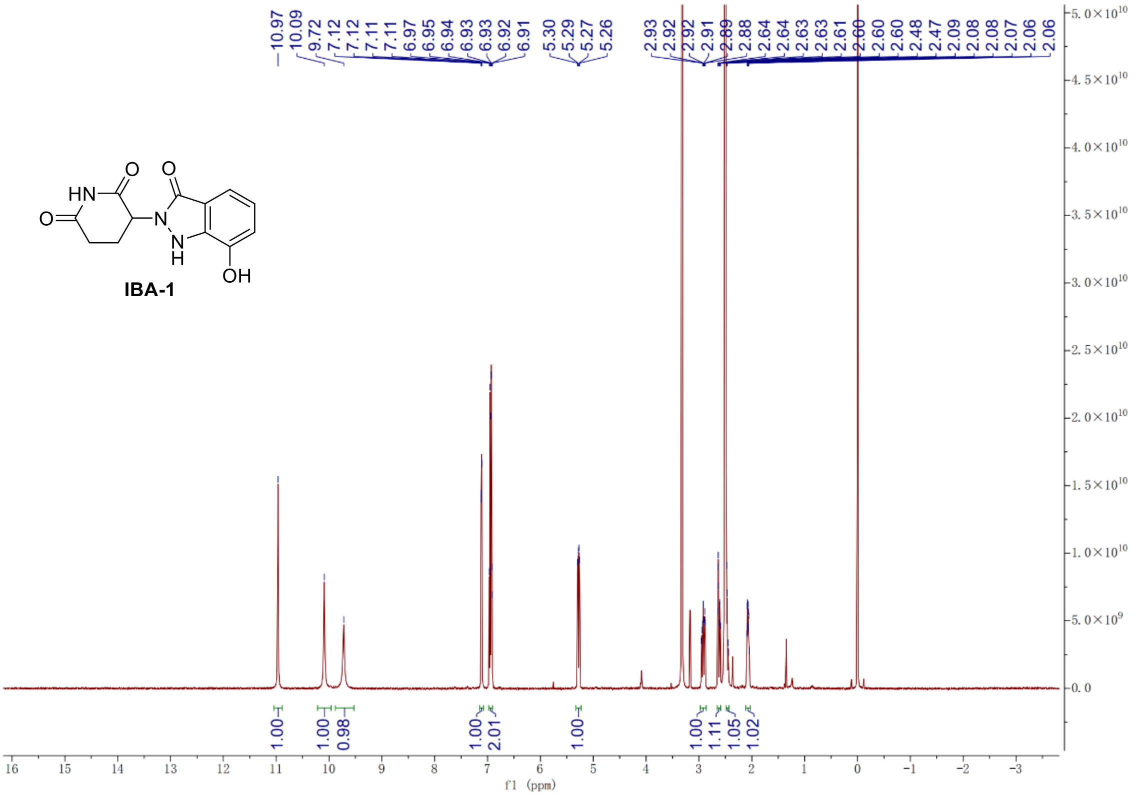

$^{13}$C NMR Spectrum of Compound **IBA-1**

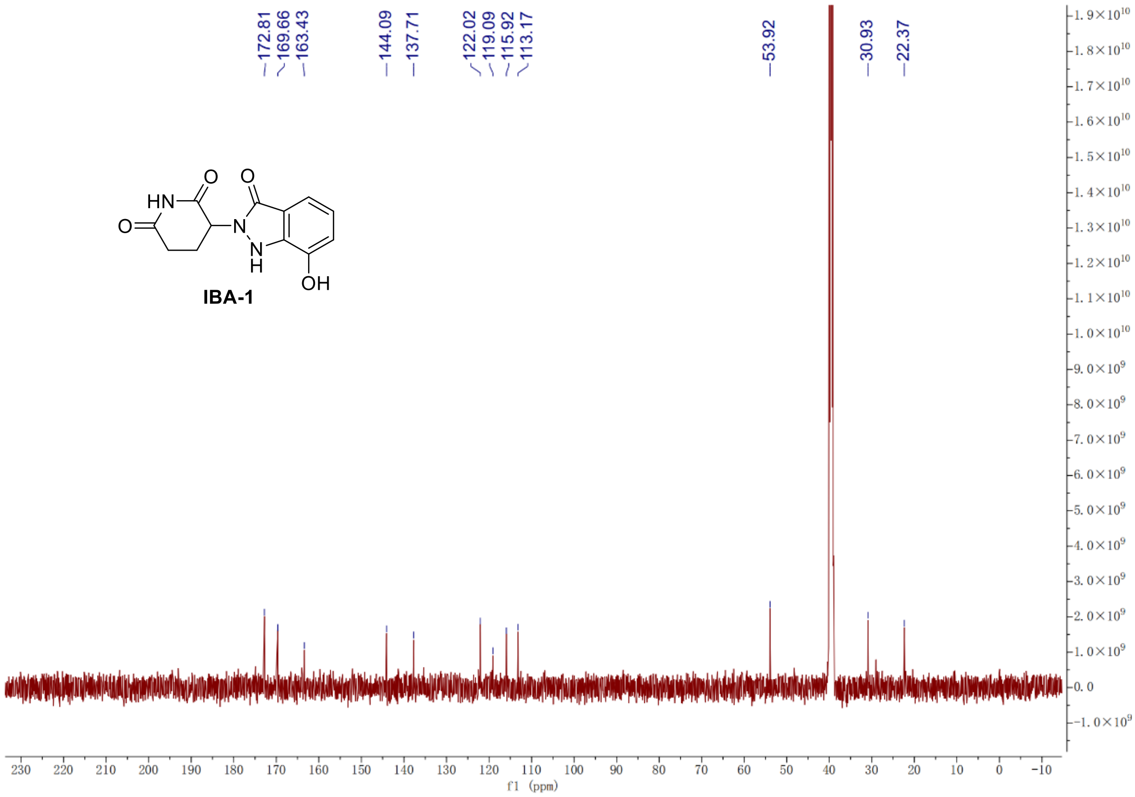

#### [1]H NMR Spectrum of Compound **IBA-2**

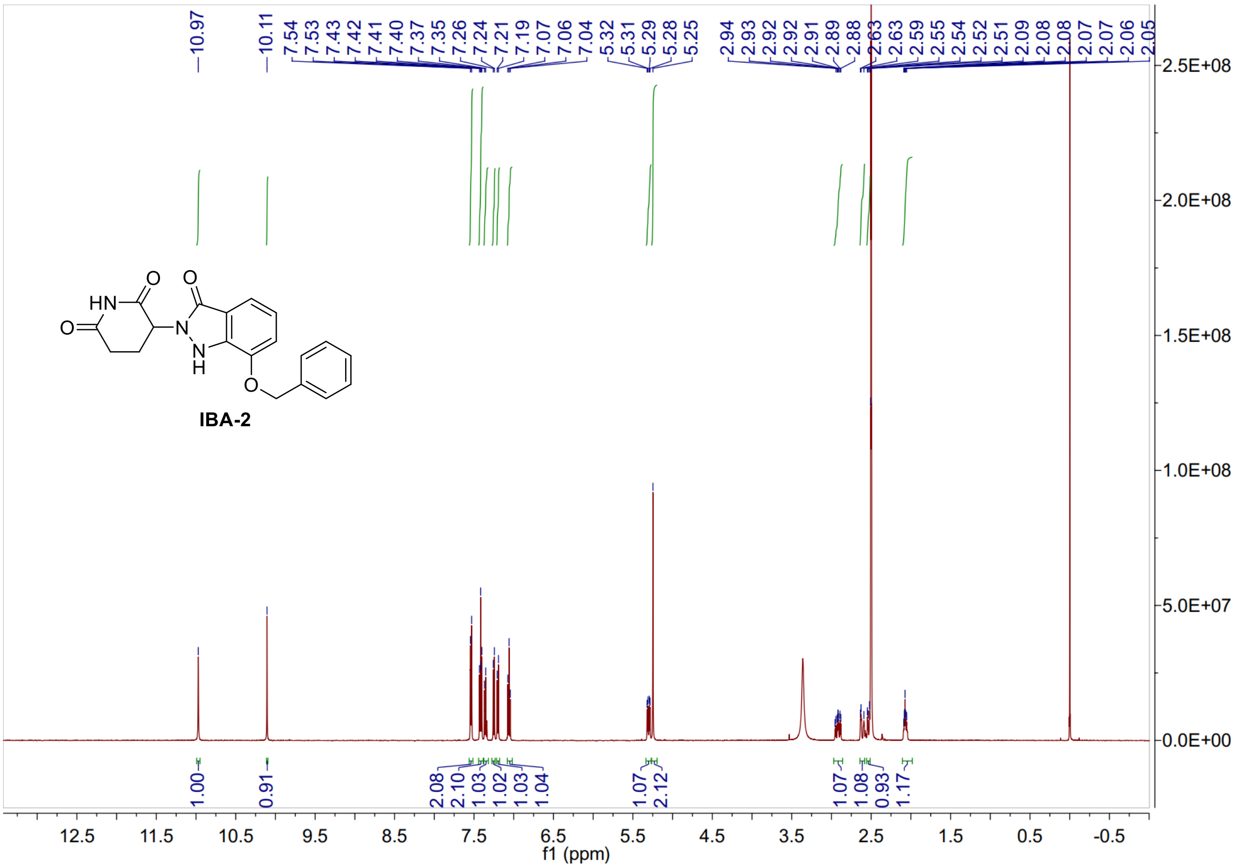

#### [13]C NMR Spectrum of Compound **IBA-2**

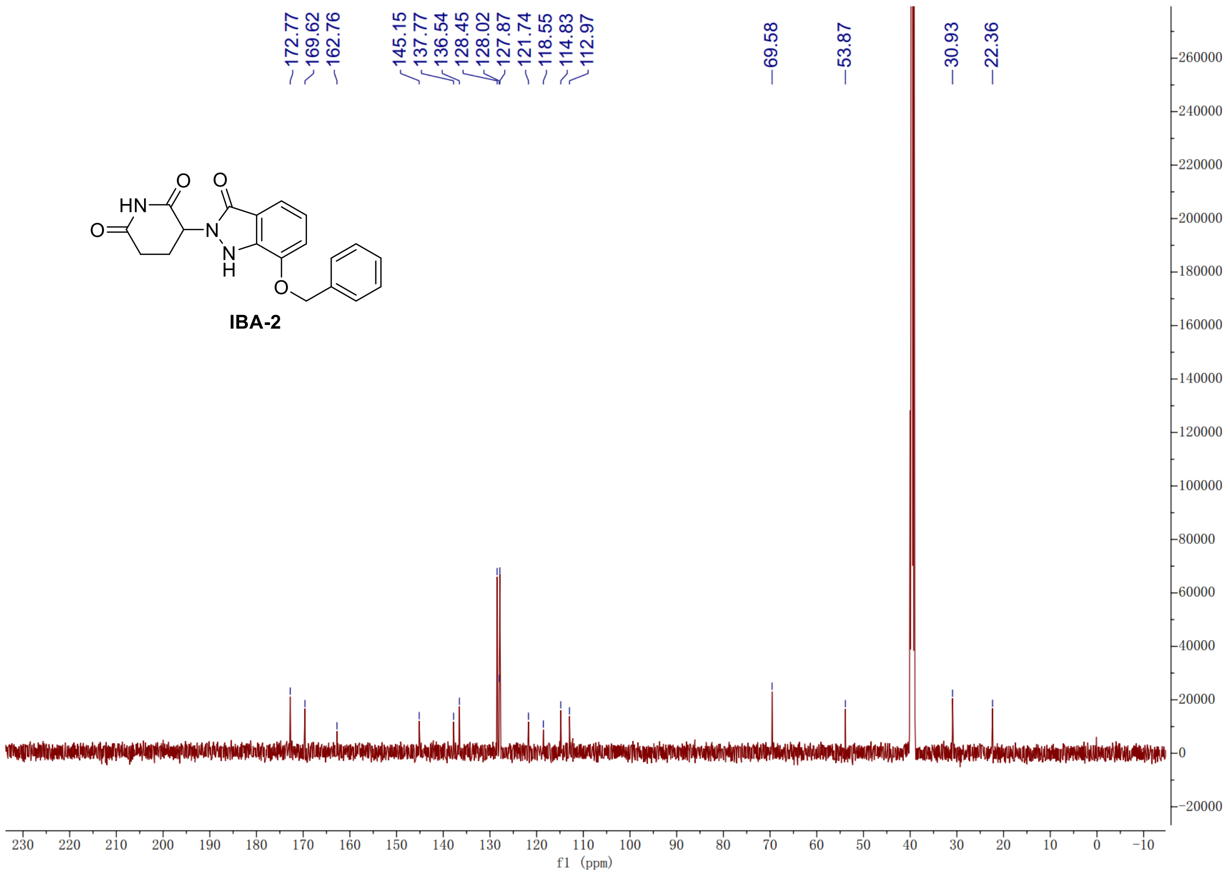

#### $^{1}H$ NMR Spectrum of Compound **IBA-3**

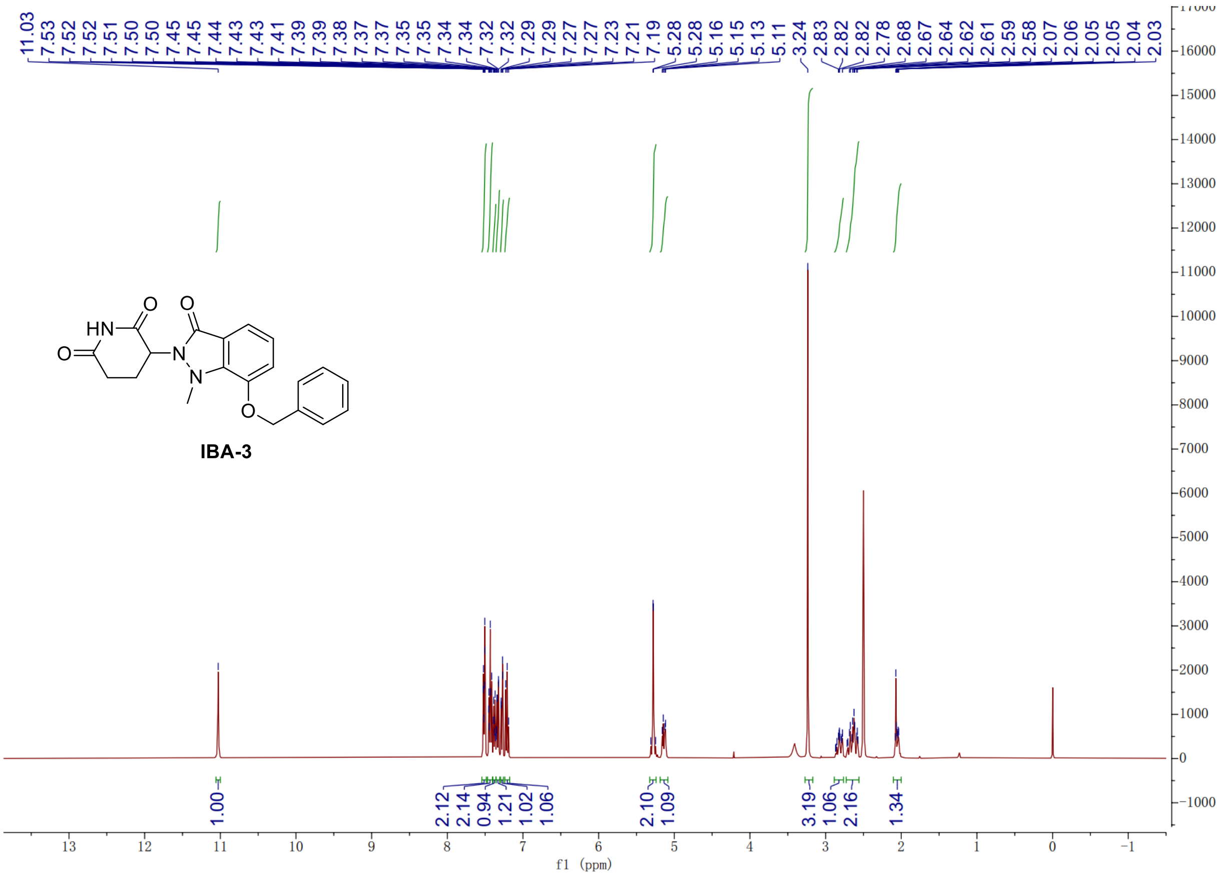

#### $^{13}C$ NMR Spectrum of Compound **IBA-3**

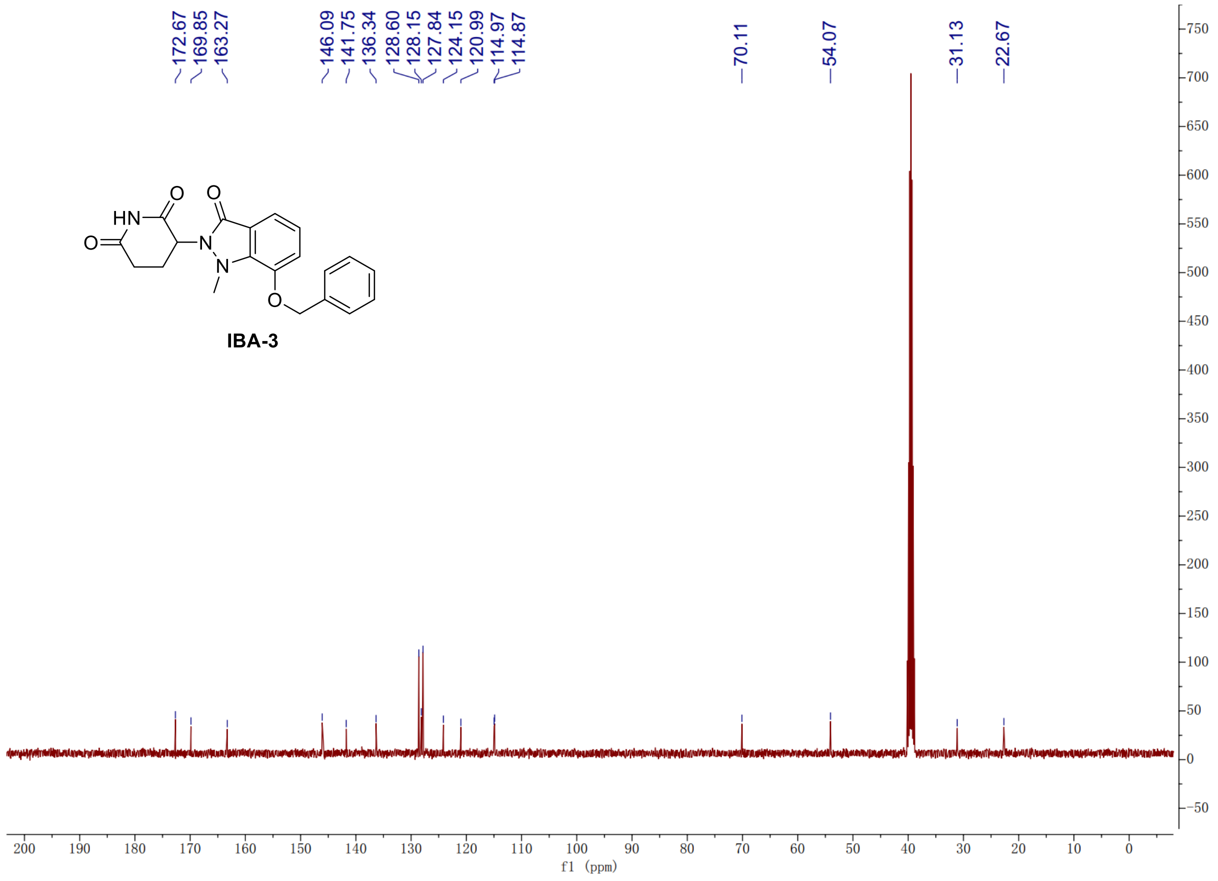

#### $^{1}$H NMR Spectrum of Compound **IBA-4**

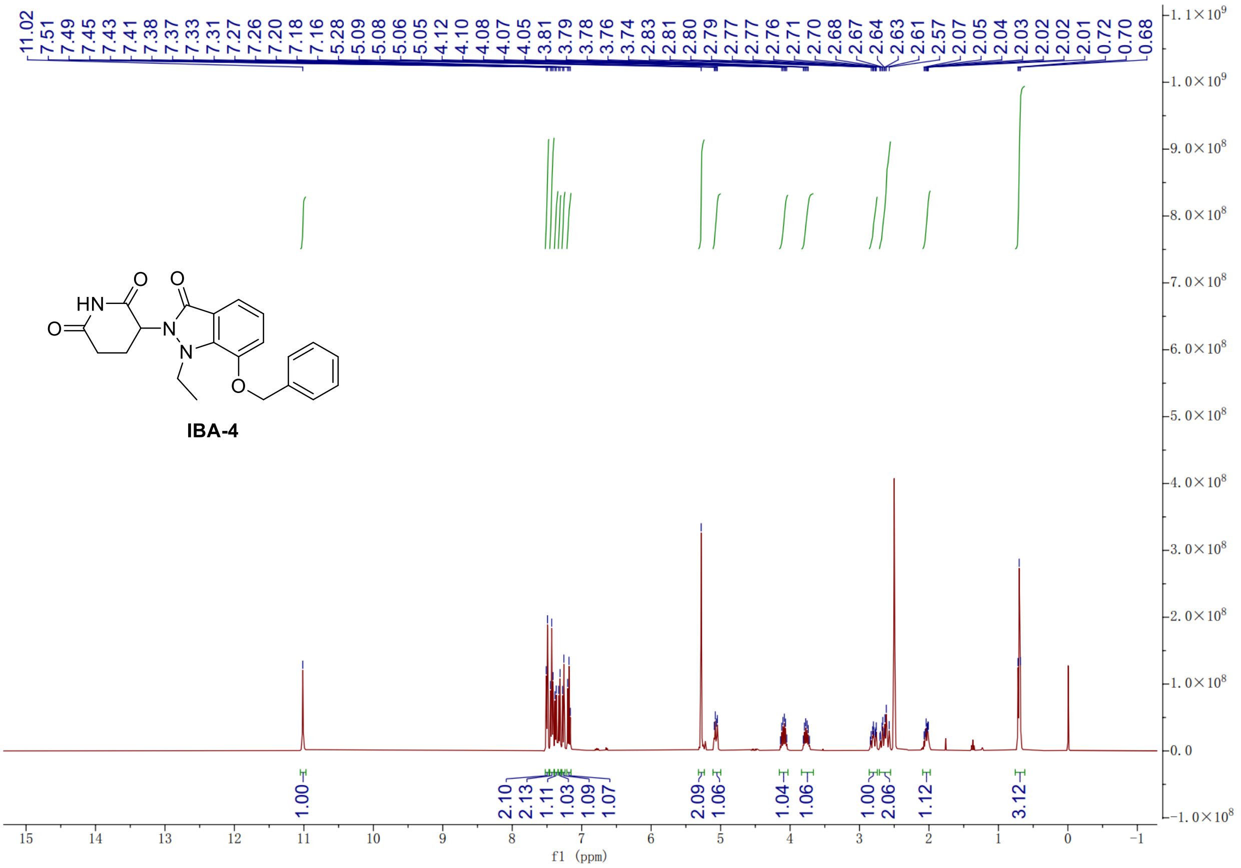

#### $^{13}$C NMR Spectrum of Compound **IBA-4**

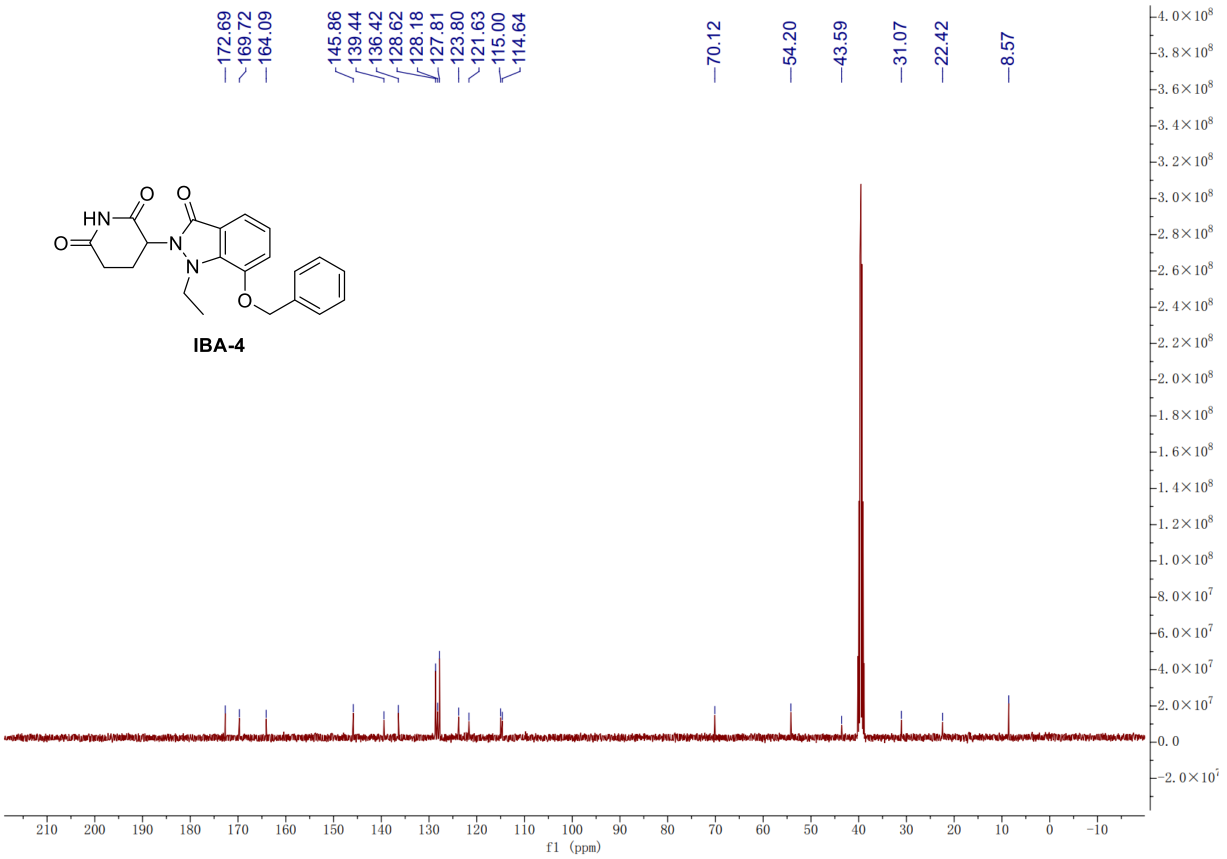

#### $^{1}$H NMR Spectrum of Compound **IBA-5**

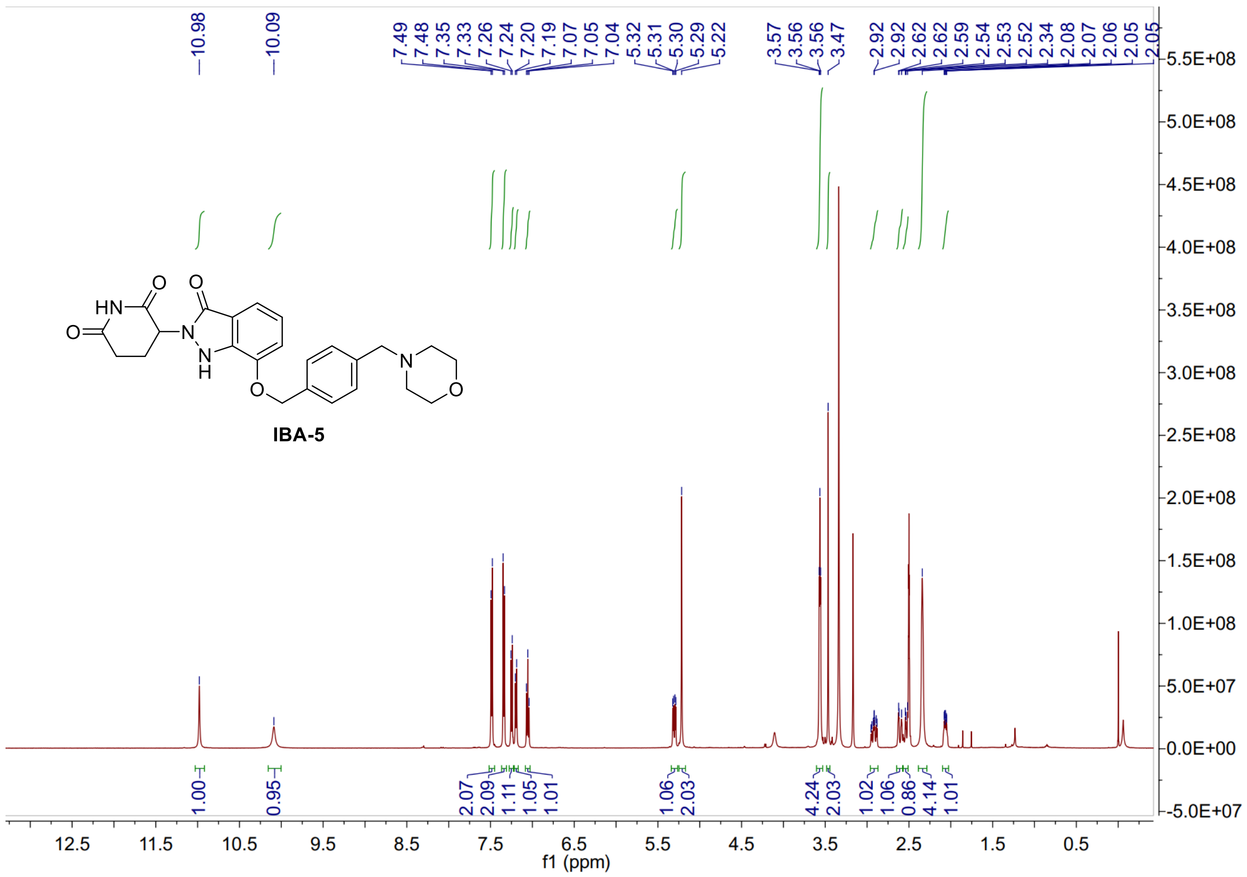

#### $^{13}$C NMR Spectrum of Compound **IBA-5**

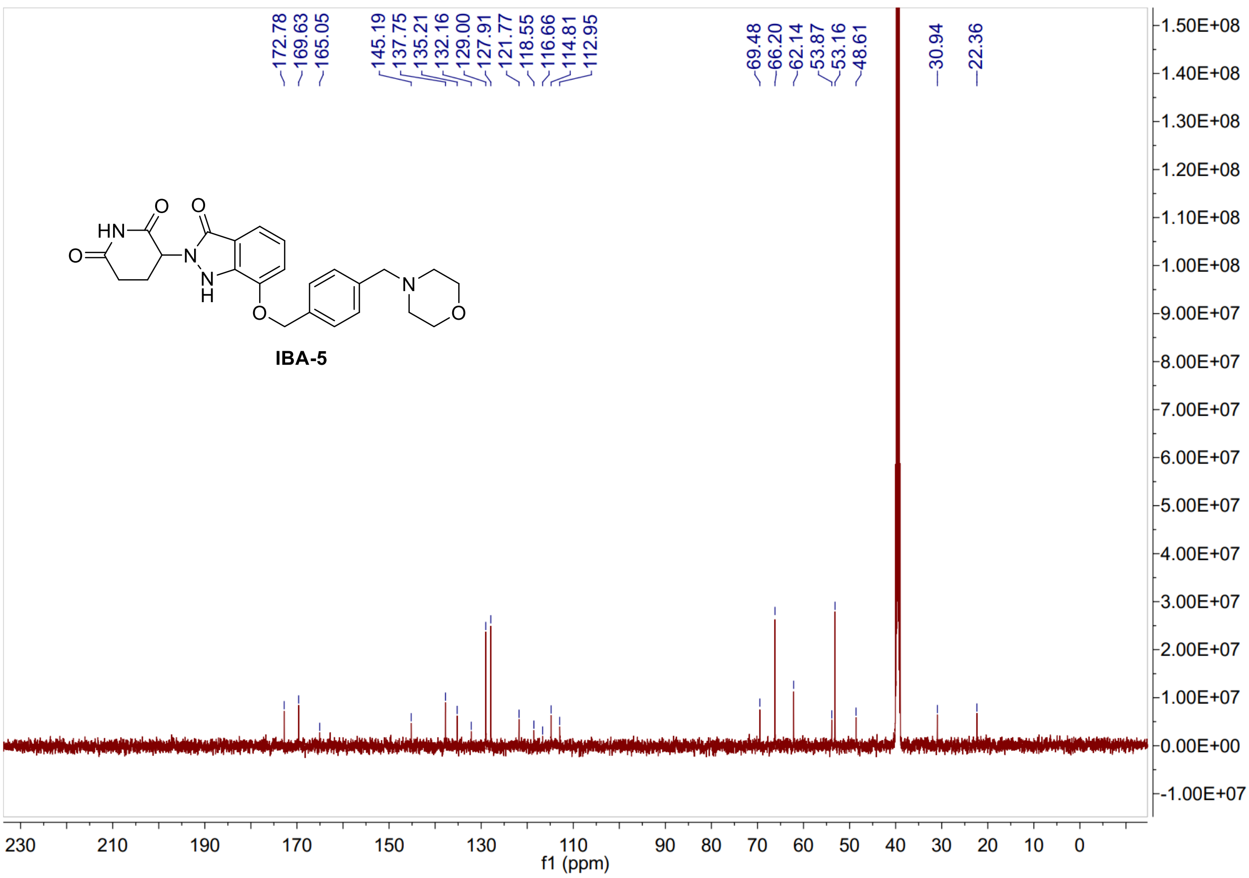

### $^{1}$H NMR Spectrum of Compound **IBA-6**

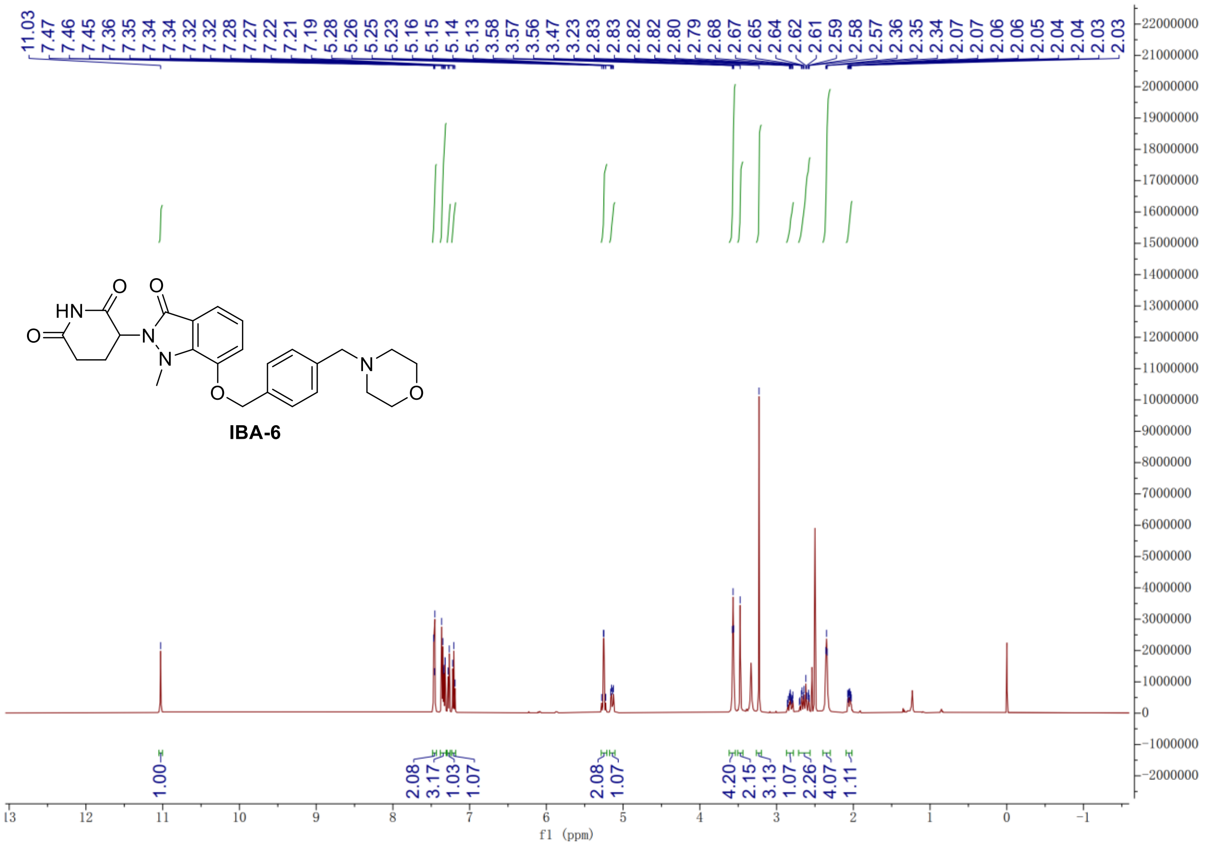

### $^{13}$C NMR Spectrum of Compound **IBA-6**

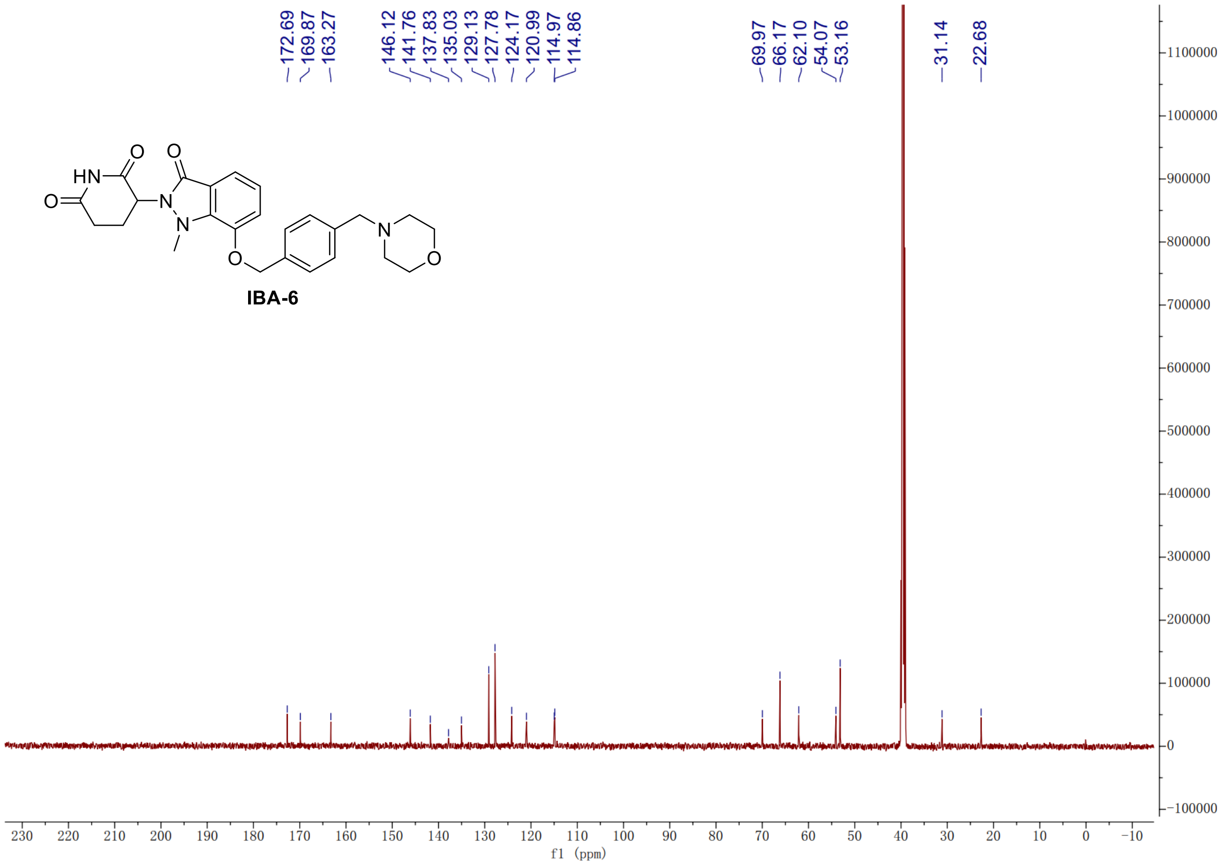

#### $^{1}$H NMR Spectrum of Compound **IBA-7**

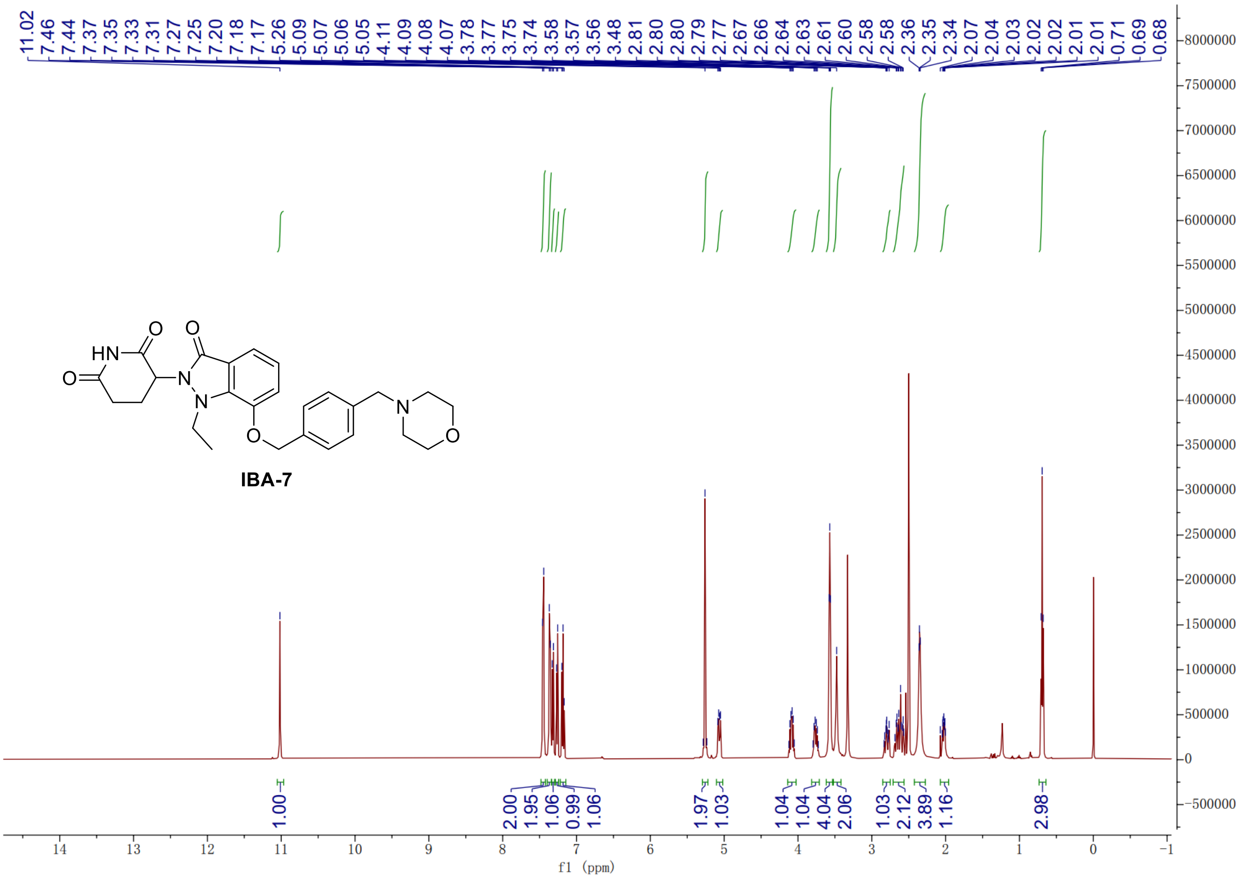

#### $^{13}$C NMR Spectrum of Compound **IBA-7**

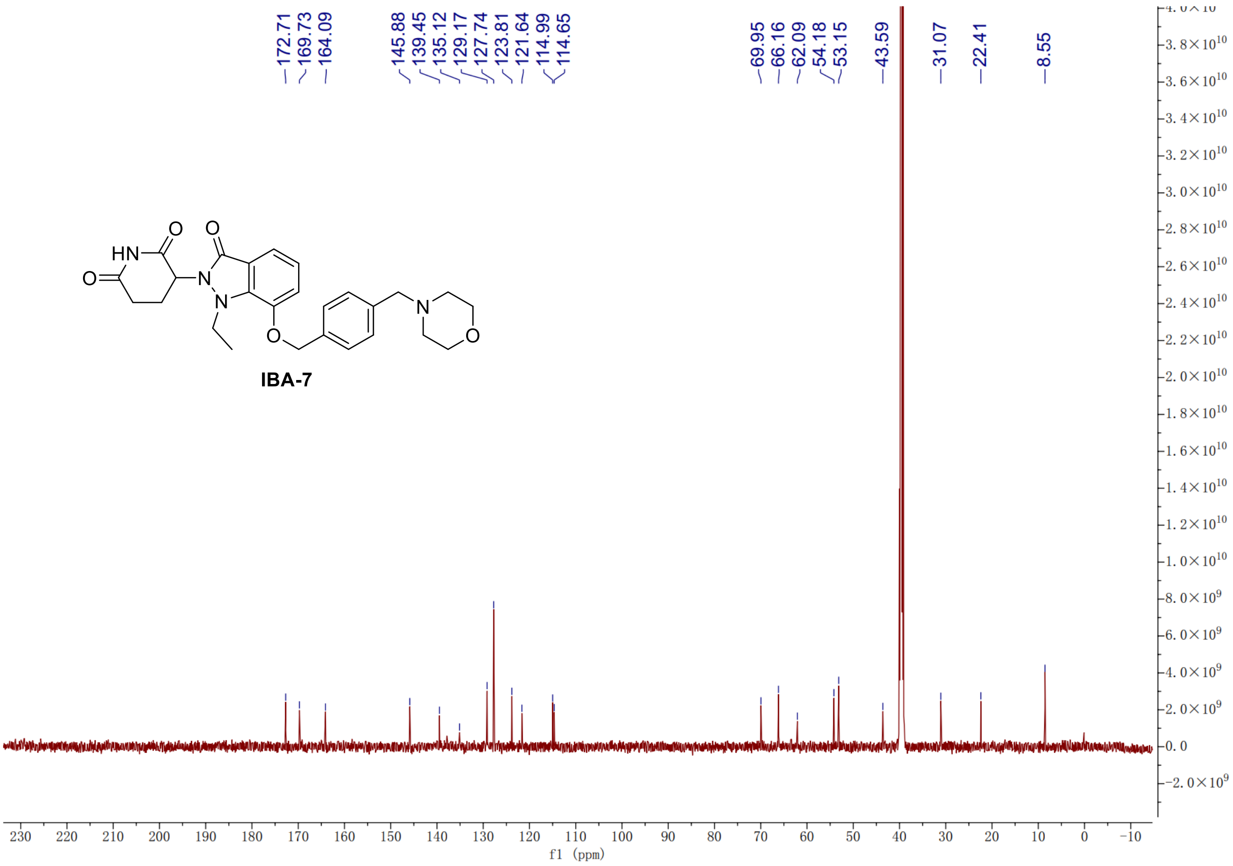

### [1]H NMR Spectrum of Compound **IBA-8**

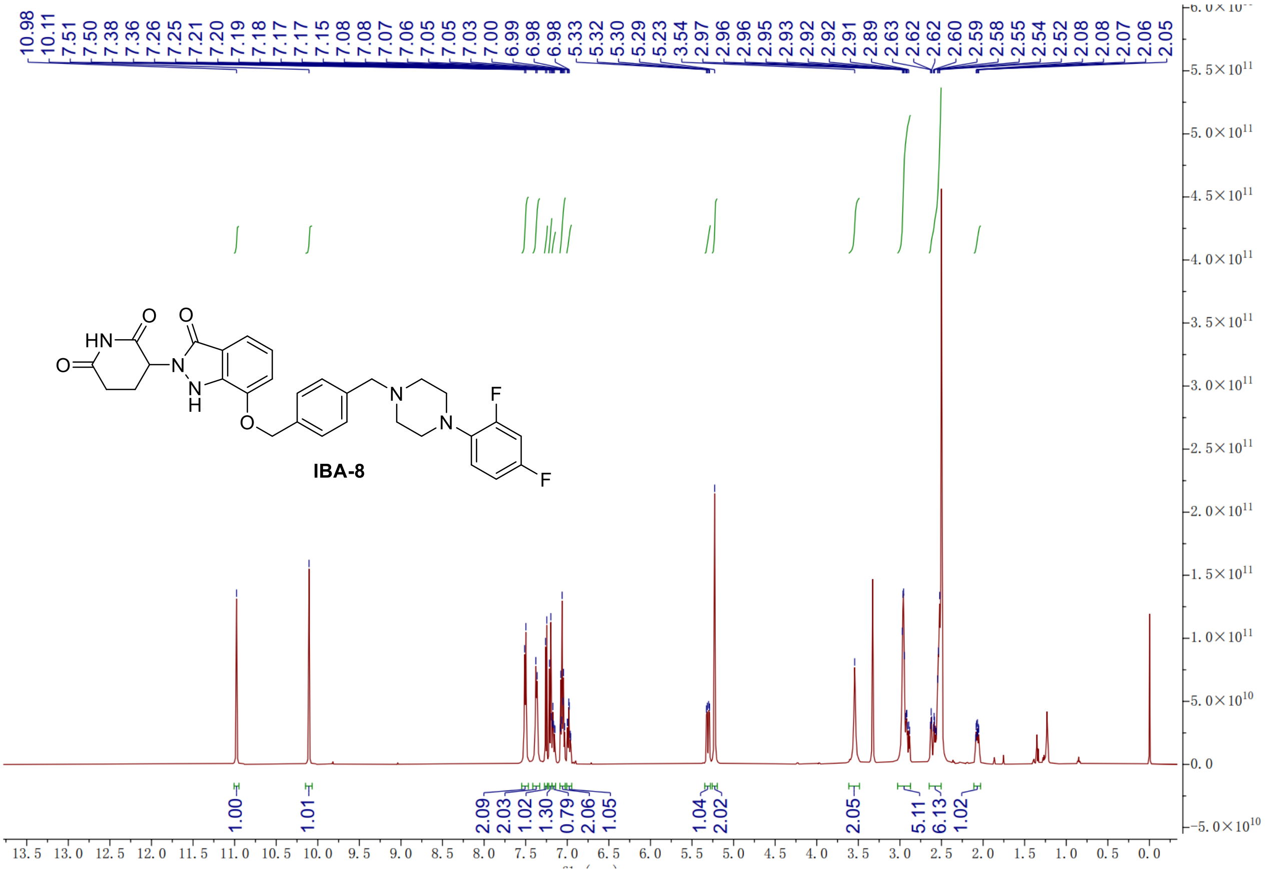

### [13]C NMR Spectrum of Compound **IBA-8**

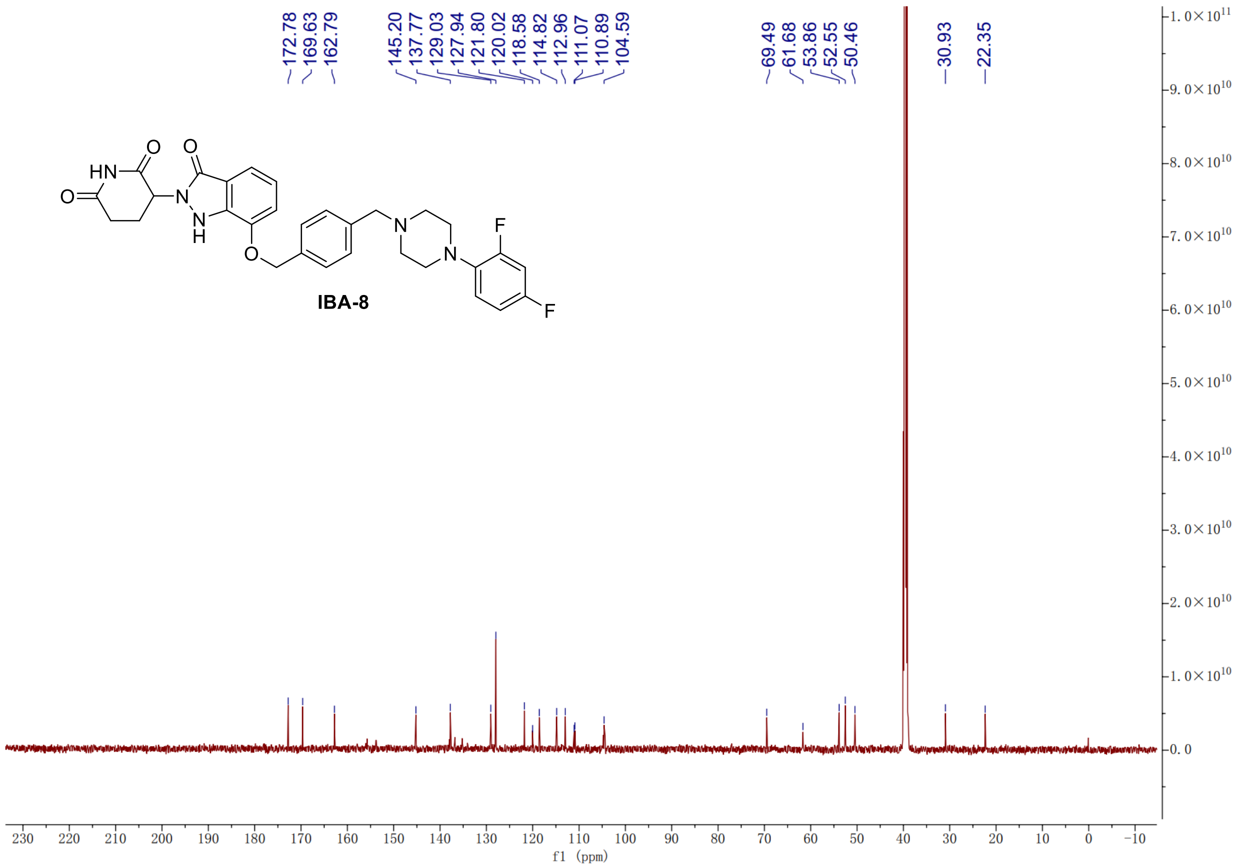

#### [1]H NMR Spectrum of Compound **IBA-9**

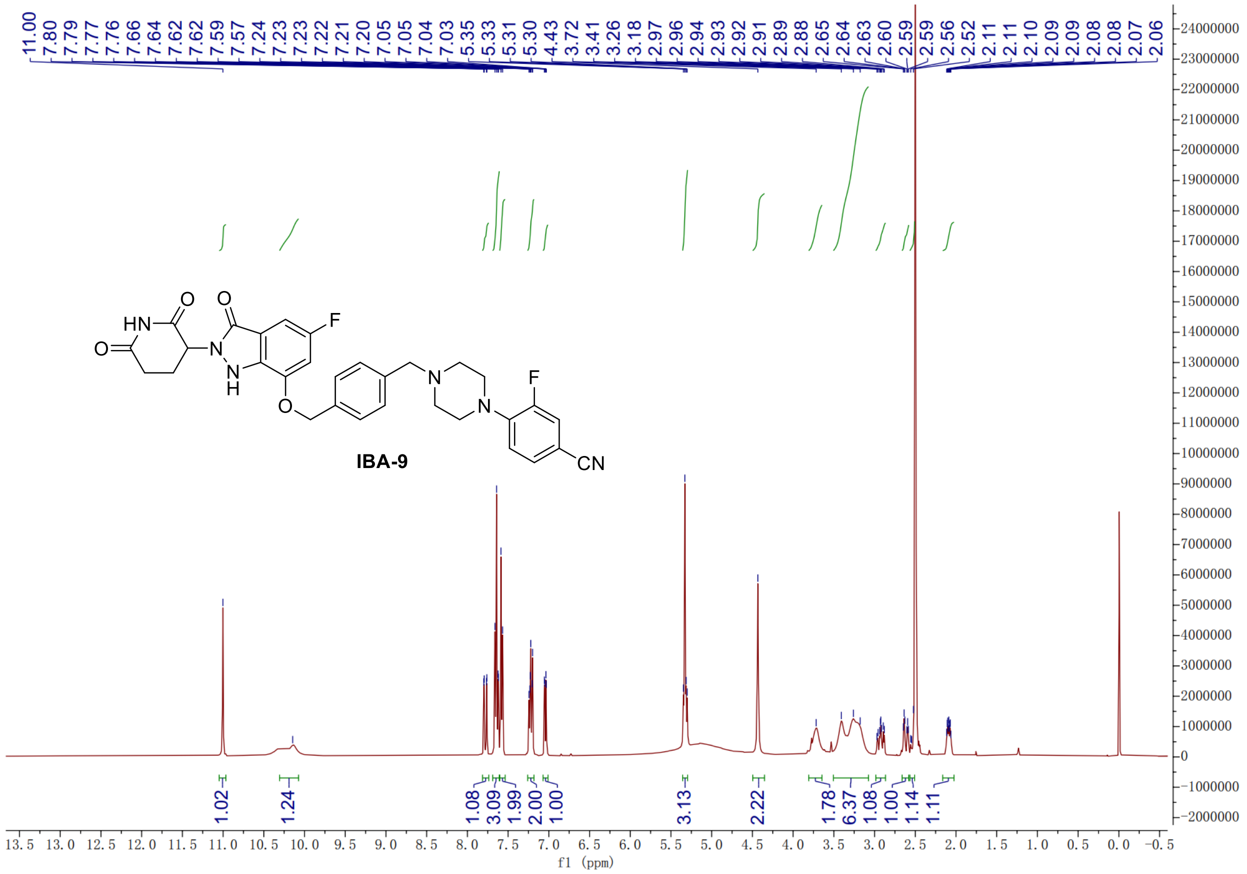

#### [13]C NMR Spectrum of Compound **IBA-9**

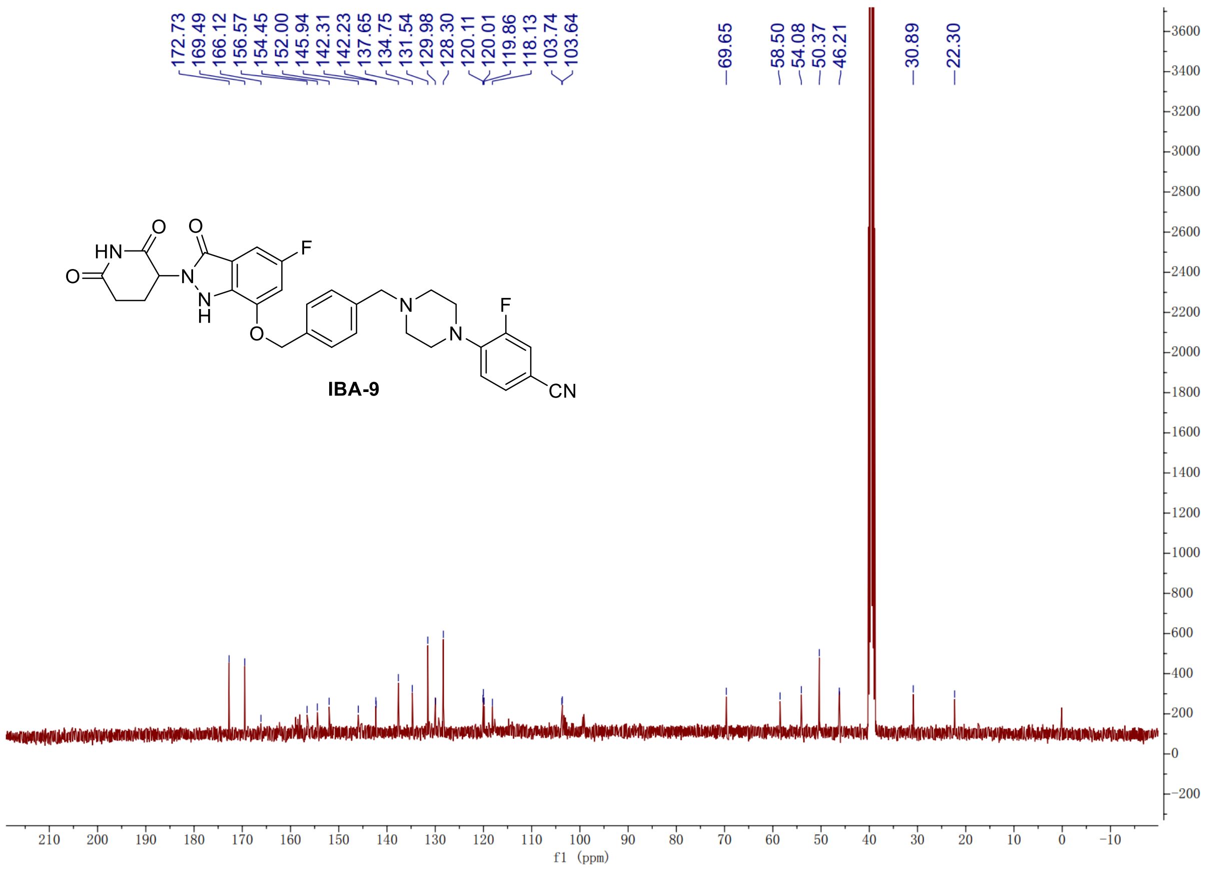

#### [1]H NMR Spectrum of Compound **IBA-10**

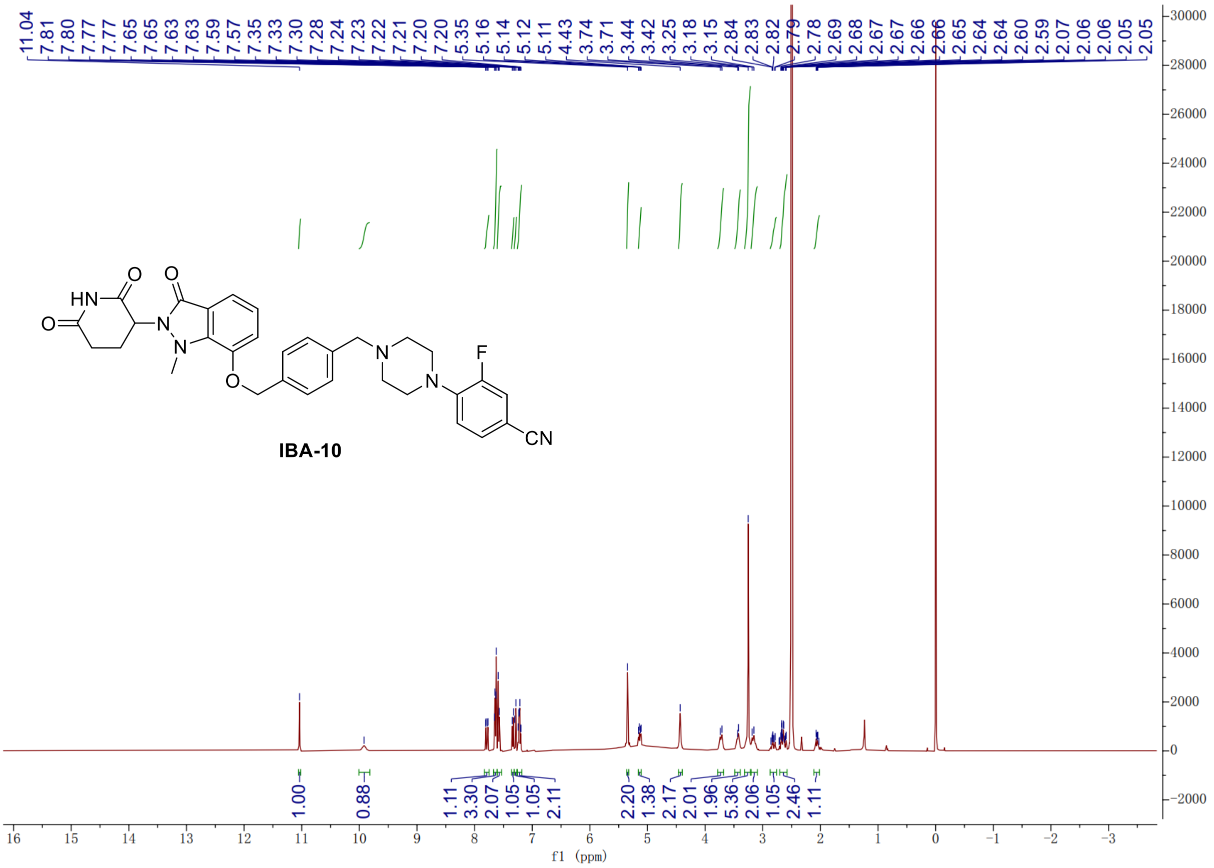

#### [13]C NMR Spectrum of Compound **IBA-10**

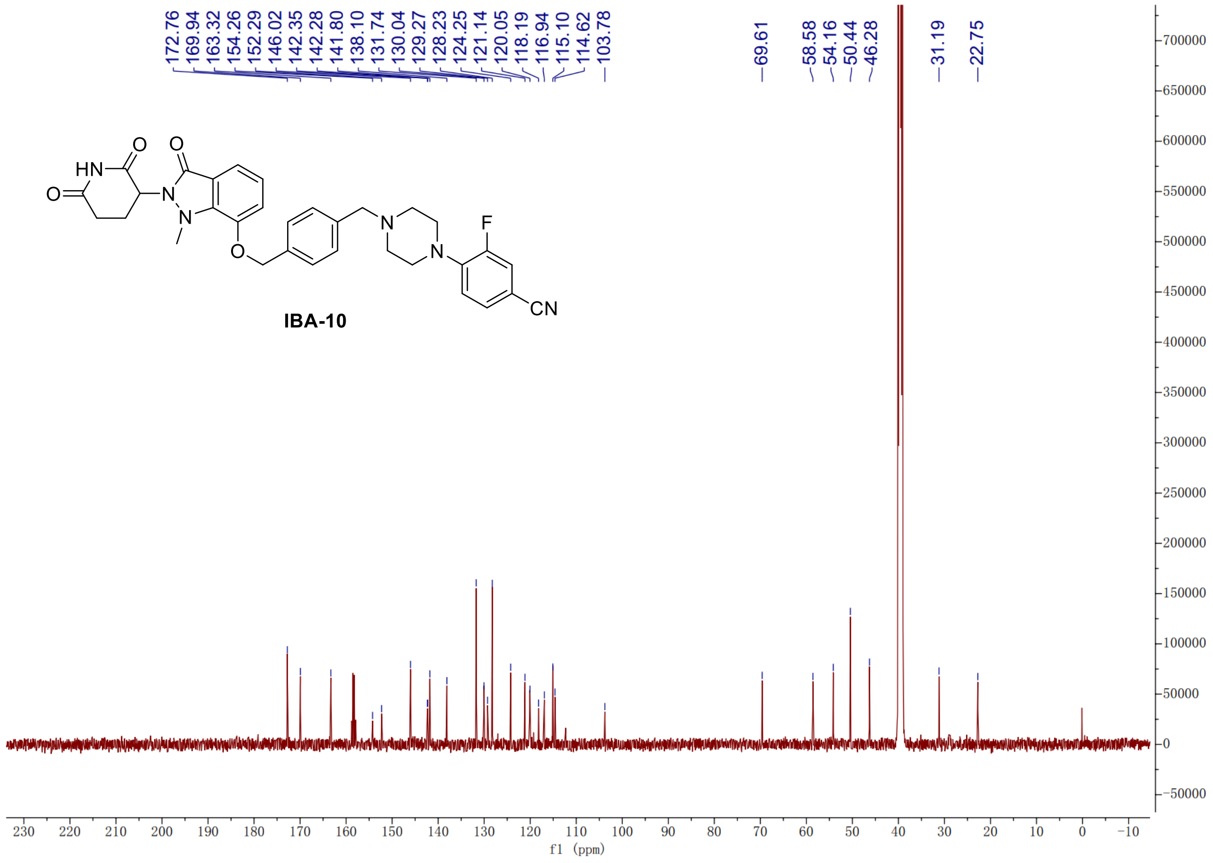

#### [1]H NMR Spectrum of Compound **IBA-11**

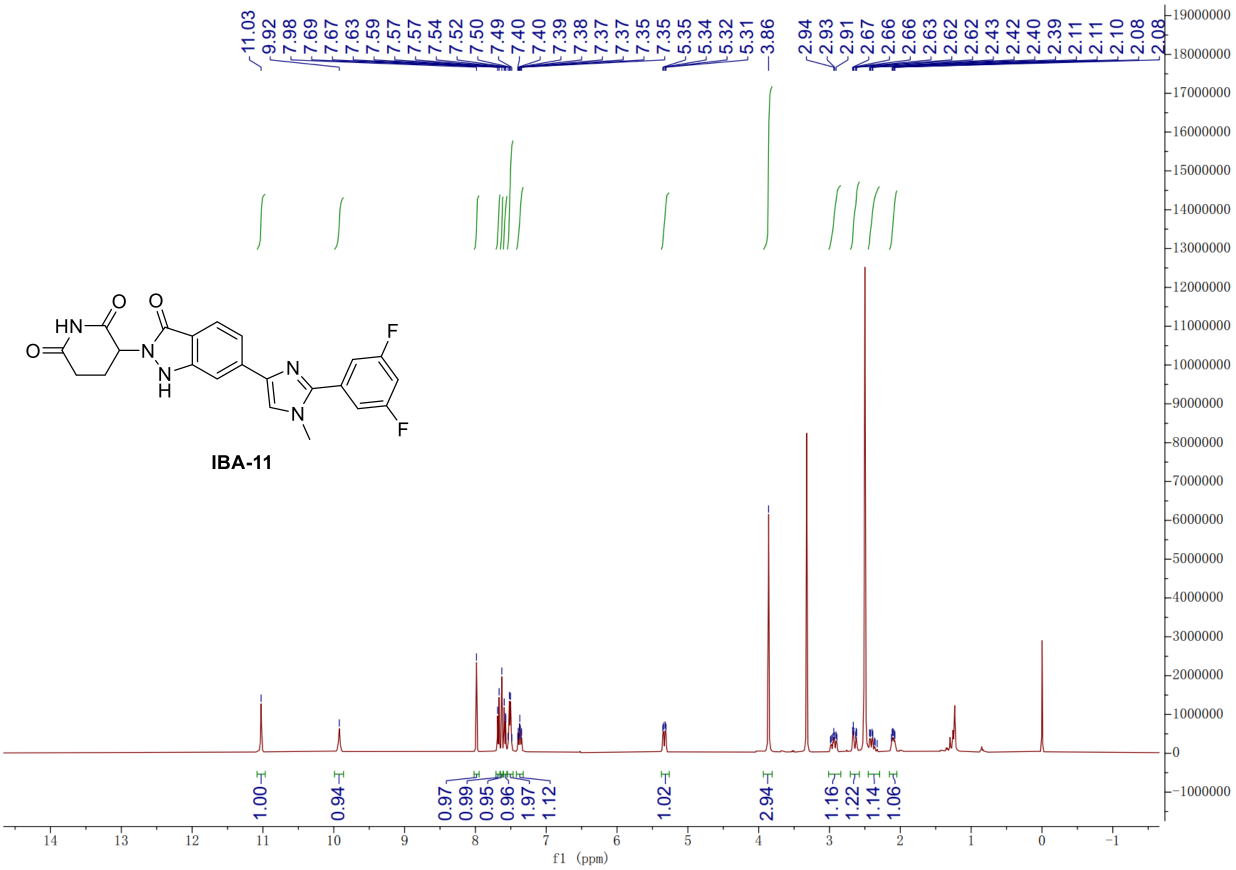

#### [13]C NMR Spectrum of Compound **IBA-11**

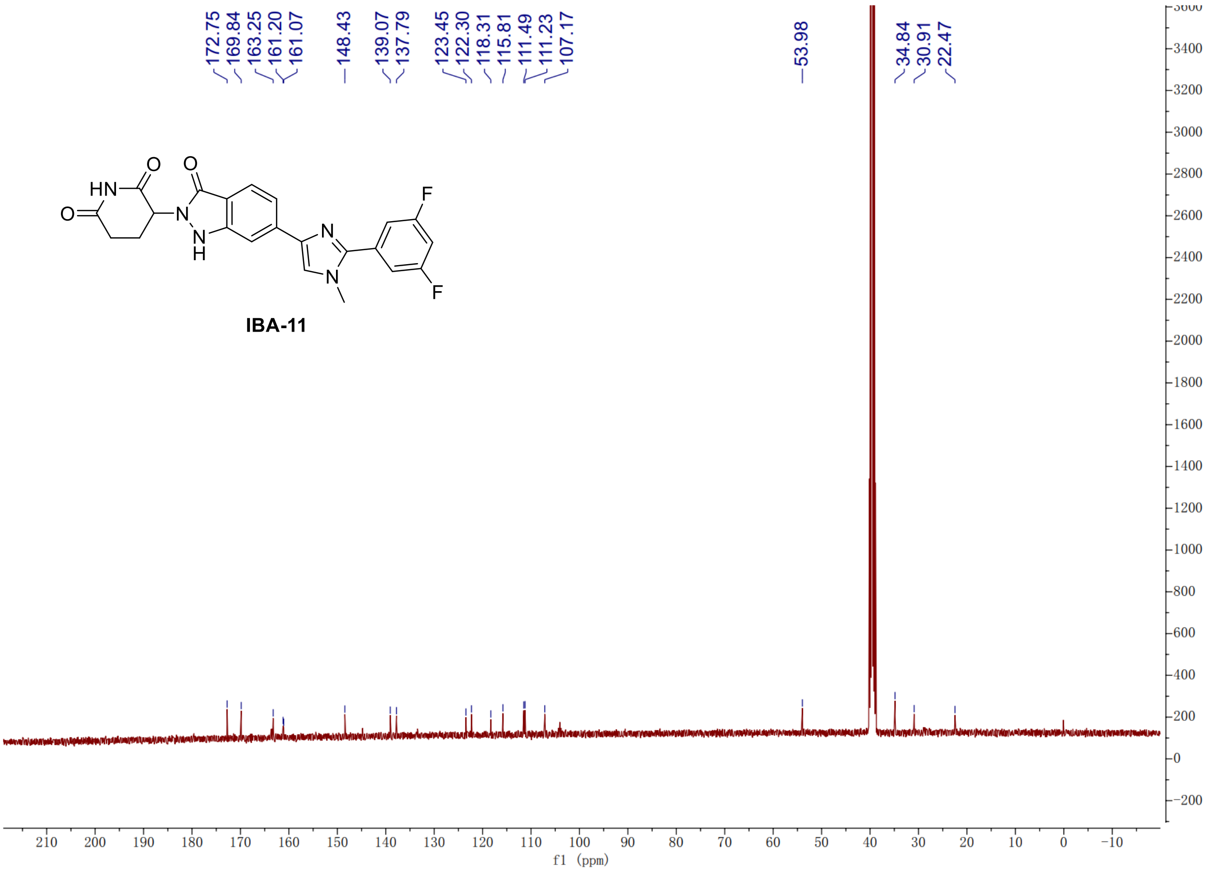

[1]H NMR Spectrum of Compound **IBA-12**

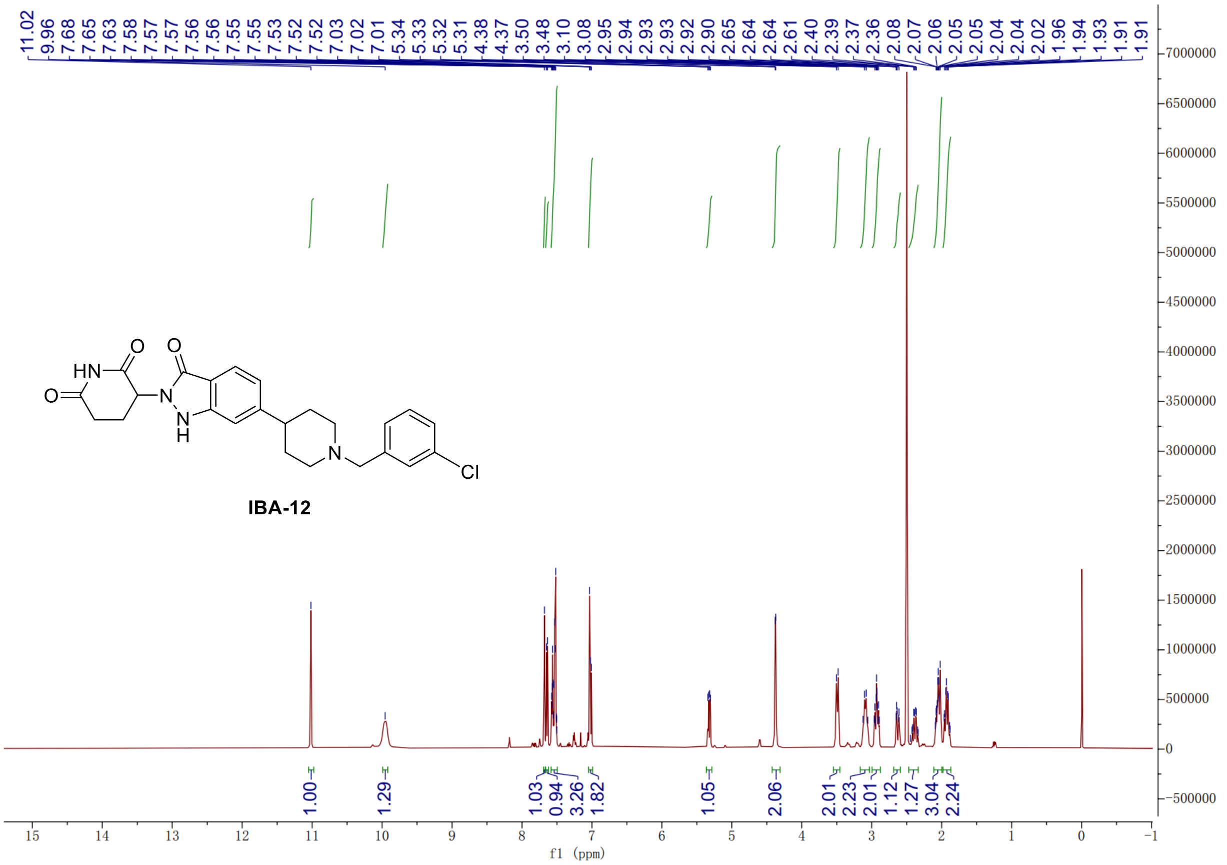

[13]C NMR Spectrum of Compound **IBA-12**

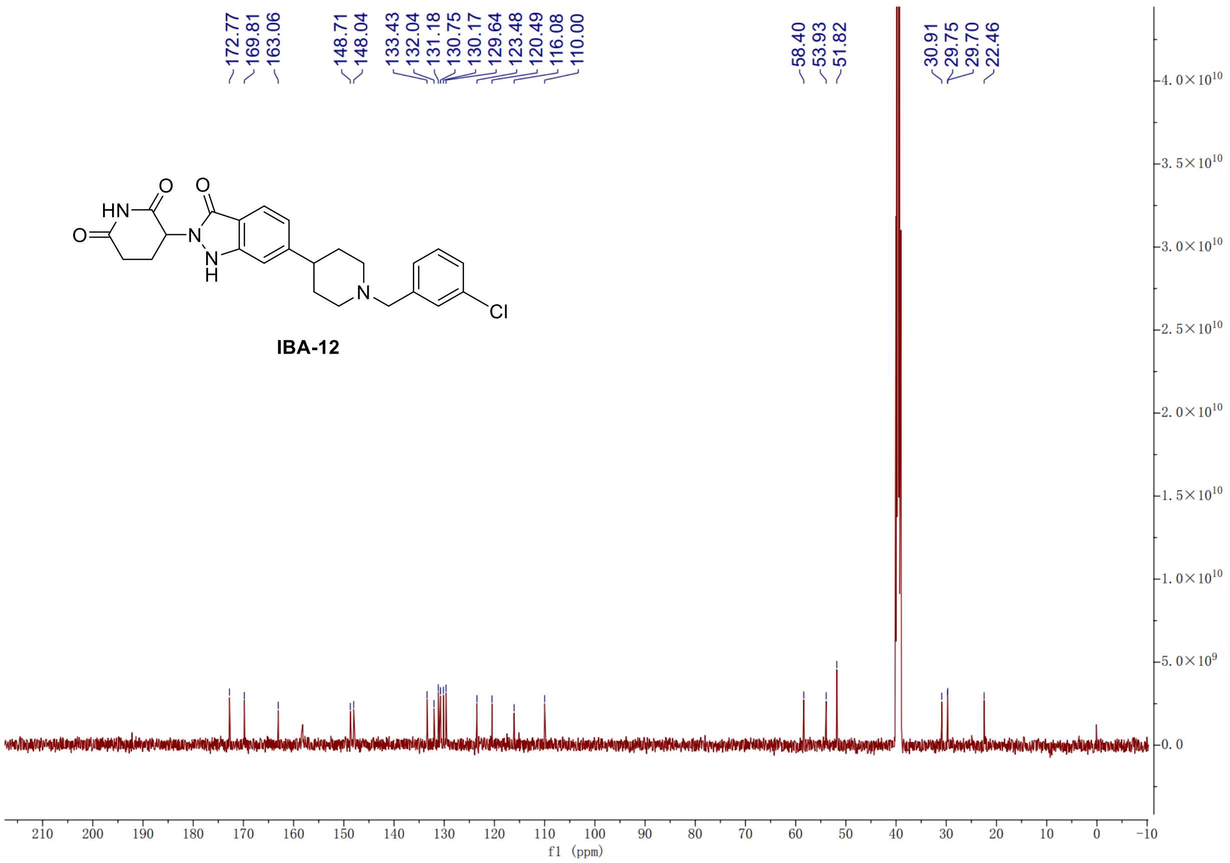
